# Huygens synchronization of medial septal pacemaker neurons generates hippocampal theta oscillation

**DOI:** 10.1101/2021.01.22.427736

**Authors:** Barnabás Kocsis, Sergio Martínez-Bellver, Richárd Fiáth, Andor Domonkos, Katalin Sviatkó, Péter Barthó, Tamás F. Freund, István Ulbert, Szabolcs Káli, Viktor Varga, Balázs Hangya

## Abstract

Episodic learning and memory retrieval are critically dependent on a hippocampal 4-12 Hz oscillatory ‘clock’ signal, the theta oscillation. This clock is largely externally paced, by a network of GABAergic neurons in the medial septum (MS). Theoretical studies suggested a range of hypotheses how this network may achieve theta synchrony; however, experimental evidence is still lacking. By recording multiple single MS neurons and hippocampal local field potential oscillations simultaneously, with both acute and chronically implanted silicon probes, we show that MS pacemaker units oscillate at individual frequencies within the theta range in rodents. Synchronization of MS neuron frequencies, accompanied by an elevation of firing rates, was found to parallel hippocampal theta formation in multiple rodent model systems. This suggests a general mechanism for theta synchronization, akin to the synchronization of weakly coupled pendulum clocks observed by Huygens in the 17^th^ century. We optogenetically identified the MS pacemaker units as parvalbumin-expressing GABAergic neurons, while the previously enigmatic MS glutamatergic neurons were mostly theta-activated non-rhythmic cells. Our data were consistent with a network model of partially connected single-compartment inhibitory pacemaker neurons, in which synchronization and de-synchronization in the frequency domain upon waxing and waning tonic excitatory drive was sufficient to toggle MS network output between theta and non-theta states. These results provide experimental and theoretical support to a frequency-synchronization mechanism for pacing hippocampal theta, which may serve as an inspirational prototype for the countless examples of synchronization processes in the central nervous system from Nematoda to Anthropoda to Chordate and Vertebrate phyla.

## Introduction

Exploratory behaviors are accompanied by a 4-12 Hz theta oscillation in the hippocampus of rodents, carnivores and primates, including humans (Buzsáki, 2002; Buzsáki and Moser, 2013; Kahana et al., 1999; Rutishauser et al., 2010). This theta oscillation provides a temporal reference signal for hippocampal networks, which is crucial for episodic learning and memory retrieval (Buzsáki, 2006). GABAergic neurons of the MS have been identified as pacemaker neurons of hippocampal theta rhythms. First, the MS and the hippocampus are reciprocally interconnected by septo-hippocampal GABAergic, cholinergic and glutamatergic, as well as hippocampo-septal GABAergic connections (Freund and Antal, 1988; Huh et al., 2010; Toth et al., 1993; Zaborszky et al., 2012). Second, MS neurons show strong phase coupling to hippocampal theta, with a subset of neurons remaining theta rhythmic in the absence of theta oscillation in the hippocampus (Barrenechea et al., 1995; Borhegyi et al., 2004; Ford et al., 1989; Petsche et al., 1962; Stewart and Fox, 1989; Varga et al., 2008). Third, lesioning or blocking all, or selectively the GABAergic septo-hippocampal projections abolishes hippocampal theta (Brazhnik and Vinogradova; Teitelbaum et al., 1975; Yoder and Pang, 2005). Fourth, it is possible to induce hippocampal theta oscillation by stimulating the medial septum electrically, or MS GABAergic neurons optogenetically (Green and Arduini, 1954; Zutshi et al., 2018). Fifth, GABAergic neurons of the MS functionally lead the hippocampal network during theta activity, as their activity changes precede correlated changes in both hippocampal interneuron firing and network activity (Hangya et al., 2009).

Despite the long-recognized importance of hippocampal theta oscillation, the mode of theta-frequency synchronization within the septal GABAergic network has not been tested experimentally. Based on theoretical work, it was proposed that individual pacemaker neurons may oscillate at different frequencies in non-theta states and synchronize in the frequency domain upon increased depolarizing input, giving rise to hippocampal theta rhythm (Ujfalussy and Kiss, 2006; Wang, 2002). This mechanism was termed ‘Huygens synchronization’ in the physics of coupled oscillators, and was hypothesized that it may also apply to neuronal synchrony (Oliveira and Melo, 2015; Ramirez et al., 2016). Two influential models assumed the presence of two separate GABAergic populations that inhibit each other through a so-called ‘ping pong’ mechanism, either within the MS network (Cutsuridis and Poirazi, 2015; Ujfalussy and Kiss, 2006) or involving the hippocampo-septal GABAergic projection neurons (Wang, 2002). Theta amplitude modulation through increased MS firing rates was proposed as a direct layer of control (Bland et al., 1996; Denham and Borisyuk, 2000; Tokuda et al., 2019), whereas a change in MS bursting patterns could also impact theta generation (Joshi et al., 2017; Simon et al., 2006; Sotty et al., 2003). The role of the hyperpolarization-activated and cyclic nucleotide-gated (HCN) channels was emphasized as part of the potential underlying ionic mechanisms (Kocsis and Li, 2004; Varga et al., 2008), whereas other studies investigated the effects of direct modulation of potassium or chloride conductance either in the MS or in the hippocampus (Bland et al., 1996; Hajós et al., 2004; Zou et al., 2011). However, beside frequency synchronization, other mechanisms are also plausible including synchronization in the phase domain without changes in oscillation frequency (Borhegyi et al., 2004; Ujfalussy and Kiss, 2006). Additionally, while the identification of PV-expressing GABAergic neurons as pacemakers is widely accepted, little is known about the exact role of other medial septal cell types in theta generation (Lee et al., 1994; Robinson et al., 2016; Vandecasteele et al., 2014).

Therefore, we investigated potential theta generation mechanisms experimentally by recording single neurons from the medial septum, with concurrent local field potentials (LFP) from the hippocampus using multichannel silicon electrode arrays. To address the generality of results across theta oscillations observed in awake or anesthetized rodents (Kramis et al., 1975; Stewart and Fox, 1989), we obtained three data sets: from urethane anesthetized rats, urethane anesthetized mice and awake, drug-free mice. We used optogenetic tagging (Kvitsiani et al., 2013; Lima et al., 2009) in anesthetized mice to unveil the genetic identity of the key players of theta formation. We found that hippocampal theta onset was accompanied by Huygens (frequency) synchronization of MS pacemaker neurons otherwise rhythmic at more distinct frequencies in the theta range. This mechanism held across the three rodent theta models tested, arguing for a general synchronization mechanism despite the known behavioral and pharmacological heterogeneity of hippocampal theta rhythms (Buzsáki, 2006; Kramis et al., 1975). While we could confirm the pacemaker role of PV-expressing GABAergic neurons (Freund and Antal, 1988; Hangya et al., 2009; Varga et al., 2008), glutamatergic neurons showed strong firing rate increase during theta without much rhythmicity, suggesting they may contribute to theta-associated increase of tonic drive to the pacemakers (Ford et al., 1989; Green and Arduini, 1954; Oddie et al., 1996; Vandecasteele et al., 2014; Yang et al., 2014). These results were consistent with a circuit model of the medial septum in which a homogeneous network of inhibitory neurons was capable of synchronizing and desynchronizing in the frequency domain upon changing levels of tonic excitatory input.

## Results

### Recording medial septal and hippocampal data in three rodent models of theta oscillations

We set out to investigate how the medial septal network generates hippocampal theta oscillations. To achieve this, it is necessary to co-register the activity of the MS network with hippocampal local field oscillations (Figure 1). Hippocampal theta oscillations have traditionally been investigated in both urethane anesthetized and awake rodents (Borhegyi et al., 2004; Buzsáki, 2002; Klausberger and Somogyi, 2008; Kramis et al., 1975; Mikulovic et al., 2018; Stewart and Fox, 1989). To explore the commonalities and differences across different theta model systems, we collected three independent data sets. We recorded multiple single neurons from the MS using silicon probes from anesthetized rats (n = 903 neurons from N = 7 rats; 416 and 636 neurons active during both theta and non-theta states or at least one of them, respectively; Figure S1A-B), anesthetized mice (n = 1155 neurons from N = 11 mice; 617 and 950 active during both or at least one of theta and non-theta states; Figure 1A-D) and awake mice (n = 312 neurons from N = 4 mice; 265 and 289 active during both or at least one of the states; Figure S1C-D). Simultaneously, we recorded local field potentials (LFP) from all layers of the dorsal CA1 area of the hippocampus using linear silicon probe electrodes (Figure 1E; Figure S2A-B). We detected the hippocampal layers based on combined information from histology and the phase reversal of the hippocampal theta oscillation (Buzsáki, 2006; Buzsaki et al., 1986; Green et al., 1960) (Figure S2C). We used the LFP from stratum radiatum / lacunosum-moleculare to precisely detect theta epochs, since theta oscillation had the largest amplitude in those layers (Buzsáki, 2002). Theta frequency boundaries were defined specifically for the three types of recordings based on LFP spectra, confirming the known observation of slower oscillations under urethane anesthesia (Borhegyi et al., 2004; Varga et al., 2008) (Figure 1F). Theta detection was based on the theta/delta spectral ratio, as commonly applied (McNamara et al., 2014; Wikenheiser and Redish, 2013) (Figure 1G-H).

**Figure 1.**
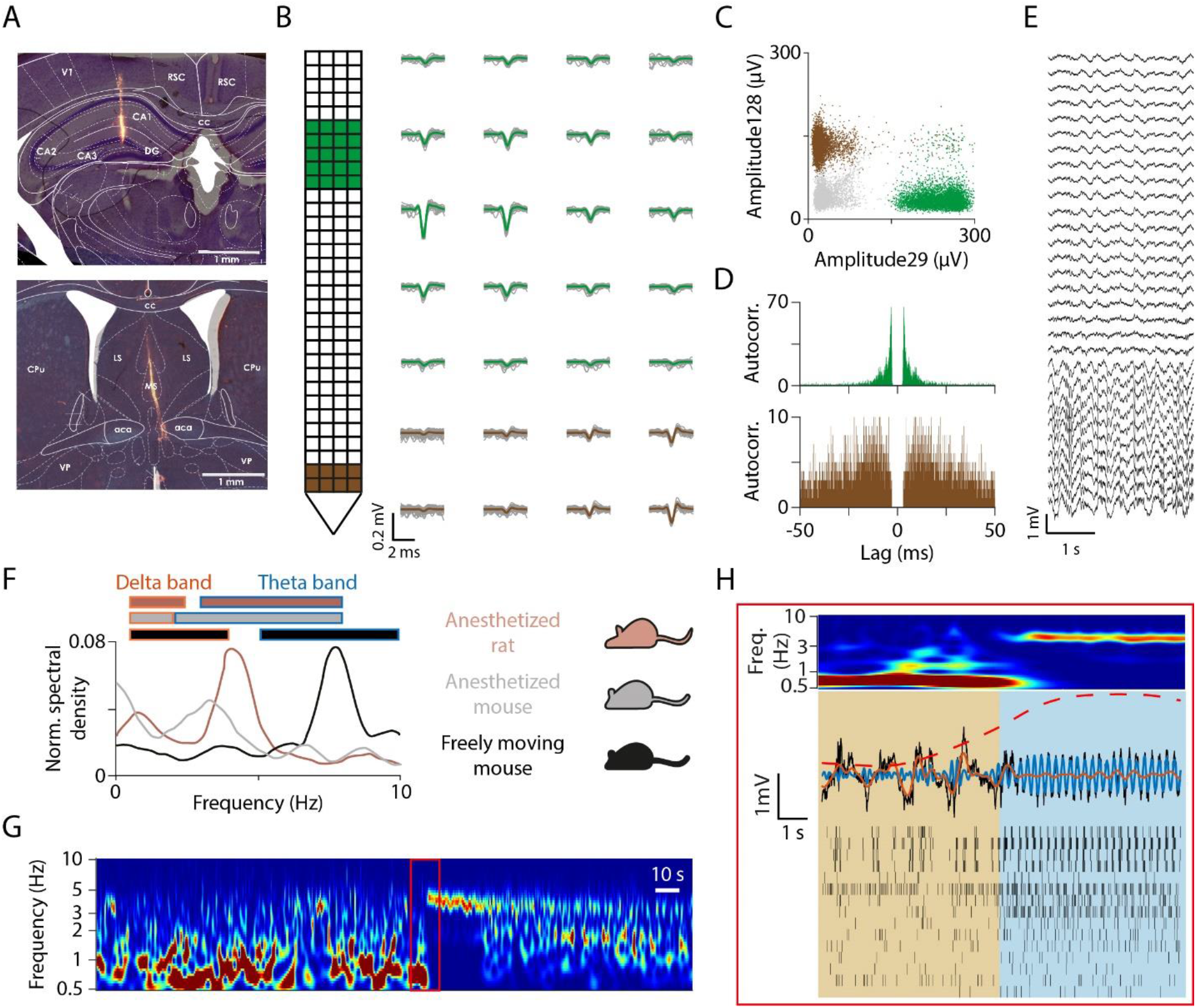
Dual recording of hippocampal local field potentials and single neuron activity from the medial septum. (A) Histology of the silicon probe tracks in the mouse hippocampus (top) and medial septum (bottom). Nissl-staining; red, DiI. See also Figure S1 for histology in rats. (B) Medial septal single units were collected using a 128-channel silicon probe in anesthetized mice (see Methods for specifics on different probes used in our experiments). Example spike waveforms for two neurons are shown for the channels marked on the schematic representation of the probe (green and brown, average waveforms; grey, 100 randomly selected individual spike). (C) Amplitudes (peak-to-valley) on selected channels show clear separation of the example cells. (D) Autocorrelograms of the example neurons. (E) Hippocampal LFP was collected concurrently using a linear silicon probe. A 3-second-long example recording is shown (two broken channels were removed). (F) We collected data from three rodent models of theta oscillations: urethane-anesthetized rats (brown), urethane-anesthetized mice (grey) and awake mice (black). Representative Fourier spectra of hippocampal LFP recordings from the three model systems. Theta and delta band boundaries were determined based on Fourier spectra of hippocampal LFPs for each model separately, indicated by color coded horizontal bars on top. (G) Representative hippocampal LFP wavelet spectrogram from an anesthetized rat recording. (H) Top, theta onset is enlarged from panel G. Middle, raw LFP (black), LFP filtered in the theta (blue) and delta frequency band (orange) and theta-delta power ratio (dashed line). Bottom, raster plot of individual units (n = 15) recorded from the medial septum concurrently. Note the change in network activity at theta onset.

### Putative pacemaker neurons show constitutive theta-rhythmic activity

We developed an unbiased method to define functional cell classes of MS neurons based on their rhythmic firing properties. Theta Rhythmicity Index was defined based on the normalized autocorrelation peak in the theta frequency band (see Methods). Since we observed that some MS neurons showed slow rhythmic activity, more prominent in rats than in mice, we defined a Delta Rhythmicity Index similarly. Significant rhythmic activity was tested against bootstrap distributions generated from simulated Poisson-neurons that were matched with MS neurons in their firing rates (Figure S3).

We found that a subpopulation of MS neurons showed theta-rhythmic bursting activity both when theta oscillation was present and when it was absent in area CA1 of the hippocampus (Figure 2A). These neurons showed prominent theta oscillations in their autocorrelograms (Figure 2B-C), strong phase locking to CA1 theta oscillation but no strong phase relationship with hippocampal delta band activity during non-theta episodes (Figure 2D-G). We confirmed the presence of two separate clusters of neurons that showed anti-phase preference to hippocampal theta, as reported before (Borhegyi et al., 2004; Joshi et al., 2017; Varga et al., 2008). This constitutively theta-rhythmic population was found in anesthetized rats (n = 29/416), anesthetized mice (n = 47/617) and freely moving mice (n = 35/265; note that these numbers represent conservative estimates based on significant rhythmicity; Figure S4, Figure S5). They are commonly assumed to be the putative theta pacemaker neurons (Borhegyi et al., 2004; Ford et al., 1989; Hangya et al., 2009; Stewart and Fox, 1989; Varga et al., 2008).

**Figure 2.**
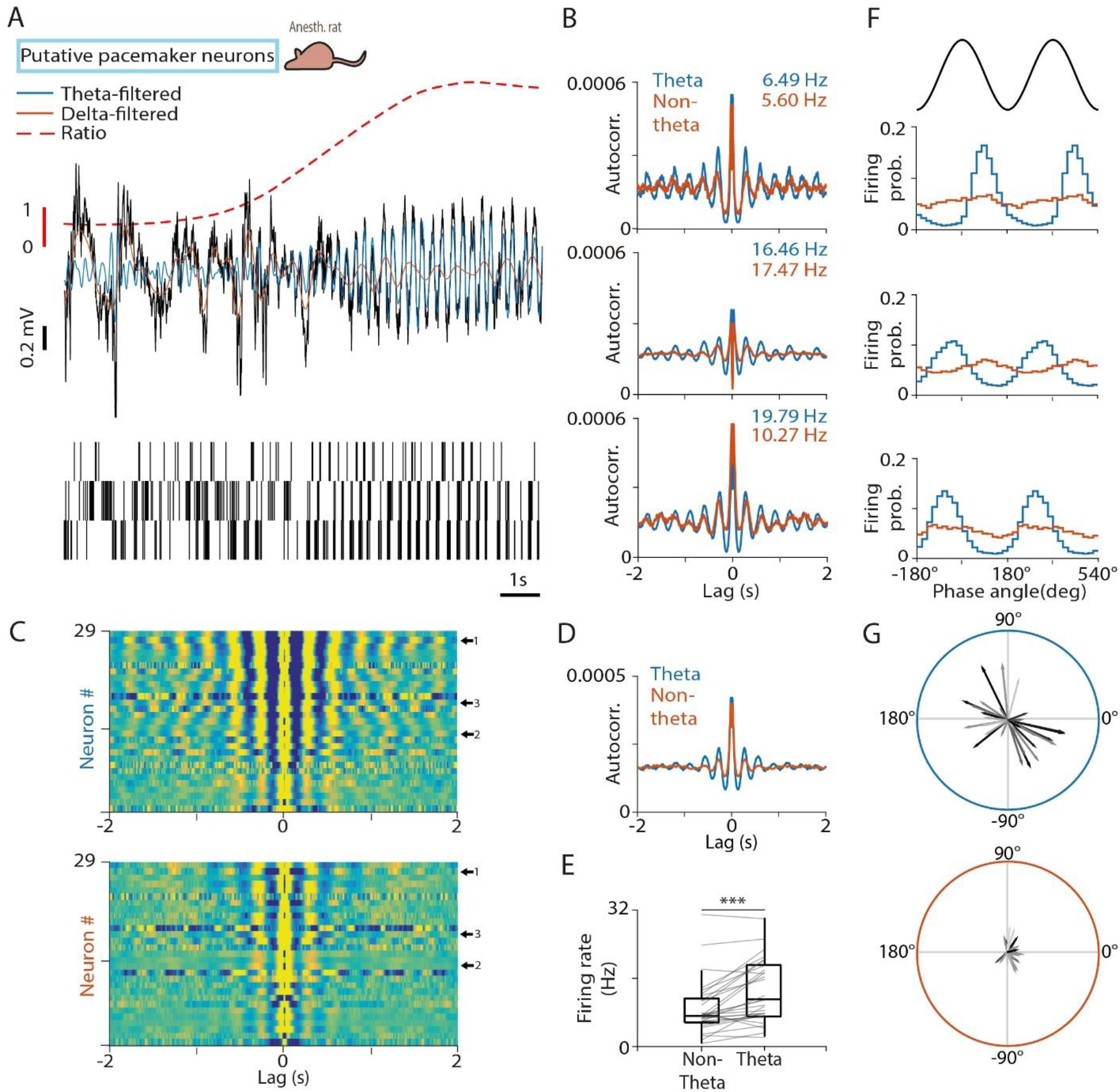
Putative pacemaker neurons of the MS. (A) Top, black, raw LFP from the CA1 shows a delta-to-theta state transition. Orange, LFP filtered in the delta band; blue, LFP filtered in the theta band; dashed, theta-delta amplitude ratio. Bottom, spike raster of three examples of putative pacemaker neurons from the same recording session. Recordings are from urethane-anesthetized rats; see Figure S4 and Figure S5 for the anesthetized and awake mouse recordings. (B) Autocorrelograms of the three example neurons in panel (A). Numbers indicate average firing rates during theta and non-theta segments. (C) Autocorrelograms of all putative pacemaker neurons during theta (top) and non-theta (bottom) segments. Arrows indicate the example neurons. (D) Average autocorrelogram of all putative pacemaker neurons. (E) Median firing rate of putative pacemaker neurons during non-theta and theta segments. Boxes and whiskers represent interquartile ranges and non-outlier ranges, respectively. Lines correspond to individual neurons. Firing rate was significantly higher during theta. ***, p < 0.001, Wilcoxon signed-rank test. (F) Phase histogram of the example neurons in panel (A) relative to delta (orange) and theta (blue) oscillations. Two oscillatory cycles are shown. (G) Phase-locking of all putative pacemaker neurons to theta (top) and delta (bottom) oscillation in polar coordinates. Angle, circular phase; length, mean vector length; greyscale corresponds to average firing rate (black, higher firing rate). Note that neurons formed two opposite phase clusters during theta, whereas phase-locking strength to delta oscillations was low for putative pacemaker neurons.

### Putative ‘follower’, theta-skipping and tonically active MS neurons

A considerable MS population ‘followed’ the ongoing activity of the septo-hippocampal network. These neurons were theta-rhythmic during hippocampal theta oscillation but not when hippocampal theta was absent, often phase locked to both theta and delta band oscillations. In anesthetized rats, many of these neurons showed slow rhythmic activity at delta band frequencies in the absence of hippocampal theta (Figure 3; Figure S6), while in mice, these putative ‘follower’ neurons were mostly non-rhythmic or slow firing in non-theta states (‘theta follower neurons’, Figure S7, Figure S8, Figure S9). Some neurons exhibited ‘follower’ activity only in the non-theta state, remaining non-rhythmic or silent during hippocampal theta.

**Figure 3.**
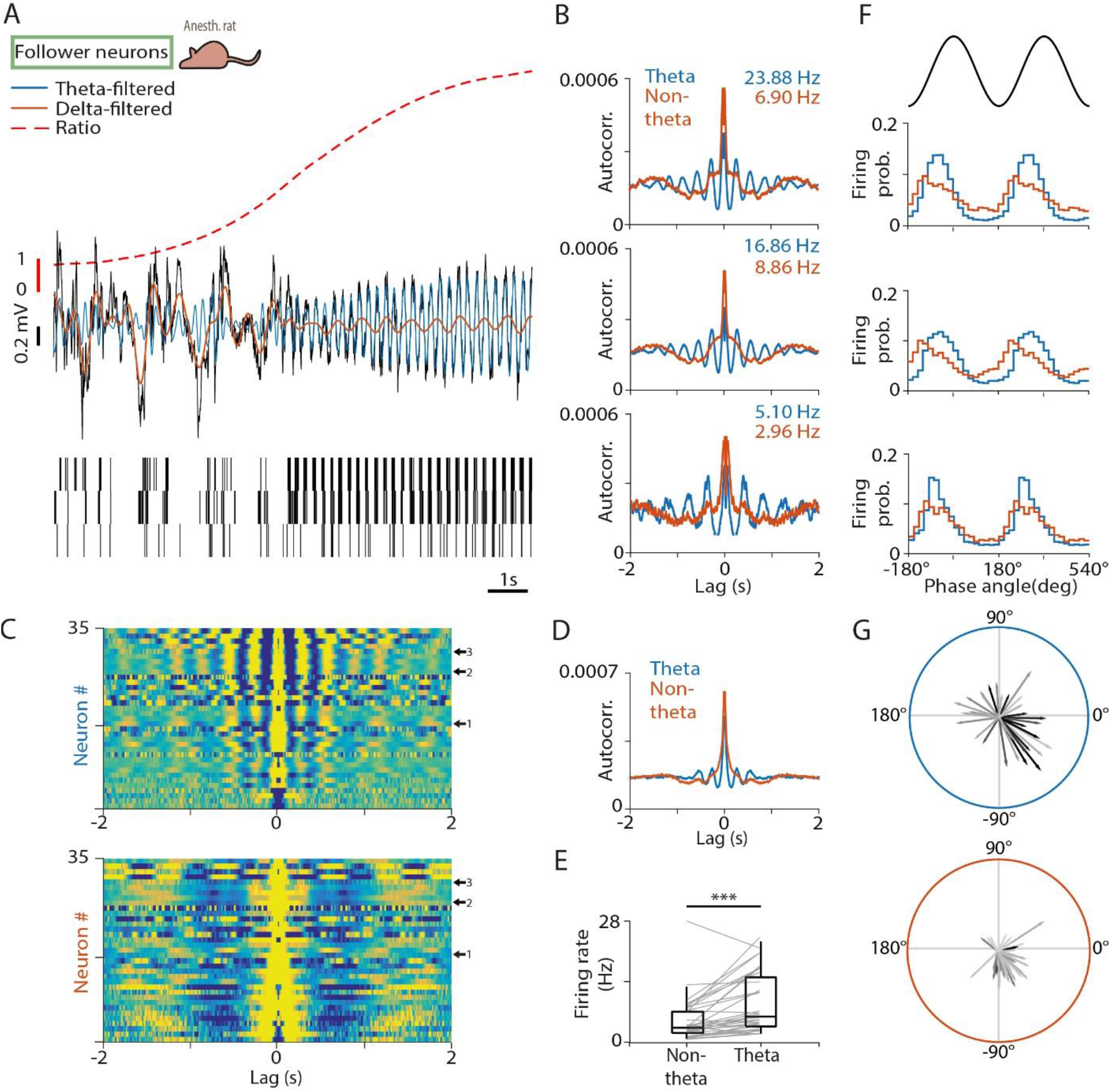
Follower MS neurons. (A) Top, black, raw LFP from the CA1 shows a delta-to-theta state transition. Orange, LFP filtered in the delta band; blue, LFP filtered in the theta band; dashed, theta-delta amplitude ratio. Bottom, spike raster of three example neurons from the same recording session. ‘Follower’ neurons showed delta-rhythmic activity during non-theta epochs and theta-rhythmic activity during theta epochs in urethane-anesthetized rats. See Figure S7 for mouse recordings. (B) Autocorrelograms of the three example neurons in panel (A). Numbers indicate average firing rates during theta and non-theta segments. (C) Autocorrelograms of all follower neurons during theta (top) and non-theta (bottom) segments. Arrows indicate the example neurons. (D) Average autocorrelogram of all follower neurons. (E) Median firing rate of follower neurons during non-theta and theta segments. Boxes and whiskers represent interquartile ranges and non-outlier ranges, respectively. Lines correspond to individual neurons. Firing rate was significantly higher during theta. ***, p < 0.001, Wilcoxon signed-rank test. (F) Phase histogram of the example neurons in panel (A) relative to delta (orange) and theta (blue) oscillations. Two oscillatory cycles are shown. (G) Phase-locking of all follower neurons to theta (top) and delta (bottom) oscillation in polar coordinates. Angle, circular phase; length, mean vector length; greyscale corresponds to average firing rate (black, higher firing rate). Many of these neurons were phase-locked to both hippocampal oscillations.

A subpopulation of MS neurons, identified previously as ‘theta-skipping’, showed slow rhythmic activity regardless whether theta oscillation was present in CA1 (Figure S10). While some of these neurons doubled their rhythmic frequency at theta onset (but still in the delta range), many cells of this population maintained a constant slow rhythmicity, often firing regularly in every second theta cycle. Interestingly, while this group was rather prominent in anesthetized rats, these cells were only rarely found in mice (Figure S11), revealing species-specific differences in the MS theta generation network (anesthetized rats, 48/416; anesthetized mice, 4/617; freely moving mice, 1/265).

Finally, a group of MS cells showed regular rhythmic firing patterns resembling those of striatal cholinergic interneurons (Inokawa et al., 2010); therefore, we called this novel MS group ‘tonically active neurons’ (Figure S12A). Although these neurons fired rhythmically with a frequency in the theta band (Figure S12B-C), they showed little to no overall phase locking to hippocampal activity (Figure S12D-G), typically exhibiting rhythmicity at slightly higher frequencies than ongoing CA1 theta oscillation (based on autocorrelation peak, median ± s.e. of median, 5.68 ± 0.34, 4.29 ± 0.41 and 8.40 ± 0.37 Hz during theta segments in anesthetized rats, anesthetized mice and freely moving mice, respectively). They did not show the long bursts with 20-50 ms interspike intervals (ISIs) known to be characteristic of the pacemaker group, but some of them occasionally fired short, fast bursts of action potentials (ISI <15 ms; Figure S13). They were found in all three rodent theta models tested (anesthetized rats, 18/416; anesthetized mice, 49/617; freely moving mice, 8/265; Figure S14 and Figure S15). These neurons also showed a firing rate change during theta oscillation. Interestingly however, while their firing rate dropped at theta onset in rats (15/18 neurons), previously coined as a ‘theta-OFF’ pattern(Ford et al., 1989) (Figure S12E), they exhibited a theta-associated firing rate increase in mice, both anesthetized (34/49 neurons) and awake (8/8 neurons; Figure S14 and Figure S15).

Together with those cells that did not exhibit any rhythmic activity according to our conservative rhythmicity analysis, the above-described groups of MS neurons constitute the entire medial septal population (Figure S16; Supplemental Table 1).

### Septal pacemakers synchronize their burst frequencies

In order to gain information on the neural synchronies between the above-described groups, we calculated pairwise crosscorrelations for pairs of MS cells (n = 527 pairs in anesthetized rats). We observed strong correlations in the theta frequency band within and between putative pacemakers, putative followers and theta-skipping neurons (Figure S17). Surprisingly, almost no correlations were detected between the tonically active and other cell groups. Therefore, we reasoned that this group might not participate in the hippocampal theta generating mechanisms of the septum and their rhythmic firing serves other, yet unknown purposes.

The putative pacemaker neurons showed strong crosscorrelations with other pacemakers and rhythmic neurons that showed theta phase-locking. We found that during CA1 theta, these cells also showed elevated firing rate and somewhat higher rhythmicity frequency as measured by the first peak in their autocorrelograms, arguing for a stronger excitatory drive during theta (Figure 4A-B). Interestingly, we only found moderate changes in burst parameters, suggesting that the bursting mechanisms of MS pacemakers are mostly intrinsic and show only slight modulation with changing network states (Figure 4C). Since the change in burst parameters and rhythmicity frequency seemed too small to account for the observed elevated firing rate, we argued that pacemakers should be skipping more theta cycles during non-theta states. We confirmed this by calculating the proportion of cycles skipped during non-theta and theta activity (Figure 4D).

**Figure 4.**
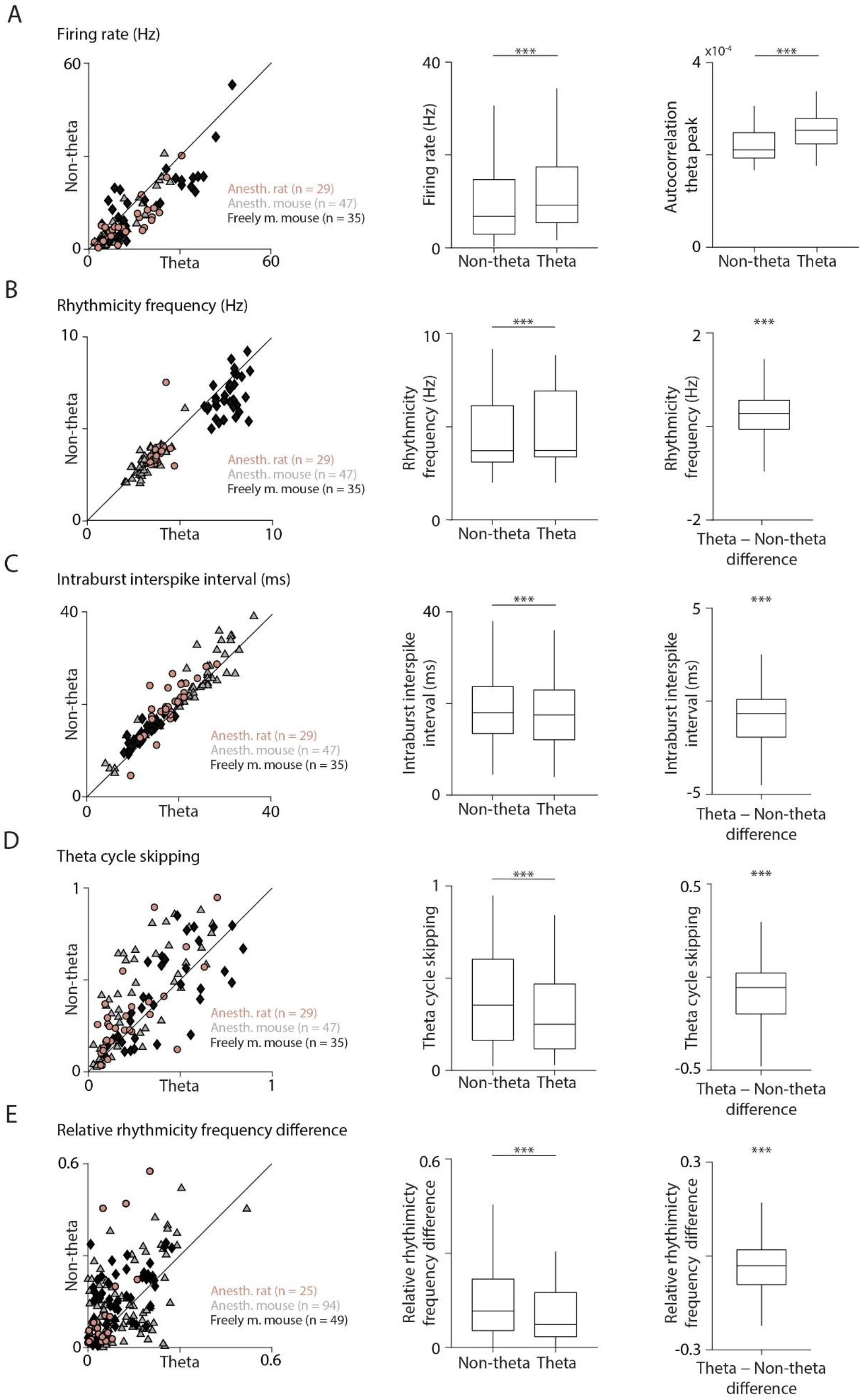
Septal pacemakers synchronize their burst frequencies. (A) Firing rates of MS putative pacemakers were higher during theta oscillation then during non-theta segments. Left, scatter plot shows the firing rate of all putative pacemaker neurons, color coded according to the rodent theta models. Middle, box-whisker plot showing statistics on pooled data. See Supplemental Table 2 for statistics on the three data sets separately. Right, autocorrelation peak in the theta frequency band. ***, p < 0.001, Wilcoxon signed-rank test for all statistical comparisons in this figure. All box-whisker plots show median, interquartile range and non-outlier range in this figure. (B) Left, scatter plot of rhythmicity frequency of putative pacemakers measured by the time lag of the first autocorrelation peak in the theta band. Middle and right, statistics on pooled data indicated higher rhythmicity frequency during hippocampal theta. (C) Left, scatter plot of average intraburst interspike intervals for putative pacemaker neurons. Middle and right, the moderate decrease of interspike intervals during theta oscillation indicates a slight elevation of intraburst frequency. (D) Left, scatter plot of the ratio of skipped theta cycles of putative pacemakers during non-theta and theta segments. Middle and right, statistics on pooled data indicated that pacemaker neurons skipped more theta cycles during non-theta segments. (To calculate theta skipping during non-theta segments, cycles were defined based on the autocorrelation peak of the neurons; see Methods). (E) Left, scatter plot of relative frequency difference of individual pacemaker neurons for simultaneously recorded pairs, normalized to the larger of the frequencies. Middle and right, statistics on pooled data indicated that induvial pacemakers became more similar in frequency during theta oscillation.

Next, we analyzed the pairs of putative pacemakers further to address how the synchrony of these neurons changes during hippocampal theta oscillation. Based on theoretical work, we hypothesized that pacemaker neurons may synchronize their frequencies, thus increasing constructive interference across individual oscillators, providing a stronger theta-rhythmic output towards the hippocampus (Ramirez et al., 2016; Ujfalussy and Kiss, 2006; Wang, 2002; Willms et al., 2017). This would imply that pairwise differences in rhythmicity frequency of putative pacemakers decrease during theta oscillation compared to non-theta episodes. Indeed, we found smaller differences in individual theta frequencies when hippocampal theta was present when we pooled all pacemaker pairs from the three data sets (Figure 4E). This difference was significant both for anesthetized rats and freely moving mice, and marginally significant when only anesthetized mice were considered (Supplemental Table 2). It is important to note that this frequency-synchronization was detected after normalizing the relative frequency difference to the individual frequencies of the tested neurons, thus a change in the individual theta frequencies could not account for this effect.

### Network model of the medial septum confirms frequency (Huygens)-synchronization mechanism

We reasoned that if increased tonic excitation leads to higher firing rates and frequency-synchronization in putative pacemakers that results in a strong theta-modulated output, as our data suggested, then we should be able to reproduce this finding in a simple network model of MS pacemakers.

We simulated putative pacemakers as single-compartment conductance-based model neurons. The pacemaker was based on a minimalistic model constructed to produce fast spiking burst dynamics and contained a transient sodium, a delayed rectifier, and a slowly inactivating D-type potassium channel (Figure 5A). The model neuron expressed rhythmic discharges upon current injection with a rhythmicity frequency in the delta-theta range, controlled by the inactivation time constant of the D-type potassium channel (Figure 5B). Septal pacemaker neurons are known to express strong H-currents (Kocsis and Li, 2004; Xu et al., 2004), hence we also included an HCN channel in the septal pacemaker model, which successfully reproduced the previously reported H-current characteristics (Figure 5C).

**Figure 5.**
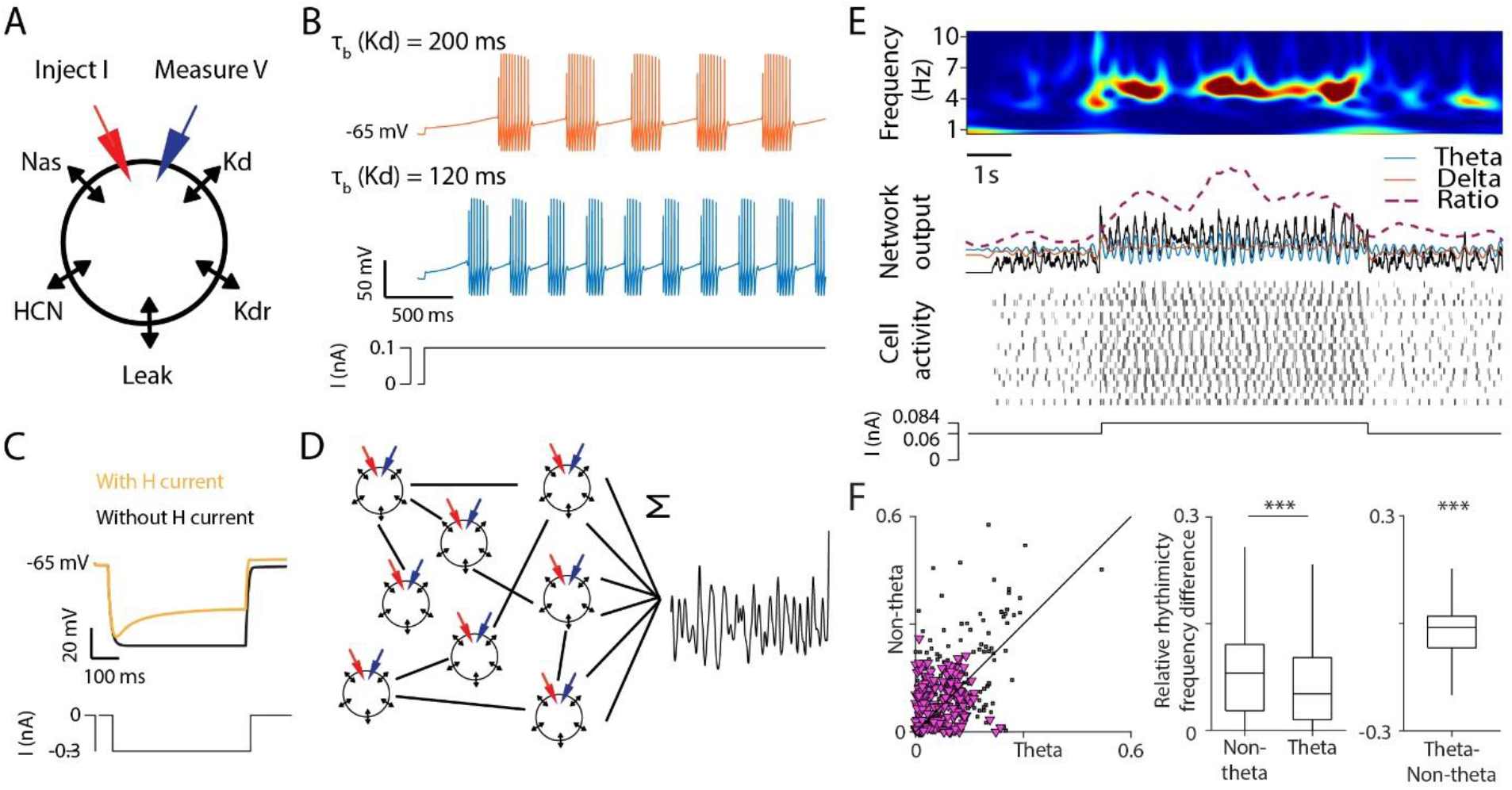
A homogeneous conductance-based network model of the MS pacemaker circuit is capable of theta-synchronization upon tonic excitation. (A) Schematic of the single-compartment model neuron. The inhibitory neuron was equipped with transient sodium (Nas), slowly inactivating D-type potassium (Kd), delayed rectifier potassium (Kdr), hyperpolarization activated cyclic nucleotide gated cation (I_H_) channels and passive leak channels. An electrode delivering current is attached to the cell (IClamp()). (B) Rhythmicity frequency of the model neuron can be controlled by the τ_b_(Kd) inactivation time constant. Slower inactivation (200 ms) also increases burst length. (C) Adding I_H_ caused a sag response to hyperpolarizing current, similar to MS GABAergic neurons reported in the literature (see e.g. Fig. 1C in (Xu et al., 2004)). The moderate rebound effect did not induce rebound spikes. (D) Schematics of the network. 20 model cells were connected using the following parameters: connection rate (CR), mean synaptic strength (variance was fixed at 10%), mean baseline stimulation strength, variance of the baseline stimulation strength. Spike trains were summed and convolved with a 50 ms Gaussian window to model network output. (E) Simulation of network behavior (50% connection rate; 0.5 nS mean synaptic strength; 60 pA mean baseline stimulation with 10% variance). Neurons synchronize in the theta frequency band in response to a tonic increase of stimulation and desynchronize when stimulation is reset to baseline. Top, wavelet spectrogram of network output. Middle, network output (black, raw; blue, filtered in the theta band; orange, filtered in the delta band; dashed, theta-delta ratio) and simulated spike raster. Bottom, injected current. (F) Left, scatter plot of relative frequency difference of individual pacemaker neurons for simultaneous pairs, normalized to the larger of the frequencies; real data in black (three data sets pooled, see Figure 4), model data overlaid in magenta (200 randomly selected data points). Middle and right, statistics on pooled data indicated individual model pacemakers became more similar in frequency during theta. Box-whisker plots show median, interquartile range and non-outlier range.

Next, we built a network model by connecting the pacemaker neurons with inhibitory synapses, simulating GABA-A receptor kinetics with a reversal potential at −70 mV that triggered fast IPSCs. All neurons received a non-rhythmic (‘tonic’) somatic excitatory current to mimic the changing excitatory drive to the septal pacemaker population. To model the impact of the medial septal output on the hippocampal local field potential, we convolved the averaged septal firing with a Gaussian kernel; this signal is referred to as ‘septal output’ hereinafter (Figure 5D). We found that at low tonic excitation levels, the septal output showed irregular activity with low levels of theta oscillation. Increasing the tonic drive resulted in a strong theta oscillatory output, which returned to the irregular activity when the tonic current was decreased (Figure 5E). Thus, the level of non-rhythmic excitation could toggle the model pacemaker network between theta and non-theta output, as suggested earlier (Bland et al., 1996; Hajszan et al., 2004) and consistent with our neural recordings. To test the robustness and parameter-dependence of this finding, we systematically explored the parameter space (Figure S18). We found a discrete maximum of the model’s synchronization capabilities at a mean baseline excitation of 60 pA, which was largely independent of the connectivity rate. Interestingly, very strong baseline excitation could result in a mirrored activity (decreased theta-synchronization upon increased excitation indicated by a synchronization score <0.5), sometimes also observed in anesthetized preparations. Fixing baseline tonic excitation at the optimum, the model network was capable of synchronizing at a range of mean synaptic strength levels and connection rates.

This model allowed us to test whether a homogeneous network of interconnected, minimalistic pacemaker units exhibits the same frequency-synchronization properties as we found in the septal network *in vivo*. To address this, we repeated the same battery of analyses on the model that we ran on the neural data. We found that model neurons increased their firing rate in theta state (Figure S19A), accompanied by a slight elevation in rhythmicity frequency (Figure S19B) and a strong reduction in theta-skipping (Figure S19D), with a consistent but relatively small change of burst properties (Figure S19C). When we compared pairwise differences in pacemaker frequencies, we found a strong theta-associated decrease (Figure 5F), suggesting the presence of a frequency-synchronization mechanism. Thus, the model behaved similarly to the rodent theta generating networks, confirming that Huygens-type frequency-synchronization might be a general mechanism operating in brain networks, similar to many mechanical systems.

### Most MS pacemakers are parvalbumin-expressing GABAergic neurons, whereas MS glutamatergic neurons are ‘tonic theta ON’

Finally, to investigate the genetic and neurotransmitter identity of septal neurons, we performed optogenetic tagging in urethane-anesthetized mice in a separate set of experiments. Theta oscillation was induced by tail pinch and we performed theta detection and septal neuron grouping based on rhythmicity as described before. We performed optogenetic stimulation in PV-Cre (n = 2), VGAT-Cre (n = 2) or VGluT2-Cre (n = 3) mice in order to identify parvalbumin-expressing, GABAergic and glutamatergic MS neurons, respectively (Gritti et al., 2006; Sotty et al., 2003).

We found that many PV-expressing neurons were theta-rhythmic (Figure 6A-H), and n = 10/18 neurons were characterized as putative pacemakers. Surprisingly, we only found a small fraction of rhythmic neurons among the identified GABAergic septal neurons, which were instead dominated by non-rhythmic and follower neurons (Figure 6I-P). Since a subset of GABAergic MS neurons express PV (Kiss et al., 1990; Unal et al., 2015), the few rhythmic GABAergic neurons may overlap with the previous PV-expressing population. These results suggest that parvalbumin-expression marks the theta generating population, confirming previous studies (Borhegyi et al., 2004; Freund, 1989; Varga et al., 2008).

**Figure 6.**
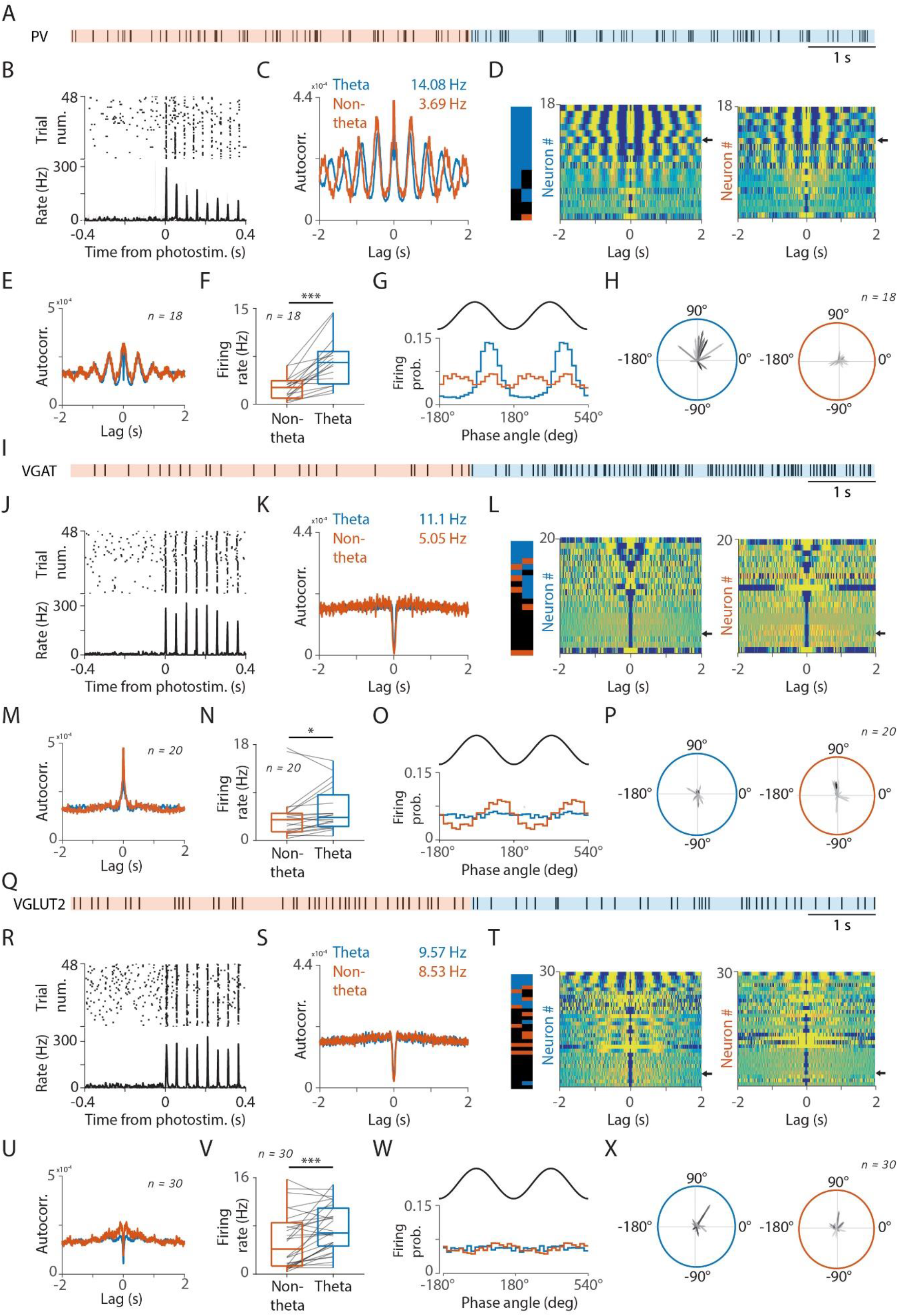
Optogenetic identification of MS neuron types in anesthetized mice. (A) Example spike raster of a PV-expressing MS neuron during non-theta (orange) and theta (blue) segments. (B) Raster plot (top) and corresponding peri-stimulus time histogram (bottom) of an example PV-expressing light-sensitive MS neuron aligned to the onset 20 Hz photostimulation. (C) Autocorrelograms of the example neuron during non-theta (orange) and theta (blue) segment. Numbers in the top right corner denote average firing rate during non-theta (orange) and theta (blue). (D) Autocorrelograms of all identified PV-expressing MS neurons, sorted by rhythmicity index. Left, color bar indicates significant theta-rhythmicity (blue) and delta-rhythmicity (orange) during non-theta (left) and theta (right) segments. Middle, autocorrelograms during theta segments. Right, autocorrelograms during non-theta segments. The arrows indicate the example neuron. Most PV-positive MS neurons were pacemakers. (E) Average autocorrelogram of all identified PV-expressing MS neurons. (F) Firing rate change between non-theta and theta segments of all identified PV-expressing MS neurons (n = 18). (G) Phase-locking of the example neuron to delta (orange) and theta (blue) oscillations. (H) Phase-locking of all identified PV-expressing MS neurons to theta (left) and delta (right) oscillations. Greyscale indicates average firing rate. (I) Example spike raster of a VGAT-expressing MS neuron during non-theta (orange) and theta (blue) segments. (J) Raster plot (top) and corresponding peri-stimulus time histogram (bottom) of an example light-sensitive VGAT-expressing MS neuron aligned to the onset 20 Hz photostimulation. (K) Autocorrelograms of the example neuron during non-theta (orange) and theta (blue) segment. Numbers in the top right corner denote average firing rate during non-theta (orange) and theta (blue). (L) Autocorrelograms of all identified VGAT-expressing MS neurons, sorted by rhythmicity index. Left, color bar indicates significant theta-rhythmicity (blue) and delta-rhythmicity (orange) during non-theta (left) and theta (right) segments. Middle, autocorrelograms during theta segments. Right, autocorrelograms during non-theta segments. The arrows indicate the example neuron. Most VGAT-positive MS neurons were non-rhythmic or follower neurons. (M) Average autocorrelogram of all identified VGAT-expressing MS neurons. (N) Firing rate change between non-theta and theta segments of all identified VGAT-expressing MS neurons (n = 20). (O) Phase-locking of the example neuron to delta (orange) and theta (blue) oscillations. (P) Phase-locking of all identified VGAT-expressing MS neurons to theta (left) and delta (right) oscillations. Greyscale indicates average firing rate. (Q) Example spike raster of a VGLUT2-expressing MS neuron during non-theta (orange) and theta (blue) segments. (R) Raster plot (top) and corresponding peri-stimulus time histogram (bottom) of an example VGLUT2-expressing light-sensitive MS neuron aligned to the onset 20 Hz photostimulation. (S) Autocorrelograms of the example neuron during non-theta (orange) and theta (blue) segment. Numbers in the top right corner denote average firing rate during non-theta (orange) and theta (blue). (T) Autocorrelograms of all identified VGLUT2-expressing MS neurons, sorted by rhythmicity index. Left, color bar indicates significant theta-rhythmicity (blue) and delta-rhythmicity (orange) during non-theta (left) and theta (right) segments. Middle, autocorrelograms during theta segments. Right, autocorrelograms during non-theta segments. The arrows indicate the example neuron. Few VGLUT2-positive MS neurons showed theta-rhythmicity. (U) Average autocorrelogram of all identified VGLUT2-expressing MS neurons. (V) Firing rate change between non-theta and theta segments of all identified VGLUT2-expressing MS neurons (n = 30). (W) Phase-locking of the example neuron to delta (orange) and theta (blue) oscillations. (X) Phase-locking of all identified VGLUT2-expressing MS neurons to theta (left) and delta (right) oscillations. Greyscale indicates average firing rate.

The rhythmicity properties of MS glutamatergic neurons have not been investigated before, leaving all assumptions about their role in theta generation at best uncertain. We found that the overwhelming majority of optogenetically identified MS glutamatergic neurons showed strong increase of firing rate (fold increase median ± s.e. of median, 1.42 ± 0.34) without any theta-rhythmicity during hippocampal theta state (Figure 6Q-X). Only few glutamatergic neurons were theta-rhythmic, 4/30 characterized as pacemakers. This identifies glutamatergic neurons as previously unidentified ‘tonic theta ON’ neurons, first described by the seminal studies of Bland and colleagues (Ford et al., 1989). Thus, local glutamatergic neurons may contribute to the increased excitatory drive to the pacemaker neurons during theta synchronization, whereas it is less likely that they form an integral part of the rhythm generation network.

## Discussion

We categorized medial septal neurons based on their rhythmic properties. Putative pacemaker neurons showed constitutive theta rhythmic activity and synchronized their frequencies during hippocampal theta oscillations in different rodent theta model systems. This mechanism, known as Huygens synchronization, was reproduced by a network model of minimalistic MS pacemaker units, demonstrating that a homogeneous group of pacemakers can synchronize and desynchronize via inhibitory connections upon changing levels of tonic excitatory drive. Optogenetic tagging confirmed that pacemaker neurons are PV-expressing GABAergic cells, while non-PV GABAergic neurons likely do not participate in rhythm genesis but rather ‘follow’ the ongoing activity. Glutamatergic septal neurons were ‘tonic theta ON’ type, which suggests that they contribute to the non-rhythmic excitatory control of the pacemaker rhythm generator network.

The septo-hippocampal pathway through the fimbria-fornix was identified as a major input to the Cornu Ammonis as early as 1909 by Cajal (Ramón y Cajal, 1909). After Jung and Kornmüller described the 4-7 Hz rhythmic activity in the rabbit hippocampus (Jung and Kornmüller, 1938) that was later coined the theta rhythm (Green and Arduini, 1954), the link between the two was established with multiple techniques. Septal lesions, either complete, selective cholinergic or GABAergic, eliminated, whereas stimulating the MS electrically, or cholinergic, glutamatergic or GABAergic MS neurons optogenetically, could elicit hippocampal theta (Bland et al., 2007; Dannenberg et al., 2015; Green and Arduini, 1954; Lee et al., 1994; Leung and Shen, 2004; Robinson et al., 2016; Roland et al., 2014; Smythe et al., 1992; Vandecasteele et al., 2014; Yoder and Pang, 2005). Furthermore, MS neurons fired rhythmically phase locked to hippocampal theta, and some exhibited theta-rhythmic properties even in the absence of hippocampal theta rhythm (Borhegyi et al., 2004; Petsche et al., 1962; Stewart and Fox, 1989; Varga et al., 2008). These findings quickly put the medial septum on the center stage in theta generation. However, the mechanism by which MS neurons achieve hippocampal theta has proven to be a hard problem.

First, MS cell types presented a puzzle. While early pharmacological evidence pointed to the cholinergic septo-hippocampal projection, lesion, anatomical and recording studies indicated that GABAergic and glutamatergic neurons are also involved in theta formation (Freund and Antal, 1988; Hajszan et al., 2004; Leung and Shen, 2004; Manseau et al., 2005; Robinson et al., 2016; Smythe et al., 1992; Sotty et al., 2003; Yoder and Pang, 2005). Therefore, we chose to start from an unbiased physiological characterization of MS units, extending previous electrophysiology studies (Apartis et al., 1998; Borhegyi et al., 2004; Duque et al., 2000; Ford et al., 1989; Hassani et al., 2009; Simon et al., 2006; Stewart and Fox, 1989; Varga et al., 2008). Bland and colleagues characterized MS neurons as theta-on and theta-off based on their firing rates in theta and non-theta states; cells were also grouped as ‘phasic’ (theta-bursting) or ‘tonic’ (non-rhythmic) based on their firing patters. Other studies indicated the presence of lower frequency rhythms including ‘slow oscillations’ and rhythmic activity related to breathing (Buzsaki et al., 1986; Tsanov, 2015; Tsanov et al., 2014; Wolansky et al., 2006). In accordance, we also observed rhythmic activity in the delta frequency band, especially in urethane-anesthetized rats. Therefore, we characterized MS neurons based on their rhythmicity during hippocampal theta and non-theta episodes as non-rhythmic, delta-rhythmic or theta-rhythmic. While this resulted in 9 theoretical combinations, we found that most MS neurons belonged to a few characteristic types.

Stewart and Fox found that after eliminating theta oscillation in the hippocampus of urethane-anesthetized rats using atropine, a subset of MS neurons remained theta rhythmic (Stewart and Fox, 1989). In another study, 15% of septal neurons were found phasic bursting even during non-theta epochs under anesthesia (Ford et al., 1989). We identified putative pacemaker neurons that fired theta-rhythmic bursts regardless whether theta oscillation was present in CA1. This group comprised 13% of MS neurons in freely moving mice; however, we found a lower percentage of rhythmic bursting neurons in anesthetized rodents, in line with some previous reports (King et al., 1998; Stewart and Fox, 1989). The differences in percentage of pacemakers across reports may stem both from biased sampling, as the MS can be identified during recording based on the presence of theta-rhythmic activity, as well as from strictness of definitions for rhythmicity. We applied relatively conservative inclusion criteria for significant rhythmicity (Figure S3), in order to ensure data quality necessary for analyzing mechanisms of rhythmic synchronization, which may have resulted in a lower percentage of putative pacemakers in our data sets.

The presence of this putative pacemaker group has been confirmed many times (Borhegyi et al., 2004; Ford et al., 1989; Hangya et al., 2009; Joshi et al., 2017). Based on anatomical studies (Freund and Antal, 1988; Freund and Gulyás, 1991; Unal et al., 2015), selective lesions (Roland et al., 2014; Yoder and Pang, 2005) and firing patterns of identified GABAergic neurons (Borhegyi et al., 2004; Brazhnik and Fox, 1997; Hangya et al., 2009; Hassani et al., 2009; Serafin et al., 1996; Varga et al., 2008), it was suggested that hippocampally projecting GABAergic neurons form the septal theta pacemaker circuit. More specifically, the expression of PV and HCN, the channel protein that underlies the H-current, has been associated with rhythmic pacemaking properties (Borhegyi et al., 2004; Kocsis and Li, 2004; Varga et al., 2008; Xu et al., 2004). However, other studies suggested that rhythmic MS glutamatergic or cholinergic neurons may act as the pacemakers (Manseau et al., 2005; Mysin et al., 2015; Smythe et al., 1992; Ujfalussy and Kiss, 2006) (but see Ref. (Vandecasteele et al., 2014)). Here, we confirmed by optogenetic identification of multiple MS cell types that PV-expressing GABAergic neurons probably constitute the majority of the constitutive theta rhythmic group. We found that non-PV GABAergic neurons, despite some earlier suggestions (Simon et al., 2006), mostly fire non-rhythmically; similarly, little rhythmic activity was found among glutamatergic MS neurons. We confirmed that septal pacemakers formed two groups with opposing phase preference with respect to the hippocampal theta oscillation, as reported before (Borhegyi et al., 2004; Joshi et al., 2017; Varga et al., 2008).

In the 17^th^ century, Christiaan Huygens observed that two pendulum clocks, coupled through a house beam, tended to synchronize over time, which he referred to as ‘sympathy of two clocks’ (Huygens, 1673; Oliveira and Melo, 2015; Ramirez et al., 2016; Willms et al., 2017). Such synchronization of coupled oscillators has been found in many dynamical systems and has been hypothesized to be present in neuronal networks as well (Equihua and Ramirez, 2018; Ramirez et al., 2016). This would mean that individual oscillators become more similar to each other in their frequency and period time. We identified this mechanism among septal pacemaker neurons, suggesting that a Huygens synchronization mechanism underlies the generation of a strong theta rhythmic output transmitted to the hippocampus. The two dominant oscillatory behaviors that emerge in coupled oscillators during Huygens synchronization are in-phase and anti-phase synchronization (Korteweg, 1906; Ramirez et al., 2014). This may, at least in part, underlie the observation of anti-phase firing of PV-expressing neuron groups in the MS. Importantly, experiments and modeling studies observed less common modes of Huygens synchronization, including partial synchronization, when frequencies are synchronized but the phase difference between the oscillators is less than 180 degrees; ‘quenching’, when the two oscillators suppress each other; ‘beating death’, in which one oscillator is suppressed but remains oscillatory due to the influence of the other, and amplitude modulation in consequence of mutual influence (Ramirez et al., 2014). It will be exciting to study whether and how the broad theory behind Huygens synchronization, often investigating limit behavior of complex non-linear ordinary differential equation systems, can be applied to neural circuits.

Previous models of the septal pacemaker network either explored GABAergic (Denham and Borisyuk, 2000; Ujfalussy and Kiss, 2006; Wang, 2002) or glutamatergic (Ujfalussy and Kiss, 2006) pacing mechanisms. These previous works already contained indirect reference to potential Huygens synchronization mechanisms. For instance, Wang suggested in his seminal theoretical work that desynchronized septal pacemaker units may oscillate at different frequencies (see Fig.4 in ref. (Wang, 2002)). Additionally, Ujfalussy et al. wrote that ‘blocking the GABAergic synapses in our ping-pong model causes desynchronization of these neurons but the cells remain theta periodic.’ However, Huygens synchronization has neither been experimentally tested nor directly modelled, and previous models mostly focused on exploring the ways different oscillator networks may interplay, such as two GABAergic MS groups with opposing firing phase (Ujfalussy and Kiss, 2006), MS and CA1 GABAergic groups (Denham and Borisyuk, 2000; Wang, 2002; Zou et al., 2011) or MS GABAergic and glutamatergic neurons (Mysin et al., 2015). We found that even a minimalistic, homogeneous network of individual MS GABAergic pacemakers may be capable of generating coherent theta-rhythmic output upon increased tonic excitation. While desynchronization of the MS network was rarely addressed in previous theoretical work, we also found that our MS model network showed rapid spontaneous desynchronization when tonic excitation was decreasing. This MS model network could recapitulate multiple features of the in vivo synchronization process including Huygens synchronization, increased firing rate, faster rhythmic firing, decreased theta-skipping and a slight elevation of intra-burst firing frequency. We based this model on a previously published minimalistic conductance-based model capable of rhythmic bursting (Golomb et al., 2007). Since H-current has been consistently demonstrated in MS pacemaker neurons, we equipped this model neuron with HCN-channels that match the reported properties of MS neurons (Morris et al., 2004; Sotty et al., 2003). While it is still debated whether the H-current is a major determinant of MS rhythmic pacing (Kocsis and Li, 2004; Xu et al., 2004), our model may provide a tool to further explore this question in later studies. Finally, although we demonstrated that a homogeneous group of MS inhibitory neurons may be capable of pacing hippocampal theta, this does not rule out the possibility of other contributing neuronal populations. Notably, clarifying the potential role of the hippocampo-septal feedback projection (Katona et al., 2017; Toth et al., 1993; Wang, 2002) requires further investigations.

We found a substantial elevation of firing rates in MS pacemaker neurons upon theta onset. This finding is in line with previous suggestions that an increased tonic excitatory drive can switch the MS network into theta state (Lu et al., 2011; Müller and Remy, 2018; Oddie et al., 1996; Ujfalussy and Kiss, 2006). Multiple cell types and pathways were proposed to contribute to this excitatory drive including MS cholinergic, glutamatergic and ascending brainstem inputs (Hajszan et al., 2004; Leranth and Frotscher, 1989; Müller and Remy, 2018; Oddie and Bland, 1998; Zaborszky et al., 2012). We found that identified MS glutamatergic neurons strongly increased their firing rates in a non-rhythmic fashion at the onset of hippocampal theta oscillation. Therefore, we suggest that MS glutamatergic neurons may be the dominant relay that conveys changing levels of tonic excitation to the MS pacemaker network, in agreement with anatomical findings showing that local excitatory neurons provide the majority of excitatory inputs to MS GABAergic neurons (Hajszan et al., 2004). Our findings also suggest that MS glutamatergic neurons constitute a large part of the ‘tonic theta ON’ group defined by Bland and colleagues (Ford et al., 1989). This reasoning also agrees with the growing body of evidence showing that MS glutamatergic neurons are involved in conveying locomotion-related activation to other cell types of the MS and beyond (Fuhrmann et al., 2015; Justus et al., 2017; Oddie et al., 1996; Teitelbaum et al., 1975).

Previous studies demonstrated theta-skipping behavior of rhythmic neurons both in the entorhinal cortex (Brandon et al., 2013; Jeffery et al., 1995) and in the medial septum (Kay et al., 2020; King et al., 1998; Varga et al., 2008). Theta-skipping was characterized by rhythmic activity with double the period length, i.e. half the frequency of ongoing hippocampal theta oscillation, by neurons firing rhythmically during every second theta cycle. Importantly, Brandon and colleagues elegantly demonstrated that the presence of medial septal input was necessary for theta-skipping behavior in the entorhinal cortex in rats (Brandon et al., 2013). Moreover, Hasselmo showed that theta-skipping behavior in the entorhinal cortex may rely on inhibitory rebound mechanisms, potentially crucial to the formation of characteristic grid cell firing fields in the entorhinal cortex (Hasselmo, 2014). We confirmed the presence of theta-skipping neurons in the rat medial septum in urethane-anesthetized animals. While some of these neurons fired regularly in every second cycle, other neurons showed a less regular pattern, occasionally skipping more than one cycle. Interestingly, we found very few theta-skipping neurons in mice, either under anesthesia or in freely moving animals. Hasselmo demonstrated that theta-rhythmic input from the medial septum combined with inhibitory rebound mechanisms are sufficient to generate entorhinal theta-skipping (Hasselmo, 2014); therefore, theta-skipping in the MS may not be necessary for grid pattern formation in mice.

We found a group of regular rhythmic, tonically active neurons both in rats and mice. It is tempting to speculate that these neurons may be cholinergic. First, their firing patterns resemble cholinergic neurons in previous publications (Brazhnik and Fox, 1997; Mamad et al., 2015; Tsanov, 2015). Second, we identified regular rhythmic cholinergic neurons from other structures of the basal forebrain (Laszlovszky et al., 2020). Third, Lee et al. found rhythmically firing basal forebrain cholinergic neurons that were not phase locked to hippocampal theta (Lee et al., 2005). Strikingly, we also found that these neurons, despite firing rhythmically at theta range frequencies, showed little correlation with ongoing theta or other rhythmic groups of the septal network. Jones and colleagues proposed that theta rhythmic cholinergic neurons may be more linked with a neocortical theta oscillation that can be recorded from the posterior cingulate cortex (Lee et al., 2005). In accordance, Destrade and Ott proposed a retrosplenial or cingulate cortical theta source (Destrade and Ott, 1982) and we have also reported behavior-specific synchronization of regular rhythmic nucleus basalis cholinergic neurons with theta-band activity in the auditory cortex (Laszlovszky et al., 2020). This would confirm the notion that the role of septal cholinergic neurons in hippocampal theta generation is mostly indirect through GABAergic neurons, controlling theta amplitude but not participating in rhythmic theta pacing (Dannenberg et al., 2015; Lee et al., 1994; Yang et al., 2014, 2017). We expect that it will be one of the most exciting future directions to better understand how multiple theta rhythms interact dynamically in the brain, including synchronization with active sensing processes (Moore et al., 2013; Ranade et al., 2013; Semba and Komisaruk, 1984), entraining local pacemaker groups (Manseau et al., 2008) and interacting with cellular resonance properties (Zemankovics et al., 2010).

We found a strong correspondence of rhythmic medial septal cell types and synchronization mechanisms across urethane-anesthetized rats, urethane-anesthetized mice and feely moving mice. This argues for common general mechanisms for septo-hippocampal theta-rhythmic synchronization. However, we also noted some differences. Part of these differences may stem from known distinctions of urethane-and atropine-sensitive theta oscillations (Kramis et al., 1975; Li et al., 2007), with different underlying mechanisms (Losonczy et al., 2010; Mikulovic et al., 2018; Nicola and Clopath, 2019; Winterer et al., 2019), the former abolished during urethane anesthesia. Indeed, we confirmed the higher frequency of theta oscillations in freely moving mice (Buzsáki and Moser, 2013; Varga et al., 2008). Nonetheless, we also found cross-species differences. Most notably, anesthetized rats showed strong hippocampal rhythmicity in the delta band during non-theta episodes, mostly absent in mice. In accordance, we found delta-rhythmic septal neurons in rats but very few in mice. Maybe relatedly, theta-skipping neurons were found in rats but rarely in mice. Another interesting difference was the different behavior of tonically active neurons upon theta: their firing rate decreased in rats but increased in mice. Marder and colleagues observed that often the rhythmic firing activities are controlled and not the exact underlying ion channel constellations, leading to unexpected variability in channel abundance with stable oscillatory parameters (Alonso and Marder, 2019). It is possible that, similar to this principle, the theta-rhythmic activity of the septo-hippocampal system is evolutionarily conserved, with a larger variance in the exact implementation (Payne et al., 2020). Further experiments and modeling will help gauge the significance of this variability.

## Methods

### Animals

Wild type C57BL/6 mice (n = 11; 27.4 g ± 4.9 g, 7 males) were used for acute mouse recordings; Wistar rats (N = 7; 200–400 g; males) were used for acute rat recordings; Sst-IRES-Cre mice with C57BL/6J genetic background (n = 4; 28-30 g; males) were used for the chronic mouse recordings; 4 PV-IRES-Cre males (genetic background FVB/AntFx), 4 VGAT-IRES-Cre males (genetic background Bl6Fx) and 5 BAC-Vglut2-Cre male and female mice (genetic background C57BL/6J) were used for optogenetic tagging. All experiments were performed according to the EC Council Directive of September 22, 2010 (2010/63/EU), with all procedures being reviewed and approved by the Animal Care Committee of the Research Centre for Natural Sciences or the Institutional Animal Care and Use Committee of the Institute of Experimental Medicine and by the National Food Chain Safety Office of Hungary.

### Surgery and electrophysiological recordings

#### Acute rat recordings

Rats were anesthetized with an intraperitoneal injection of urethane (40%; dose, 0.37 ml/100 g). A homeothermic heating pad connected to a rectal probe held body temperature constant (36°C). The top of the head was shaved, the rat was placed in a stereotaxic frame (David Kopf Instruments, Tujunga, US), the skin and the connective tissue above the calvaria were removed and the skull was cleared. A craniotomy was opened over the medial septum and another one over the right hippocampus. A 32-channels linear silicon probe (A1×32-6mm-50-177; NeuroNexus Technologies, Ann Arbor, US) was lowered into the right CA1 (anterior-posterior (AP): −4.5 mm and medial-lateral (ML): 3 mm) and a 32-channels Buzsaki-type silicon probe (Buzsaki32; NeuroNexus Technologies, Ann Arbor, US) was lowered to the top of the medial septum (AP: 0.4 mm and ML: −1.6 mm, 15° angle in the coronal plane). The silicon probes were dipped in red fluorescent die (1,1′-Dioctadecyl-3,3,3′,3′-Tetramethylindocarbocyanine Perchlorate, DiI) before insertion to aid later histological reconstruction. The ground electrode was secured in the nuchal muscles.

Neural signals were acquired by two National Instruments PCI-6259 cards, amplified (2000 times) and digitized at 20 kHz. After allowing 30 minutes for the brain tissue around the electrode to stabilize, a 30-minutes recording was conducted. Theta oscillation was induced by temporarily placing an insulated clip on the tail for 1 minute, repeated 3 times. After the recording, the septal probe was moved 100 μm ventrally, and the procedure was repeated. Recording sessions were conducted as long as theta oscillation could be detected via audio feedback; correct positioning of the probe in the MS was later confirmed by both histological reconstruction (Figure S1) and the presence of theta-rhythmic neurons in the recordings.

#### Acute mouse recordings

Implantations were done under general anesthesia. Mice received an intraperitoneal injection of urethane (~1.3 mg/g). If necessary, an additional dose of urethane was injected intramuscularly to maintain the depth of the anesthesia during surgery and recordings. The body temperature of the animals was maintained during the experiments at 37 °C using a thermostatically regulated heating pad (Supertech, Pécs, Hungary). Mice were placed in a stereotaxic frame (David Kopf Instruments, Tujunga, US), then the skin and the connective tissue were removed from the top of the skull. Next, two small circular craniotomies (1 mm diameter) were made on the skull with a dental drill, one for the recording probe inserted into the MS and one for the probe inserted into the hippocampus (CA1). Stereotaxic coordinates were used to determine the septal and hippocampal locations for recording (MS, AP: 0.6 mm and ML: 0.5 mm; CA1, AP: −2.2 mm and ML: 1.5 mm from the Bregma) (Paxinos et al., 2001). For post-mortem histological verification of the recording location of the probes, the backside of the silicon shank was coated with red-fluorescent dye (DiI, D-282, ~10% in ethanol, Thermo Fischer Scientific, Waltham, US) before insertion (DiCarlo et al., 1996; Fiáth et al., 2019). The septal probe was a high-density single-shank silicon probe with 128 square-shaped recording sites (Fiáth et al., 2018). The 8-mm-long shank of the probe had a cross-section of 100 μm × 50 μm (width × thickness). The closely spaced recording sites (20 μm × 20 μm) were arranged in a 32 x 4 array with 2.5 μm spacing between the edge of electrodes. The probe was mounted on a motorized micromanipulator (Robot Stereotaxic, Neurostar, Tübingen, Germany) and inserted at a slow speed (2 μm/sec) into the brain tissue to increase the single unit yield by decreasing the tissue damage (Fiáth et al., 2019). The probe was tilted at an angle of 8° from vertical in the coronal plane to have better access to the medial septum. Since the effective vertical recording area of the probe (~0.7 mm) was smaller than the dorsoventral extent of the MS (~1.5 mm), to obtain the spiking activity of as many septal neurons as possible, recordings were performed in three, slightly overlapping (~0.2 mm) depths. (Unit clusters recorded twice due this overlap, based on waveform similarity and autocorrelograms, were counted only once.) The hippocampal probe was a linear silicon probe with 32 recording sites and with a shank thickness of 50 μm (A1×32-6mm-50-177, NeuroNexus Technologies, Ann Arbor, US). The recording sites of the device had a diameter of 15 μm and an interelectrode distance of 50 μm. The probe was attached to a manual micromanipulator, then inserted at a speed of ~10 μm/sec to a dorsoventral depth of 2 mm. The two recording probes were connected to an electrophysiological recording system (RHD2000, Intan Technologies, Los Angeles, CA, USA) comprising 32-channel and 64-channel headstages.

Wideband brain electrical activity (0.1 – 7500 Hz) was recorded on 160 channels with 16-bit resolution and at 20 kHz/channel sampling rate. A stainless steel wire inserted into the neck muscle of the animal served as the reference and ground electrode during recordings. Room temperature physiological saline solution was regularly dropped into the craniotomy to prevent dehydration of the brain tissue. Recordings were conducted for 15 minutes each, consisting of 5 minutes baseline period, 5 minutes tail-pinch-induced theta oscillation, and 5 minutes recovery from stimulation.

#### Chronic mouse recordings

Sst-IRES-Cre male mice of C57BL/6J genetic background underwent bilateral injection of AAV2/5-Ef1a-DIO-ChR2-YFP-WPRE (N = 2) or AAV5-Ef1a-DIO-SwiChRca-TS-EYFP-WPRE (N = 2) into the dorsal hippocampi (coordinates: AP −2.1 and −2.5 mm and ML ±1.5 and 1.6 mm from Bregma with tip at DV −1.2 mm from brain surface; 2 × 150-200 nl on both sides) using standard surgery techniques. Silicon probes for multichannel electrophysiological recordings were implanted above the medial septum and into the dorsal hippocampus 40-83 days after the virus injection. The surgery was performed under isoflurane anesthesia induced by ketamine-xylazine (4:1) combination diluted 6x in Ringer’s lactate solution (intraperitoneal injection, dose 0.01 ml/1g body weight). Mice were head-fixed in a stereotaxic frame (David Kopf Instruments, Tujunga, US) and their body temperature and respiratory rate were continuously monitored. After local disinfection (Betadine) and analgesia (10% Lidocaine-spray), the cranium was exposed and cleaned for application of adhesive agent (OptiBond XTR, Kerr Corporation, Orange, US). Craniectomies were performed for stereotaxis-guided implantation of a 32-channels linear type and a 32-channels, 4-shank Buzsáki-type silicon probe into the dorsal hippocampus (coordinates: AP −2.5 mm and ML + 2 mm from Bregma with tip at DV −2.1 mm from brain surface) and above the medial septum (AP +0.9 mm and ML +0.9 mm, at an angle of 12°, and with tip at DV −2.8 mm from brain surface), respectively. Both probes were mounted on a custom-made, adjustable micro-drive. The probes were coated with red fluorescent DiI for later histological confirmation of the recording site. After the recovery from the surgery, the tip of the septal probe was lowered in steps of 75-150 micrometers per day to reach the zone of cells with theta-modulated firing pattern. Within this zone, the probe was advanced in steps of 45 micrometers per day as long as the theta-modulated firing pattern was still present. In case of SwiChR-transfected animals, each shank of the septal probe was equipped with an optical fiber (50 micrometers core diameter, 0.22 NA) with tip positioned 75-100 micrometers above the uppermost recording site and glued by optical adhesive. In case of ChR2-transfected animals, one optical fiber (105 micrometers core diameter, 0.22 NA) was implanted to illuminate the fimbria, independently from the septal silicon probe. In one animal, a Buzsáki-type silicon probe was implanted into the hippocampus, instead of the linear probe. In this case, the probe was advanced stepwise to reach the pyramidal layer of the dorsal hippocampus after the post-surgery recovery period. The craniectomies were sealed with artificial dura (Cambridge NeuroTech Ltd, Cambridge, UK). The probe-micro-drive assemblies were shielded by copper mesh preventing the contamination of the recordings by environmental electric noise. The mesh was covered by dental acrylate. Two stainless steel wires inserted above the cerebellum served as ground and reference for the electrophysiological recordings. Before finishing the surgery, Buprenorphine (dose: 0.045 μg/1g body weight) was injected subcutaneously. Recordings were started after a one-week-long post-surgery recovery and habituation to connectorization.

The electrophysiological activity was registered by a multiplexing data acquisition system (KJE-1001, Amplipex Ltd, Szeged, Hungary) at 20 kHz sampling rate. The position of the animal was tracked by a marker-based, high speed (120 frame/s) motion capture system and reconstructed in 3D (Motive, OptiTrack, NaturalPoint Inc, Corvallis, US). Recordings were conducted both in the animals’ home cage (dominated by sleep) and while mice were placed on a linear track (dominated by active movement). Home cage and linear track recordings were concatenated for data analysis to include sufficient amount of time spent in theta and non-theta states. Thus, the majority of theta segments originated from movement epochs, while non-theta segments represented both quiet wakefulness and sleep. As the main focus was on general oscillatory mechanisms, these states were not subdivided further. Optogenetic manipulation of somatostatin-expressing neurons was carried out separately; those recordings were not analyzed in the present study. The septal probe was lowered 45 micrometers after the recordings each day, through a total of 8-35 days of recording.

#### Optogenetic tagging in mice

4 PV-IRES-Cre males (genetic background FVB/AntFx), 4 VGAT-IRES-Cre males (genetic background Bl6Fx) and 5 BAC-Vglut2-Cre males and females (genetic background C57BL/6J) were used to assess the neurochemical nature of MS cells; 2 PV-Cre (n = 17, 1), 2 VGAT-Cre (n = 17, 3) and 3 VGLUT2-Cre (n = 15, 9, 6) mice yielded tagged neurons (numbers in brackets). Mice were anesthetized with an i.p. injection of ketamine-xylazine (0.166 and 0.006 mg/kg, respectively). After shaving and disinfecting (Betadine) the scalp, local anesthetic was applied (Lidocaine). Animals were positioned in the stereotaxic frame and the eyes were protected with eye ointment (Laboratories Thea). After opening the skin, the skull was cleaned and the head of the animal was leveled using Bregma, Lambda and a pair of lateral points equidistant from the sagittal suture.

A trephine hole was performed in order to access the MS with a 10º lateral angle (MS 10°, antero-posterior +0.90 mm, lateral, 0.90 mm). An adeno-associated virus vector allowing Cre-dependent expression of channelrhodopsin2 [AAV 2/5. EF1a.Dio.hChR2(H134R)-eYFP.WPRE.hGH] was injected into the MS at 3.95, 4.45 and 5.25 mm depth from skull surface (200 nl at each depth, 100 nl at 25 nl s^−1^ and 100 nl at 5 nl s^−1^). After the viral injections, the skin was sutured and local antibiotics (Neomycin) and analgesics (Buprenorphine 0.1 mg kg-1, s.c.) were applied.

Around two weeks after the virus injection, the animals were anesthetized with an i.p. injection of 20% urethane (Sigma-Aldrich, 0.007 ml g-1 body weight). The depth of anesthesia was evaluated by pinching the paw, tail, or ear of the animal, and by checking the ocular reflex. When no reflexes were elicited, the throat was shaved, and topical lidocaine was applied. A heating pad was used to keep constant body temperature of the animals. A tracheotomy was performed in order to sustain a constant airflow (Moldestad et al., 2009). The animals were placed in a stereotaxic frame and, after opening the skin and leveling the skull, a cranial window was made above the MS (silicon probe MS 10°, antero-posterior, +0.90 mm, lateral, 0.90 mm; optic fiber MS 5º contralateral, antero-posterior, +0.90 mm, lateral, −0.50 mm), the hippocampus (silicon probe HPC, antero-posterior, −2.20 mm, lateral, 1.50 mm) and two above the cerebellum for reference electrode placement. A Neuronexus A1×32-6mm-50-177-CM32 silicon probe was placed in the hippocampus at 2.20 mm depth from skull surface. A Neuronexus Buzsaki32-H32_21 mm probe was lowered to the dorsal boundary of the MS at a 10º lateral angle (3.95 mm from skull surface). Reference electrodes for both probes were placed in the cerebellum, keeping around 5 mm distance between them, and a ground electrode was placed in the spinotrapezius muscle. A 200 μm core optic fiber (Thorlabs) was lowered 500 μm above the shanks of the MS probe. The MS probe and the optic fiber were lowered in consecutive recordings (100 μm steps for the probe; 50 μm steps for the optic fiber), spanning the entire depth of the MS. Extracellular data were collected by the Open Ephys data acquisition system, digitized at 30 kS/s. Each recording session consisted of an optical tagging period of two minutes, followed by a baseline period of five minutes of spontaneous activity. Three consecutive repetitions of one-minute tail pinch induced theta activity followed by one-minute control recording were applied, finishing the recording session with another two-minute-long optical tagging period. For optimizing the probability of finding tagged cells, we used the Online Peri-Event Time Histogram (OPETH) Open Ephys compatible plugin that allowed us to visualize the direct effect of the optogenetic activation at the cell population level. OPETH (Széll et al., 2020) allowed us to reduce recording time by more efficient ‘hunting’ for light responsive neurons in the MS and to reduce light-triggered artifacts in our recordings, improving data quality and further analysis.

#### Histological verification of the recording location

To detect the tracks of the silicon probes in the brain tissue, we used a histological procedure similar to that described previously (Borhegyi et al., 2004; Fiáth et al., 2018, 2019). In brief, the animal was deeply anesthetized after the recordings, then transcardially perfused with physiological saline solution followed by a fixative solution containing 4% paraformaldehyde in 0.1 M phosphate buffer (PB, pH = 7.4). After that, the fixed brain was gently removed from the skull and stored at 4 °C in the fixative solution until processed further. Histological processing started by cutting 60-μm-thick coronal sections with a vibratome (Leica VT1200, Leica Microsystems, Wetzlar, Germany). Then, the brain sections were washed in 0.1 M PB, mounted onto microscopic slides and air dried. To identify brain sections containing fluorescent marks of DiI corresponding to the probe track, the slides were examined under a light microscope (Leica DM2500, Leica Microsystems) equipped with a fluorescence LED illumination source (SFL4000, Leica; Zeiss, Oberkochen, Germany in case anesthetized rats) and with a digital camera (DP70 or DP73, Olympus, Tokyo, Japan). Relevant sections from the anesthetized mouse experiments were processed for cresyl violet (Nissl) staining, dehydrated in xylene and coverslipped with DePex (SERVA Electrophoresis, Heidelberg, Germany); sections from the anesthetized rat experiments were stained for choline-acetyltransferase (ChAT; mouse monoclonal anti-ChAT (Umbriaco et al., 1994)) to visualize the MS; no staining for sections from the awake mouse experiments was performed. Finally, to verify the recording location based on the stereotaxic mouse brain atlas (Paxinos et al., 2001), sections containing the track of the silicon probe were photographed under the microscope.

#### Data analysis

All signal processing codes were implemented in MATLAB 2016a and 2020a (Mathworks, Natick, US).

#### Hippocampal state detection

Pyramidal layer was detected from linear silicon probe recordings based on the documented phase reversal of hippocampal theta oscillation below the pyramidal layer and verified based on histological reconstruction (Figure S2) (Buzsáki, 2006; Buzsaki et al., 1986; Green et al., 1960). Theta detection was performed on a single channel from the stratum radiatum (400 μm below the detected pyramidal layer in rats and 250 μm below the pyramidal layer in mice), since theta amplitude was largest in this layer, allowing the most robust detection (Buzsáki, 2002, 2006). In one chronically implanted mouse, a Buzsaki-type probe was used in the hippocampus; in this case, the deepest channel was used for LFP analysis, which appeared to be below the pyramidal layer based on theta phase reversion, thus providing phase values consistent with other recordings. The appropriate theta and delta frequency band boundaries were defined separately for the three different rodent models based on the Fourier spectra of the recordings (Figure 1B). Our observations confirmed known spectral differences between urethane-anesthetized and awake rodents (frequency boundaries in anesthetized rats, delta, 0.5-2.5 Hz; theta, 3-8 Hz; in anesthetized mice, delta, 0.5-2 Hz, theta, 2-8 Hz; in awake mice, delta, 0.5-4 Hz, theta, 5-10; in optogenetic experiments, frequency boundaries were optimized for each mouse individually). Raw LFP traces were resampled at 1 kHz and bandpass filtered in the theta and delta bands using the built-in finite impulse response filter (fir1.m) with zero-phase-lag filtering (built-in filtfilt.m function). The filtered traces were Hilbert-transformed and the instantaneous amplitude and phase values were calculated as the magnitude and phase of the complex Hilbert-transform. A simple artifact removal was performed on the theta/delta ratio by clamping values exceeding 10-fold difference. Next, the theta/delta amplitude ratio was smoothed with a moving average (window size, 5 seconds for anesthetized and 3 seconds for awake recordings; the difference in window size was due to the observed faster and more frequent state transitions in awake recordings). Theta segments were defined where the theta/delta ratio exceeded an empirically defined threshold, optimized for each dataset separately (1 for anesthetized mouse and rat recordings, 2 for awake mouse recordings and 1.5 in the optogenetic experiments); all other segments were defined as non-theta. Brief interruptions of theta or non-theta segments were not considered state switches (<5 seconds in anesthetized and μ3 seconds in awake recordings; the difference was due to the observed faster and more frequent state transitions in awake recordings). In some optogenetic tagging experiments (n = 8 recordings), only a part of the recording was analyzed due to electrical artifacts in other segments.

#### Spike sorting of septal neurons

The mouse MS recordings were fed to the Kilosort software (Pachitariu et al., 2016). Clustering was initialized with the desired number of clusters set to twice the number of channels. The Kilosort output was curated manually in the Phy template-gui graphical user interface module. We examined any potential violations of the refractory periods, slow electrode drifts, spatial distributions of action potential (AP) energies among neighboring channels, AP shapes and amplitudes and the principal components (PC) of the AP shapes. In case of noisy autocorrelograms and multiple subclusters in the PC space, we considered manual splitting of the automatically assigned cluster. The high similarity of clusters as well as lack of co-firing (‘mutual refractoriness’) in crosscorrelograms could imply that two automatically determined clusters belonged to the same neuron, in which case the clusters were merged. Finally, we calculated objective cluster quality measures based on the interspike interval histograms and cluster isolation distances, to verify putative single neuron clusters. Kilosort execution time was reduced by running on an Nvidia Geforce GTX 1080 graphic card with the Matlab Parallel Computing Toolbox. The anaesthetized rat recordings were clustered automatically in KlustaKwik (Rossant et al., 2016) (available at http://github.com/klusta-team) and curated manually based on auto- and crosscorrelograms similar to the mouse recordings, as described previously (Bartho et al., 2004).

### Septal cell rhythmicity

#### Rhythmicity indices

Theta- and delta-rhythmicity of MS neurons was judged based on rhythmicity indices derived from autocorrelograms. First, autocorrelation of the spike train was computed in a ±3 seconds window at 1 ms resolution; the central peak corresponding to the total spike number was removed and the autocorrelogram was smoothed by a 20-ms moving average and normalized to an integral of 1. Next, theta and delta peaks were determined between time lags corresponding to theta and delta frequency bands. ‘Peak values’ were averaged from the autocorrelograms in the ±20 ms neighborhoods of the peaks. ‘Baseline values’ were averaged from the ±20 ms neighborhood around the lags corresponding to half times and one and a half times the peak location (assumed troughs). Finally, the difference between the peak and baseline values was normalized to the larger of the two, yielding a Theta Index and a Delta Index between −1 and 1:

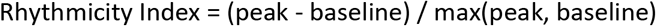

Rhythmicity indices were calculated separately for hippocampal theta and non-theta segments. Cells with insufficient number of spikes (<2 spikes/autocorrelogram bin on average) were excluded from later analysis.

To assess statistical significance of rhythmicity in the theta and delta frequency bands, we simulated a dataset of neurons exhibiting spiking generated by a Poisson-process. We determined the λ parameter of the Poisson-process individually for each simulated cell so that they matched the firing rate of an actual MS neuron. Rhythmicity Index was calculated for the simulated neurons using the same procedure as for the biological recordings, then Rhythmicity Index thresholds were determined corresponding to p < 0.05 significance levels based on this bootstrap null distribution.

#### Theta-burst Index

To differentiate between tonically active and putative pacemaker neurons, a Theta-burst Index (TBI) was introduced, quantifying the characteristic long theta-bursts putative pacemakers fire during individual theta cycles (Borhegyi et al., 2004; Varga et al., 2008). Average autocorrelograms between lags corresponding to a pre-defined window spanning the intraburst interspike intervals (anesthetized rats, 20-40ms; anesthetized mice, 20-50 ms and 20-40 ms in optogenetic tagging experiments, see below; freely moving mice, 10-30 ms) were normalized to the average autocorrelation, similar to the Rhythmicity Indices, providing a TBI between −1 and 1. The above windows were determined based on variations of theta frequencies across datasets. Using a traditional 20-50 ms window for all datasets altered the classification of only a few neurons and did not change the main results. Tonically active neurons were defined as those having a TBI <0.

#### Analysis of phase-coupling

Selected LFP channels were filtered in the theta and delta frequency bands, as described above. The filtered signals were Hilbert-transformed and the instantaneous phase values of the analytical signal were computed. Spike phases were determined by taking the instantaneous phase values corresponding to the spike times. To quantify phase-locking, mean phase and mean resultant length (measure of phase-locking strength) were calculated as the angle and magnitude of the first trigonometric moment of spike phases. Phase value distributions for individual neurons were visualized as phase histograms, and distributions over groups of neurons were visualized as polar plots, in which each neuron was represented by a vector pointing to its mean phase, with a length corresponding to its mean resultant length and color corresponding to its firing rate in greyscale.

#### Crosscorrelation

Crosscorrelations of simultaneously recorded pairs of MS neurons were calculated in ±3 second windows at 1 ms resolution, smoothed by a 20-ms moving average and normalized to an integral of 1. Crosscorrelograms were aligned to their peaks between ±1.5 seconds and averaged both within and across rhythmicity groups.

#### Pacemaker synchronization

To test how putative pacemakers synchronize during theta segments, we calculated a number of parameters and performed comparisons during theta vs. non-theta episodes. (i) Firing rates were calculated as the ratio of spike number and total segment length. (ii) Autocorrelation peak values and rhythmicity frequencies were estimated from the autocorrelograms, smoothed by a 20-ms moving average (see above). (iii) Theta-bursts were defined by interspike intervals falling in the theta-burst windows defined for the TBI (see above) and average intraburst interspike intervals were computed. (iv) To estimate theta cycle skipping, an event vector was built from first spikes of bursts and single spikes. Theta cycle length was calculated for each neuron based on its rhythmicity frequency (see above), and an edge vector was defined with uniform spacing. Events were binned by the edge vector and skipping was quantified as the ratio of empty to all cycles. Although the edge vector was only based on neuronal rhythmicity and thus could not account for individual cycle variabilities, this approach likely provides a good approximation of theta cycle skipping. Since we aimed to define this metric for both theta and non-theta segments, we had to move away from LFP-based cycle definitions for this analysis. (v) Difference in rhythmicity frequency was calculated between concurrently recorded pairs of putative pacemaker neurons and normalized to the greater of the two frequencies.

#### Data analysis of the optogenetic tagging experiments

Laser stimulation segments (first and last 2 minutes) were removed from the hippocampal recordings. Hippocampal state detection was performed on a single channel from the stratum radiatum (~250 μm below the pyramidal layer). The LFP signal was resampled at 1 kHz. Theta and delta frequency band boundaries were determined based on spectral components (theta: 1.5-4 Hz; 1-4 and 1.5-6 Hz in 1-1 mice; delta: 0.5-1.5 Hz, 0.5-1 Hz in one mouse). Theta/ delta amplitude ratio was smoothed with a 5 second window and a 1.5 threshold (1 in one mouse) was applied for theta detection. Additionally, theta amplitude had to surpass the median value of the theta-filtered signal. Less than 5-second-long epochs were not considered as state transitions. In 8 recordings, only a part of the recording was analyzed due to electrical artifacts in other segments.

MS units were sorted in Kilosort as described above. MS neurons that had sufficient number of spikes (>500 in total and median interspike interval <500 ms) were further analyzed. Note that these spike number criteria were less stringent compared to other datasets (see above) due to generally shorter recordings. Significant photoactivation (p < 0.01) was determined using the stimulus-associated spike latency test (SALT) based on specific spike timing after light flashes (Kvitsiani et al., 2013). After removing the phototagging segments from the recordings, rhythmicity indices and TBI (with [20,40] ms window) were computed as described above.

#### MS pacemaker neuron model

The model was implemented in NEURON (Carnevale and Hines, 2006). We used a fixed time step of 0.025 ms. The neuron network parameters were defined and the simulation results were processed by Matlab scripts.

We based our model on a fast spiking single compartment neuron model by Golomb et al. (Golomb et al., 2007). This choice was motivated by the model’s simplicity and its desirable firing properties, capable of high-frequency tonic firing but also rhythmic bursting, characteristic of MS pacemaker neurons. This model included a transient sodium channel (nas) that is necessary for action potential generation, the delayed rectifier potassium channel (kdr) for repolarization during fast spiking and the slowly inactivating d-type potassium channel (kd) for bursting behavior through controlling the duration of the after-hyperpolarization. We added the HCN-channel (hcn) responsible for the hyperpolarization-activated H-current, which was reported to be responsible for sag responses in MS neurons (Kocsis and Li, 2004; Morris et al., 2004; Sotty et al., 2003; Varga et al., 2008; Xu et al., 2004). We could reproduce this sag response by including the H-current (Fig.5C); we note, however, that the presence of the H-current was not a prerequisite of rhythmic bursting in the present model. Further electrophysiology data and modeling should clarify the exact channel properties of MS pacemaker neurons. Nevertheless, the present model was capable of reproducing the characteristic rhythmic bursting of MS pacemaker neurons in the range of biologically plausible rhythmicity frequencies upon current injection, serving the modeling purposes of this study.

Potential change across the cell membrane was described by the following equation.

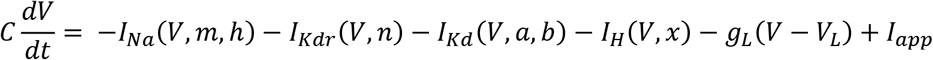

where C = 1μF/cm^2^ is the membrane capacitance, g_L_ = 0.1 mS/cm^2^ is the passive conductance, V_L_ = −60 mV is the equilibrium potential for leakage and I_app_ is the injected current in μA/cm^2^. The lower gL and higher VL compared to the Golomb-model (gL = 0.25 mS/cm^2^, VL = −70 mV) were introduced to facilitate the characteristic sag response mediated by the HCN channels. The total surface of the neuron was 5000 μm^2^.

The applied point source input current amplitude varied between 40 and 80 pA in the network simulations (below).

Transient Na current *I_Na_* was governed by the following equations.

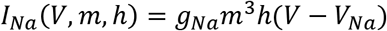

where the maximal conductance of the channels was *g_Na_* = 112.5 mS/cm^2^, the reversal potential of sodium ion was *V_Na_* = 50 mV and the dynamics of the three activation gates *m* and one inactivation gate *h* were given by

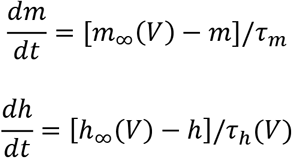

The time constant of the m gate was *τ_m_* = 0.01 ms and the voltage-dependent steady-state activation and inactivation variables *m*_∞_, *h*_∞_ and time constant of the *h* gate τ_*h*_ were

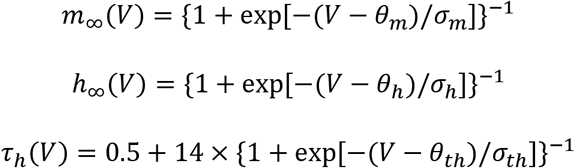

where *θ_m_* = −24 mV, *σ_m_* = 11.5 mV, *θ_h_* = −58.3 mV, *σ_m_* = −6.7 mV, *θ_th_* = −60 mV and *σ_th_* = −12 mV.

The delayed rectifier K current *I_Kdr_* was governed by

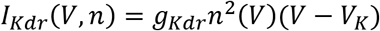

where the maximal conductance of the channels was *g_kdr_* = 225 mS/cm^2^, the reversal potential of potassium ions was *V_k_* = −90 mV and the dynamics of the two activation gates *n* were defined by

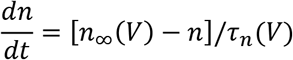

The voltage-dependent steady-state activation variable *n_∞_* and time constant of the *n* gates *τ_n_* were

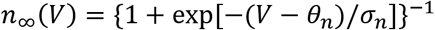

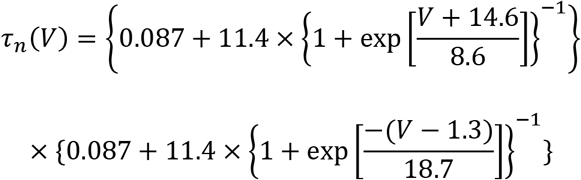

where *θ_n_* = −12.4 mV, *σ_n_* = 6.8 mV.

The slowly inactivating d-type K current *I_Kd_* was governed by

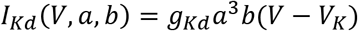

where the maximal conductance of the channels was *g_Kd_* = 1.8 mS/cm^2^ and the dynamics of the three activation gates *a* and one inactivation gate *b* were defined by

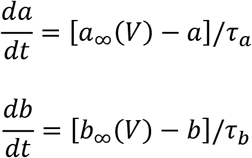

The time constant of the activation gate was *τ_a_* = 2 ms and that of the inactivation gate was *τ_b_* = 120 ms.

The voltage-dependent steady-state activation and inactivation variables *a_∞_* and *b_∞_* were

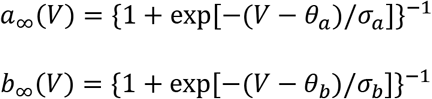

where *θ_a_* = −50 mV, *σ_a_* = 20 mV, *θ_b_* = −60 mV, *σ_b_* = −6 mV.

We introduced smaller *τ_b_* compared to the Golomb-model (where *τ_b_* was 150 ms), which resulted in faster rhythmicity that was within the delta-theta frequency range. We used a *θ_b_* of −60 mV that enabled the characteristic sag response by the H-current.

To define the HCN channel, we started from a generic HCN model by Kali and Zemankovics (Káli and Zemankovics, 2012) which used an alternative parameterization of the Hodgkin-Huxley type model as described by Borg-Graham (Borg-Graham, 1999). Next, parameters of our model were tuned to match the dynamical properties of the H-current in MS neurons (Migliore and Migliore, 2012; Xu et al., 2004). The final *IH* current was defined by

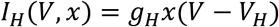

where the maximal conductance of the channels was *g_H_* = 0.1 mS/cm^2^ and the reversal potential of the cations was *V_x_* = −30 mV. The open probability of the activation gate *x* evolved as

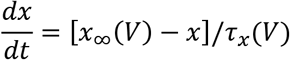

where the voltage-dependent steady-state activation variable *x_∞_* and time constant *τ_x_* were

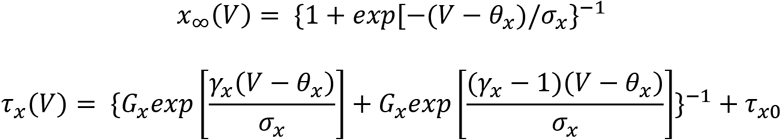

where rate coefficient was *G_x_* = 1.2353 1/ms, asymmetry parameter was *γ_x_* = 0.81, half activation was *θ_x_* = −98 mV, *σ_x_* = −6.73 mV, *τ_x0_* = 130 ms.

#### Modeling the MS pacemaker network

The neuron network parameters were defined, and the simulation results were processed in Matlab. In our network simulations, 20 inhibitory pacemaker neurons were connected at an average connection rate (CR) varied between 10 and 70% (randomly chosen connections were deleted from a fully connected graph). Inhibitory synapses were implemented with the built-in Netcon() and ExpSyn() point processes of the NEURON environment. Netcon() implements a presynaptic source object with threshold, delay and weight parameters. The threshold parameter was fixed at 0 mV and the delay was fixed at 1 ms. Expsyn() models the postsynaptic effect with a discontinuous change in conductance after each incoming spike event (which is initiated by Netcon() when the presynaptic membrane potential crosses the spike detection threshold), followed by an exponential decay with time constant *τ*:

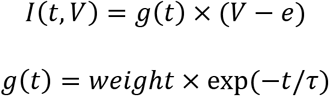

where *I* (in nA) was the current transmitted to the postsynaptic cell, *g* (in μS) was the actual conductance, *weight* (in μS) was the maximal conductance or synaptic strength, *e* (in mV) was the reversal potential, which was set to −70 mV and τ (in ms) was the exponential decay factor, which was set to 5 ms with 10% variance.

The network simulations were implemented by NEURONS’s built-in IClamp() function, which represented an electrode delivering current pulses of a given duration, amplitude and delay. Every neuron received three electrodes at their somata, delivering a sequence of three current pulses implementing weak, strong (1.4-fold), and weak tonic excitation in this order. The extent of increase in excitation strength was based on the average firing rate increase of MS glutamatergic neurons, the main excitatory input to MS GABAergic cells (Hajszan et al., 2004), determined by our optogenetic tagging experiments. During each simulation, synaptic and stimulation parameters were drawn from normal distributions with given mean and variance.

We modeled the theoretical output signal of the simulated pacemaker network as a proxy for the septal input to the hippocampus. Spike trains (i.e. the events when the simulated membrane potential reached the 0 mV spike generation threshold) were convolved with a 50 ms Gaussian window and averaged across all neurons. State detection was executed similarly as described for real data above with the following parameters: resampling rate, 1 kHz; theta band 4-6 Hz; delta band, 0.5-4 Hz; theta-ratio was smoothed and short segments (<0.5 s) handled as above; theta-delta ratio threshold, 2.

#### Exploring the parameter space of the pacemaker network model

To study the network dynamics and its dependence on key parameters, we performed a series of simulations. In each run, we simulated a 5-second-long baseline period, followed by a 10-second-long ‘theta’ period when a stronger excitatory current was injected to all pacemaker neurons (1.4-fold of baseline), followed by a 5-second-long ‘non-theta’ period when tonic excitation was reverted to baseline. We systematically varied the connection rate (10-70% in increments of 10%), mean absolute synaptic strength (0.05, 0.1, 0.5, 1, 2, 3 nS), mean (40-80 pA in increments of 10 pA) and variance (5-25% in increments of 5%) of baseline stimulation strength. We performed two sets of simulations. First, the variance of baseline stimulation was fixed at 10% and all combinations of the other parameters were tested. Second, the connection rate was fixed at 50% and the possible combinations of the remaining parameters were tested. We repeated each simulation for a given parameter arrangement 10 times by redrawing parameters from the same distributions, performing 2100 and 1500 runs in the first and second set of simulations, respectively. We evaluated the simulated models by calculating a simple synchronization score, which quantified the ratio of the total simulation time the network spent in the expected state based on the state detection described above.

#### Model network synchronization

Based on the exploration of the network model parameter space, we fixed mean baseline excitation at 60 pA with 10% variance, connectivity rate at 50% and mean synaptic strength at 0.5 nS. We performed a series of simulations (n = 60) of 60-second-long segments, with 20 seconds of baseline, followed by 20 seconds of increased excitation, followed by 20 seconds of returning to baseline excitation levels to enable large scale analysis of pacemaker synchronization (Figure 5F, Figure S19). Rhythmicity indices, TBI (theta-burst interspike interval window, 20-40 ms), significant modulations, phase preference (with respect to the theoretical population output) and pacemaker synchronization were computed as described for the biological data.

## Code availability

MATLAB and NEURON codes generated for this study are available at https://github.com/hangyabalazs/ms_sync_analysis

## Acknowledgement

We thank Katalin Lengyel for her help with histology. This work was supported by the ‘Lendület’ Program of the Hungarian Academy of Sciences (LP2015-2/2015), NKFIH KH125294, NKFIH K135561 and the European Research Council Starting Grant no. 715043 to BH; NKFIH K119650 to PB; National Brain Research Program 1.2.1-NKP-2017-00002 to PB, RF and IU; NKFIH PD124175 and PD134196 to RF; NKFIH TUDFO/51757-1/2019-ITM to IU. BH and SK were supported by the Ministry of Innovation and the National Research, Development and Innovation Office within the framework of the Artificial Intelligence National Laboratory Programme. BK was supported by the ÚNKP-20-3 New National Excellence Program of the Ministry for Innovation and Technology from the source of the National Research, Development and Innovation Fund. SMB was part of the Generalitat Valenciana Postdoctoral Fellowship Program (APOSTD/2019/003).

## Author contributions

BH developed the idea and conceptualized the manuscript. SMB, RF, AD, BH and PB performed the experiments, supervised by TFF, IU, VV and BH. BK, SMB and PB performed data analysis. BK and KS generated the figures. BK performed the modeling, supervised by SK. BH wrote the manuscript with input from all authors.

## References

Alonso, L.M., and Marder, E. (2019). Visualization of currents in neural models with similar behavior and different conductance densities. Elife 8.

Apartis, E., Poindessous-Jazat, F.R., Lamour, Y.A., and Bassant, M.H. (1998). Loss of rhythmically bursting neurons in rat medial septum following selective lesion of septohippocampal cholinergic system. J. Neurophysiol. 79, 1633–1642.

Barrenechea, C., Pedemonte, M., Nu༞z, A., and García-Austt, E. (1995). In vivo intracellular recordings of medial septal and diagonal band of Broca neurons: relationships with theta rhythm. Exp. Brain Res. 103, 31–40.

Bartho, P., Hirase, H., Monconduit, L., Zugaro, M., Harris, K.D., and Buzsaki, G. (2004). Characterization of neocortical principal cells and interneurons by network interactions and extracellular features. J. Neurophysiol. 92, 600–608.

Bland, B.H., Trepel, C., Oddie, S.D., and Kirk, I.J. (1996). Intraseptal microinfusion of muscimol: Effects on hippocampal formation theta field activity and phasic Theta-ON cell discharges. Exp. Neurol. 138, 286–297.

Bland, B.H., Declerck, S., Jackson, J., Glasgow, S., and Oddie, S. (2007). Septohippocampal properties of N-methyl-D-aspartate-induced theta-band oscillation and synchrony. Synapse 61, 185–197.

Borg-Graham, L.J. (1999). Models of Cortical Circuits. In Interpretations of Data and Mechanisms for Hippocampal Pyramidal Cell Models, (Springer, Boston, MA), pp. 19–138.

Borhegyi, Z., Varga, V., Szilagyi, N., Fabo, D., and Freund, T.F. (2004). Phase segregation of medial septal GABAergic neurons during hippocampal theta activity. J. Neurosci. 24, 8470–8479.

Brandon, M.P., Bogaard, A.R., Schultheiss, N.W., and Hasselmo, M.E. (2013). Segregation of cortical head direction cell assemblies on alternating theta cycles. Nat. Neurosci. 16, 739–748.

Brazhnik, E.S., and Fox, S.E. (1997). Intracellular recordings from medial septal neurons during hippocampal theta rhythm. Exp. Brain Res. 114, 442–453.

Brazhnik, E.S., and Vinogradova, O.S. [The effect of disconnecting the hippocampus from the septum on the activity of septal neurons]. Zh. Vyssh. Nerv. Deiat. Im. I P Pavlova 25, 1044–1052.

Buzsaki, G., Czopf, J., Kondakor, I., and Kellenyi, L. (1986). Laminar distribution of hippocampal rhythmic slow activity (RSA) in the behaving rat: current-source density analysis, effects of urethane and atropine. Brain Res. 365, 125–137.

Buzsáki, G. (2002). Theta oscillations in the hippocampus. Neuron 33, 325–340.

Buzsáki, G. (2006). Rhythms of the Brain (Oxford University Press).

Buzsáki, G., and Moser, E.I. (2013). Memory, navigation and theta rhythm in the hippocampal-entorhinal system. Nat. Neurosci. 16, 130–138.

Carnevale, N.T., and Hines, M.L. (2006). The NEURON Book (Cambridge: Cambridge University Press).

Cutsuridis, V., and Poirazi, P. (2015). A computational study on how theta modulated inhibition can account for the long temporal windows in the entorhinal-hippocampal loop. Neurobiol. Learn. Mem. 120, 69–83.

Dannenberg, H., Pabst, M., Braganza, O., Schoch, S., Niediek, J., Bayraktar, M., Mormann, F., and Beck, H. (2015). Synergy of direct and indirect cholinergic septo-hippocampal pathways coordinates firing in hippocampal networks. J. Neurosci. 35, 8394–8410.

Denham, M.J., and Borisyuk, R.M. (2000). A model of theta rhythm production in the septal-hippocampal system and its modulation by ascending brain stem pathways. Hippocampus 10, 698–716.

Destrade, C., and Ott, T. (1982). Is a retrosplenial (cingulate) pathway involved in the mediation of high frequency hippocampal rhythmical slow activity (theta)? Brain Res. 252, 29–37.

DiCarlo, J.J., Lane, J.W., Hsiao, S.S., and Johnson, K.O. (1996). Marking microelectrode penetrations with fluorescent dyes. J. Neurosci. Methods 64, 75–81.

Duque, A., Balatoni, B., Detari, L., and Zaborszky, L. (2000). EEG correlation of the discharge properties of identified neurons in the basal forebrain. J. Neurophysiol. 84, 1627–1635.

Equihua, G.G.V., and Ramirez, J.P. (2018). Synchronization of Hindmarsh-Rose Synchronization of Hindmarsh-Rose Synchronization of Hindmarsh-Rose neurons Huygens-like coupling Synchronization of Hindmarsh-Rose neurons Huygens-like coupling Huygens-like coupling. IFAC-PapersOnLine 51, 186–191.

Fiáth, R., Raducanu, B.C., Musa, S., Andrei, A., Lopez, C.M., van Hoof, C., Ruther, P., Aarts, A., Horváth, D., and Ulbert, I. (2018). A silicon-based neural probe with densely-packed low-impedance titanium nitride microelectrodes for ultrahigh-resolution in vivo recordings. Biosens. Bioelectron. 106, 86–92.

Fiáth, R., Márton, A.L., Mátyás, F., Pinke, D., Márton, G., Tóth, K., and Ulbert, I. (2019). Slow insertion of silicon probes improves the quality of acute neuronal recordings. Sci. Rep. 9, 111.

Ford, R.D., Colom, L. V., and Bland, B.H. (1989). The classification of medial septum-diagonal band cells as σ-on or σ-off in relation to hippocampal EEG states. Brain Res. 493, 269–282.

Freund, T.F. (1989). GABAergic septohippocampal neurons contain parvalbumin. Brain Res. 478, 375–381.

Freund, T.F., and Antal, M. (1988). GABA-containing neurons in the septum control inhibitory interneurons in the hippocampus. Nature 336, 403–405.

Freund, T.F., and Gulyás, A.I. (1991). GABAergic interneurons containing calbindin D28K or somatostatin are major targets of GABAergic basal forebrain afferents in the rat neocortex. J. Comp. Neurol. 314, 187–199.

Fuhrmann, F., Justus, D., Sosulina, L., Kaneko, H., Beutel, T., Friedrichs, D., Schoch, S., Schwarz, M.K., Fuhrmann, M., and Remy, S. (2015). Locomotion, Theta Oscillations, and the Speed-Correlated Firing of Hippocampal Neurons Are Controlled by a Medial Septal Glutamatergic Circuit. Neuron 86, 1253–1264.

Golomb, D., Donner, K., Shacham, L., Shlosberg, D., Amitai, Y., and Hansel, D. (2007). Mechanisms of Firing Patterns in Fast-Spiking Cortical Interneurons. PLoS Comput. Biol. 3, e156.

Green, J.D., and Arduini, A.A. (1954). Hippocampal electrical activity in arousal. J. Neurophysiol. 17, 533–557.

Green, J.D., Maxwell, D.S., Schindler, W.J., and Stumpf, C. (1960). Rabbit Eeg “Theta” Rhythm: Its Anatomical Source and Relation To Activity in Single Neurons. J. Neurophysiol. 23, 403–420.

Gritti, I., Henny, P., Galloni, F., Mainville, L., Mariotti, M., and Jones, B.E. (2006). Stereological estimates of the basal forebrain cell population in the rat, including neurons containing choline acetyltransferase, glutamic acid decarboxylase or phosphate-activated glutaminase and colocalizing vesicular glutamate transporters. Neuroscience 143, 1051–1064.

Hajós, M., Hoffmann, W.E., Orbán, G., Kiss, T., and Érdi, P. (2004). Modulation of septo-hippocampal θ activity by GABAA receptors: An experimental and computational approach. Neuroscience 126, 599–610.

Hajszan, T., Alreja, M., and Leranth, C. (2004). Intrinsic vesicular glutamate transporter 2- immunoreactive input to septohippocampal parvalbumin-containing neurons: Novel glutamatergic local circuit cells. Hippocampus 14, 499–509.

Hangya, B., Borhegyi, Z., Szilágyi, N., Freund, T.F., and Varga, V. (2009). GABAergic neurons of the medial septum lead the hippocampal network during theta activity. J. Neurosci. 29, 8094–8102.

Hassani, O.K., Lee, M.G., Henny, P., and Jones, B.E. (2009). Discharge profiles of identified GABAergic in comparison to cholinergic and putative glutamatergic basal forebrain neurons across the sleep-wake cycle. J. Neurosci. 29, 11828–11840.

Hasselmo, M.E. (2014). Neuronal rebound spiking, resonance frequency and theta cycle skipping may contribute to grid cell firing in medial entorhinal cortex. Philos. Trans. R. Soc. B Biol. Sci. 369, 20120523.

Huh, C.Y.L., Goutagny, R., and Williams, S. (2010). Glutamatergic neurons of the mouse medial septum and diagonal band of broca synaptically drive hippocampal pyramidal cells: Relevance for hippocampal theta rhythm. J. Neurosci. 30, 15951–15961.

Huygens, C. (1673). Horologium Oscillatorium: sive de motu pendulorum ad horologia aptato demonstrationes geometricae (Paris: F. Muguet).

Inokawa, H., Yamada, H., Matsumoto, N., Muranishi, M., and Kimura, M. (2010). Juxtacellular labeling of tonically active neurons and phasically active neurons in the rat striatum. Neuroscience 168, 395–404.

Jeffery, K.J., Donnett, J.G., and OʼKeefe, J. (1995). Medial septal control of theta-correlated unit firing in the entorhinal cortex of awake rats. Neuroreport 6, 2166–2170.

Joshi, A., Salib, M., Viney, T.J., Dupret, D., and Somogyi, P. (2017). Behavior-Dependent Activity and Synaptic Organization of Septo-hippocampal GABAergic Neurons Selectively Targeting the Hippocampal CA3 Area. Neuron 96, 1342–1357.e5.

Jung, R., and Kornmüller, A.E. (1938). Eine Methodik der Ableitung Iokalisierter Potentialschwankungen aus subcorticalen Hirngebieten. Arch. Psychiatr. Nervenkr. 109, 1–30.

Justus, D., Dalügge, D., Bothe, S., Fuhrmann, F., Hannes, C., Kaneko, H., Friedrichs, D., Sosulina, L., Schwarz, I., Elliott, D.A., et al. (2017). Glutamatergic synaptic integration of locomotion speed via septoentorhinal projections. Nat. Neurosci. 20, 16–19.

Kahana, M.J., Sekuler, R., Caplan, J.B., Kirschen, M., and Madsen, J.R. (1999). Human theta oscillations exhibit task dependence during virtual maze navigation. Nature 399, 781–784.

Káli, S., and Zemankovics, R. (2012). The effect of dendritic voltage-gated conductances on the neuronal impedance: a quantitative model. J. Comput. Neurosci. 33, 257–284.

Katona, L., Micklem, B., Borhegyi, Z., Swiejkowski, D.A., Valenti, O., Viney, T.J., Kotzadimitriou, D., Klausberger, T., and Somogyi, P. (2017). Behavior-dependent activity patterns of GABAergic long-range projecting neurons in the rat hippocampus. Hippocampus 27, 359–377.

Kay, K., Chung, J.E., Sosa, M., Schor, J.S., Karlsson, M.P., Larkin, M.C., Liu, D.F., and Frank, L.M. (2020). Constant Sub-second Cycling between Representations of Possible Futures in the Hippocampus. Cell 180, 552–567.e25.

King, C., Recce, M., and O’Keefe, J. (1998). The rhythmicity of cells of the medial septum/diagonal band of Broca in the awake freely moving rat: relationships with behaviour and hippocampal theta. Eur. J. Neurosci. 10, 464–477.

Kiss, J., Patel, A.J., Baimbridge, K.G., and Freund, T.F. (1990). Topographical localization of neurons containing parvalbumin and choline acetyltransferase in the medial septum-diagonal band region of the rat. Neuroscience 36, 61–72.

Klausberger, T., and Somogyi, P. (2008). Neuronal diversity and temporal dynamics: the unity of hippocampal circuit operations. Science 321, 53–57.

Kocsis, B., and Li, S. (2004). In vivo contribution of h-channels in the septal pacemaker to theta rhythm generation. Eur. J. Neurosci. 20, 2149–2158.

Korteweg, D. (1906). Les horloges sympathiques de Huygens. Arch. Neerl. Des Sci. Exactes Nat. 11, 273–296.

Kramis, R., Vanderwolf, C.H., and Bland, B.H. (1975). Two types of hippocampal rhythmical slow activity in both the rabbit and the rat: Relations to behavior and effects of atropine, diethyl ether, urethane, and pentobarbital. Exp. Neurol. 49, 58–85.

Kvitsiani, D., Ranade, S., Hangya, B., Taniguchi, H., Huang, J.Z., and Kepecs, A. (2013). Distinct behavioural and network correlates of two interneuron types in prefrontal cortex. Nature 498, 363–366.

Laszlovszky, T., Schlingloff, D., Hegedüs, P., Freund, T.F., Gulyás, A., Kepecs, A., and Hangya, B. (2020). Distinct synchronization, cortical coupling and behavioral function of two basal forebrain cholinergic neuron types. Nat. Neurosci. 23, 992–1003.

Lee, M.G., Chrobak, J.J., Sik, A., Wiley, R.G., and Buzsáki, G. (1994). Hippocampal theta activity following selective lesion of the septal cholinergic systeM. Neuroscience 62, 1033–1047.

Lee, M.G., Hassani, O.K., Alonso, A., and Jones, B.E. (2005). Cholinergic basal forebrain neurons burst with theta during waking and paradoxical sleep. J. Neurosci. 25, 4365–4369.

Leranth, C., and Frotscher, M. (1989). Organization of the septal region in the rat brain: Cholinergic‐GABAergic interconnections and the termination of hippocampo‐septal fibers. J. Comp. Neurol. 289, 304–314.

Leung, L.S., and Shen, B. (2004). Glutamatergic synaptic transmission participates in generating the hippocampal EEG. Hippocampus 14, 510–525.

Li, S., Topchiy, I., and Kocsis, B. (2007). The effect of atropine administered in the medial septum or hippocampus on high- and low-frequency theta rhythms in the hippocampus of urethane anesthetized rats. Synapse 61, 412–419.

Lima, S.Q., Hromádka, T., Znamenskiy, P., and Zador, A.M. (2009). PINP: a new method of tagging neuronal populations for identification during in vivo electrophysiological recording. PLoS One 4, e6099.

Losonczy, A., Zemelman, B. V, Vaziri, A., and Magee, J.C. (2010). Network mechanisms of theta related neuronal activity in hippocampal CA1 pyramidal neurons. Nat. Neurosci. 13, 967–972.

Lu, C.B., Ouyang, G., Henderson, Z., and Li, X. (2011). Induction of theta-frequency oscillations in the rat medial septal diagonal band slice by metabotropic glutamate receptor agonists. Neuroscience 177, 1–11.

Mamad, O., McNamara, H.M., Reilly, R.B., and Tsanov, M. (2015). Medial septum regulates the hippocampal spatial representation. Front. Behav. Neurosci. 9, 1–16.

Manseau, F., Danik, M., and Williams, S. (2005). A functional glutamatergic neurone network in the medial septum and diagonal band area. J. Physiol. 566, 865–884.

Manseau, F., Goutagny, R., Danik, M., and Williams, S. (2008). The hippocamposeptal pathway generates rhythmic firing of GABAergic neurons in the medial septum and diagonal bands: an investigation using a complete septohippocampal preparation in vitro. J. Neurosci. 28, 4096–4107.

McNamara, C.G., Tejero-Cantero, Á., Trouche, S., Campo-Urriza, N., and Dupret, D. (2014). Dopaminergic neurons promote hippocampal reactivation and spatial memory persistence. Nat. Neurosci. 17, 1658–1660.

Migliore, M., and Migliore, R. (2012). Know your current I(h): interaction with a shunting current explains the puzzling effects of its pharmacological or pathological modulations. PLoS One 7, e36867.

Mikulovic, S., Restrepo, C.E., Siwani, S., Bauer, P., Pupe, S., Tort, A.B.L.L., Kullander, K., and Leão, R.N. (2018). Ventral hippocampal OLM cells control type 2 theta oscillations and response to predator odor. Nat. Commun. 9, 3638.

Moldestad, O., Karlsen, P., Molden, S., and Storm, J.F. (2009). Tracheotomy improves experiment success rate in mice during urethane anesthesia and stereotaxic surgery. J. Neurosci. Methods 176, 57–62.

Moore, J.D., Deschênes, M., Furuta, T., Huber, D., Smear, M.C., Demers, M., and Kleinfeld, D. (2013). Hierarchy of orofacial rhythms revealed through whisking and breathing. Nature 497, 205–210.

Morris, N.P., Fyffe, R.E.W.W., and Robertson, B. (2004). Characterisation of hyperpolarization-activated currents (Ih) in the medial septum/diagonal band complex in the mouse. Brain Res. 1006, 74–86.

Müller, C., and Remy, S. (2018). Septo–hippocampal interaction. Cell Tissue Res. 373, 565–575.

Mysin, I.E., Kitchigina, V.F., and Kazanovich, Y. (2015). Modeling synchronous theta activity in the medial septum: Key role of local communications between different cell populations. J. Comput. Neurosci. 39, 1–16.

Nicola, W., and Clopath, C. (2019). A diversity of interneurons and Hebbian plasticity facilitate rapid compressible learning in the hippocampus. Nat. Neurosci. 22, 1168–1181.

Oddie, S.D., and Bland, B.H. (1998). Hippocampal formation theta activity and movement selection. Neurosci. Biobehav. Rev. 22, 221–231.

Oddie, S.D., Stefanek, W., Kirk, I.J., and Bland, B.H. (1996). Intraseptal procaine abolishes hypothalamic stimulation-induced wheel-running and hippocampal theta field activity in rats. J. Neurosci. 16, 1948–1956.

Oliveira, H.M., and Melo, L. V. (2015). Huygens synchronization of two clocks. Sci. Rep. 5, 1–12.

Pachitariu, M., Steinmetz, N., Kadir, S., Carandini, M., and Harris, K.D. (2016). Kilosort: realtime spike-sorting for extracellular electrophysiology with hundreds of channels. BioRxiv 061481.

Paxinos, G., Franklin, K.B.J., and Franklin, K.B.J. (2001). The mouse brain in stereotaxic coordinates (Academic Press).

Payne, H.L., Lynch, G.F., and Aronov, D. (2020). Precise spatial representations in the hippocampus of a food-caching bird Main text 8. BioRxiv 2020.11.27.399444.

Petsche, H., Stumpf, C., and Gogolak, G. (1962). The significance of the rabbit’s septum as a relay station between the midbrain and the hippocampus I. The control of hippocampus arousal activity by the septum cells. Electroencephalogr. Clin. Neurophysiol. 14, 202–211.

Ramirez, J.P., Fey, R.H.B., Aihara, K., and Nijmeijer, H. (2014). An improved model for the classical Huygens’ experiment on synchronization of pendulum clocks. J. Sound Vib. 333, 7248–7266.

Ramirez, J.P., Olvera, L.A., Nijmeijer, H., and Alvarez, J. (2016). The sympathy of two pendulum clocks: Beyond Huygens’ observations. Sci. Rep. 6, 1–16.

Ramón y Cajal, S. (1909). Histologie du système nerveux de l’homme & des vertébrés. (Paris: Maloine).

Ranade, S., Hangya, B., and Kepecs, A. (2013). Multiple modes of phase locking between sniffing and whisking during active exploration. J. Neurosci. 33, 8250–8256.

Robinson, J., Manseau, F., Ducharme, G., Amilhon, B., Vigneault, E., El Mestikawy, S., and Williams, S. (2016). Optogenetic activation of septal glutamatergic neurons drive hippocampal theta rhythms. J. Neurosci. 36, 3016–3023.

Roland, J.J., Janke, K.L., Servatius, R.J., and Pang, K.C.H. (2014). GABAergic neurons in the medial septum-diagonal band of Broca (MSDB) are important for acquisition of the classically conditioned eyeblink response. Brain Struct. Funct. 219, 1231–1237.

Rossant, C., Kadir, S.N., Goodman, D.F.M., Schulman, J., Hunter, M.L.D., Saleem, A.B., Grosmark, A., Belluscio, M., Denfield, G.H., Ecker, A.S., et al. (2016). Spike sorting for large, dense electrode arrays. Nat. Neurosci. 19, 634–641.

Rutishauser, U., Ross, I.B., Mamelak, A.N., and Schuman, E.M. (2010). Human memory strength is predicted by theta-frequency phase-locking of single neurons. Nature 464, 903–907.

Semba, K., and Komisaruk, B.R. (1984). Neural substrates of two different rhythmical vibrissal movements in the rat. Neuroscience 12, 761–774.

Serafin, M., Williams, S., Khateb, A., Fort, P., and Mühlethaler, M. (1996). Rhythmic firing of medial septum non-cholinergic neurons. Neuroscience 75, 671–675.

Simon, A.P., Poindessous-Jazat, F., Dutar, P., Epelbaum, J., and Bassant, M.H. (2006). Firing properties of anatomically identified neurons in the medial septum of anesthetized and unanesthetized restrained rats. J. Neurosci. 26, 9038–9046.

Smythe, J.W., Colom, L. V., and Bland, B.H. (1992). The extrinsic modulation of hippocampal theta depends on the coactivation of cholinergic and GABA-ergic medial septal inputs. Neurosci. Biobehav. Rev. 16, 289–308.

Sotty, F., Danik, M., Manseau, F., Laplante, F., Quirion, R., and Williams, S. (2003). Distinct electrophysiological properties of glutamatergic, cholinergic and GABAergic rat septohippocampal neurons: Novel implications for hippocampal rhythmicity. J. Physiol. 551, 927–943.

Stewart, M., and Fox, S.E. (1989). Two populations of rhythmically bursting neurons in rat medial septum are revealed by atropine. J. Neurophysiol. 61.

Széll, A., Martínez-Bellver, S., Hegedüs, P., and Hangya, B. (2020). OPETH: Open Source Solution for Real-Time Peri-Event Time Histogram Based on Open Ephys. Front. Neuroinform. 14, 1–19.

Teitelbaum, H., Lee, J.F., and Johannessen, J.N. (1975). Behaviorally evoked hippocampal theta waves: A cholinergic response. Science (80-.). 188, 1114–1116.

Tokuda, K., Katori, Y., and Aihara, K. (2019). Chaotic dynamics as a mechanism of rapid transition of hippocampal local field activity between theta and non-theta states. Chaos 29, 113115.

Toth, K., Borhegyi, Z., and Freund, T.F. (1993). Postsynaptic targets of GABAergic hippocampal neurons in the medial septum-diagonal band of Broca complex. J. Neurosci. 13, 3712–3724.

Tsanov, M. (2015). Septo-hippocampal signal processing: Breaking the code. Prog. Brain Res. 219, 103–120.

Tsanov, M., Chah, E., Reilly, R., and O’Mara, S.M. (2014). Respiratory cycle entrainment of septal neurons mediates the fast coupling of sniffing rate and hippocampal theta rhythm. Eur. J. Neurosci. 39, 957–974.

Ujfalussy, B., and Kiss, T. (2006). How do glutamatergic and GABAergic cells contribute to synchronization in the medial septum? J. Comput. Neurosci. 21, 343–357.

Umbriaco, D., Watkins, K.C., Descarries, L., Cozzari, C., and Hartman, B.K. (1994). Ultrastructural and morphometric features of the acetylcholine innervation in adult rat parietal cortex: An electron microscopic study in serial sections. J. Comp. Neurol. 348, 351–373.

Unal, G., Joshi, A., Viney, T.J., Kis, V., and Somogyi, P. (2015). Synaptic Targets of Medial Septal Projections in the Hippocampus and Extrahippocampal Cortices of the Mouse. J. Neurosci. 35, 15812–15826.

Vandecasteele, M., Varga, V., Berényi, A., Papp, E., Barthó, P., Venance, L., Freund, T.F., and Buzsáki, G. (2014). Optogenetic activation of septal cholinergic neurons suppresses sharp wave ripples and enhances theta oscillations in the hippocampus. Proc. Natl. Acad. Sci. U. S. A. 111, 13535–13540.

Varga, V., Hangya, B., Kránitz, K., Ludányi, A., Zemankovics, R., Katona, I., Shigemoto, R., Freund, T.F., and Borhegyi, Z. (2008). The presence of pacemaker HCN channels identifies theta rhythmic GABAergic neurons in the medial septum. J. Physiol. 586, 3893–3915.

Wang, X.J. (2002). Pacemaker neurons for the theta rhythm and their synchronization in the septohippocampal reciprocal loop. J. Neurophysiol. 87, 889–900.

Wikenheiser, A.M., and Redish, D.A. (2013). The balance of forward and backward hippocampal sequences shifts across behavioral states. Hippocampus 23, 22–29.

Willms, A.R., Kitanov, P.M., and Langford, W.F. (2017). Huygens’ clocks revisited. R. Soc. Open Sci. 4, 170777.

Winterer, J., Lukacsovich, D., Que, L., Sartori, A.M., Luo, W., and Földy, C. (2019). Single‐cell RNA‐Seq characterization of anatomically identified OLM interneurons in different transgenic mouse lines. Eur. J. Neurosci. 50, 3750–3771.

Wolansky, T., Clement, E.A., Peters, S.R., Palczak, M.A., and Dickson, C.T. (2006). Hippocampal Slow Oscillation: A Novel EEG State and Its Coordination with Ongoing Neocortical Activity. J. Neurosci. 26, 6213–6229.

Xu, C., Datta, S., Wu, M., and Alreja, M. (2004). Hippocampal theta rhythm is reduced by suppression of the H-current in septohippocampal GABAergic neurons. Eur. J. Neurosci. 19, 2299–2309.

Yang, C., McKenna, J.T., Zant, J.C., Winston, S., Basheer, R., and Brown, R.E. (2014). Cholinergic neurons excite cortically projecting basal forebrain GABAergic neurons. J. Neurosci. 34, 2832–2844.

Yang, C., Thankachan, S., McCarley, R.W., and Brown, R.E. (2017). The menagerie of the basal forebrain: how many (neural) species are there, what do they look like, how do they behave and who talks to whom? Curr. Opin. Neurobiol. 44, 159–166.

Yoder, R.M., and Pang, K.C.H. (2005). Involvement of GABAergic and cholinergic medial septal neurons in hippocampal theta rhythm. Hippocampus 15, 381–392.

Zaborszky, L., van den Pol, A., and Gyengesi, E. (2012). The Basal Forebrain Cholinergic Projection System in Mice. In The Mouse Nervous System, C. Watson, G. Paxinos, and L. Puelles, eds. (Amsterdam: Elsevier), pp. 684–718.

Zemankovics, R., Káli, S., Paulsen, O., Freund, T.F., and Hájos, N. (2010). Differences in subthreshold resonance of hippocampal pyramidal cells and interneurons: the role of h‐current and passive membrane characteristics. J. Physiol. 588, 2109–2132.

Zou, X., Coyle, D., Wong-Lin, K.F., and Maguire, L. (2011). Computational study of Hippocampal-septal theta rhythm changes due to Beta-Amyloid-Altered ionic channels. PLoS One 6.

Zutshi, I., Brandon, M.P., Fu, M.L., Donegan, M.L., Leutgeb, J.K., and Leutgeb, S. (2018). Hippocampal Neural Circuits Respond to Optogenetic Pacing of Theta Frequencies by Generating Accelerated Oscillation Frequencies. Curr. Biol. 28, 1179–1188.e3.

