## Supplemental material for "Huygens synchronization of medial septal pacemaker neurons generates hippocampal theta oscillation"

**Supplemental Figures**

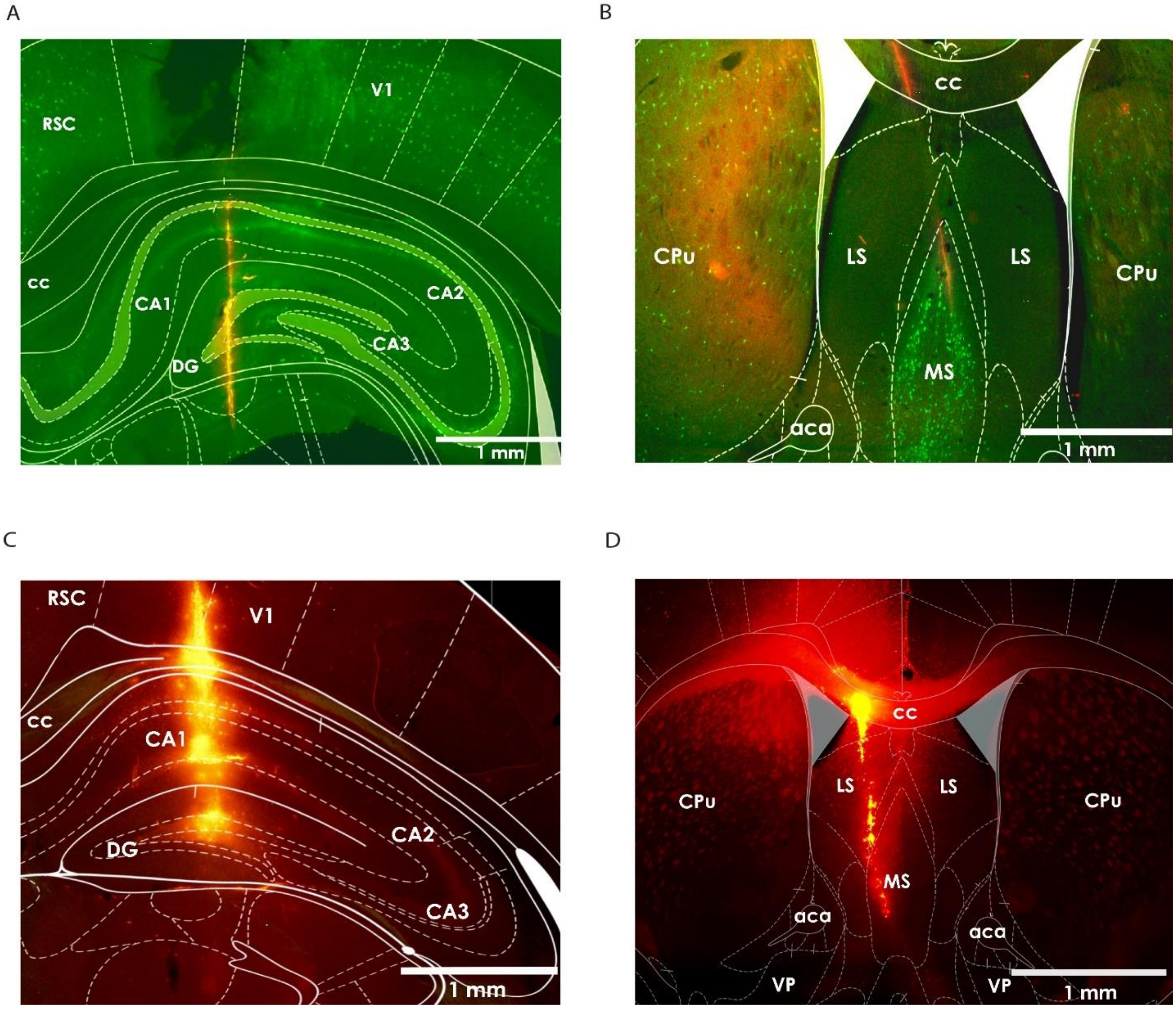


**Figure S1. Histological track reconstruction in mice and rats.** (A) Fluorescent images of a silicon probe track in the hippocampus from an acute rat recording. Green, ChAT staining; red, DiI. (B) Silicon probe track in the medial septum from an acute rat recording. (C) Silicon probe track in the hippocampus from a chronic mouse recording. Red, DiI. (D) Silicon probe track in the medial septum from a chronic mouse recording. See also Figure 1A for track reconstruction after acute mouse experiment. aca, anterior commissure; CA, cornu ammonis; cc, corpus callosum; CPu, caudate putamen; DG, dentate gyrus; LS, lateral septum; MS, medial septum; RSC, retrosplenial cortex; V1, primary visual cortex; VP, ventral pallidum.

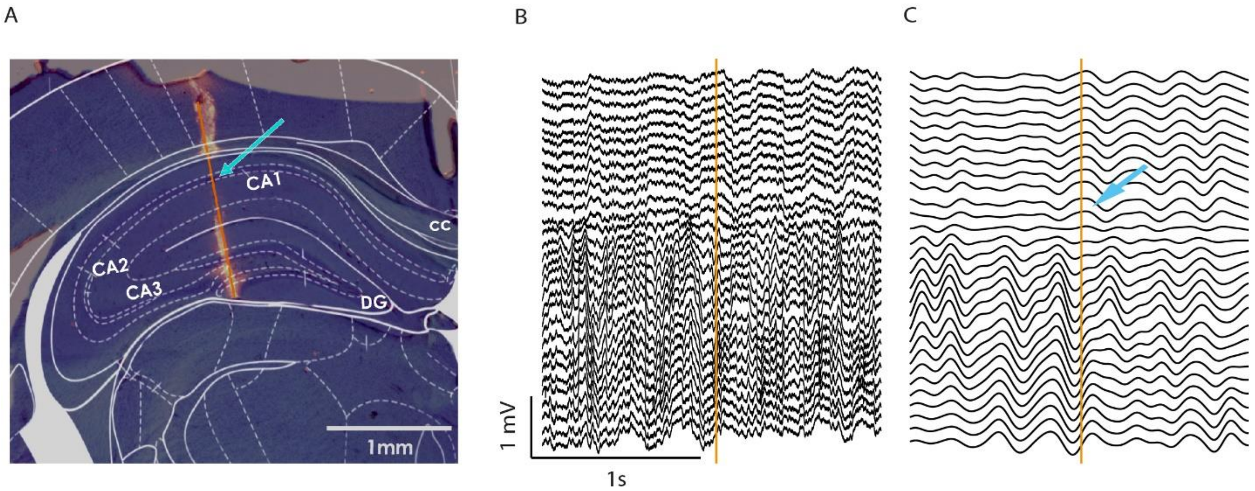


**Figure S2. Phase reversal of hippocampal theta oscillation.** (A) Fluorescent image of the silicon probe track in the hippocampus from an acute mouse experiment. An estimated 10% of tissue shrinkage during perfusion and histological processing was taken into account. (B) Raw 32-channels LFP recording from the same experiment (two broken channels were removed). (C) LFP channels filtered in the theta band. Theta phase reversal, known to occur just below the pyramidal layer, is indicated by cyan arrow. The reconstructed position of the corresponding electrode contact site is marked by cyan arrow in panel (A).

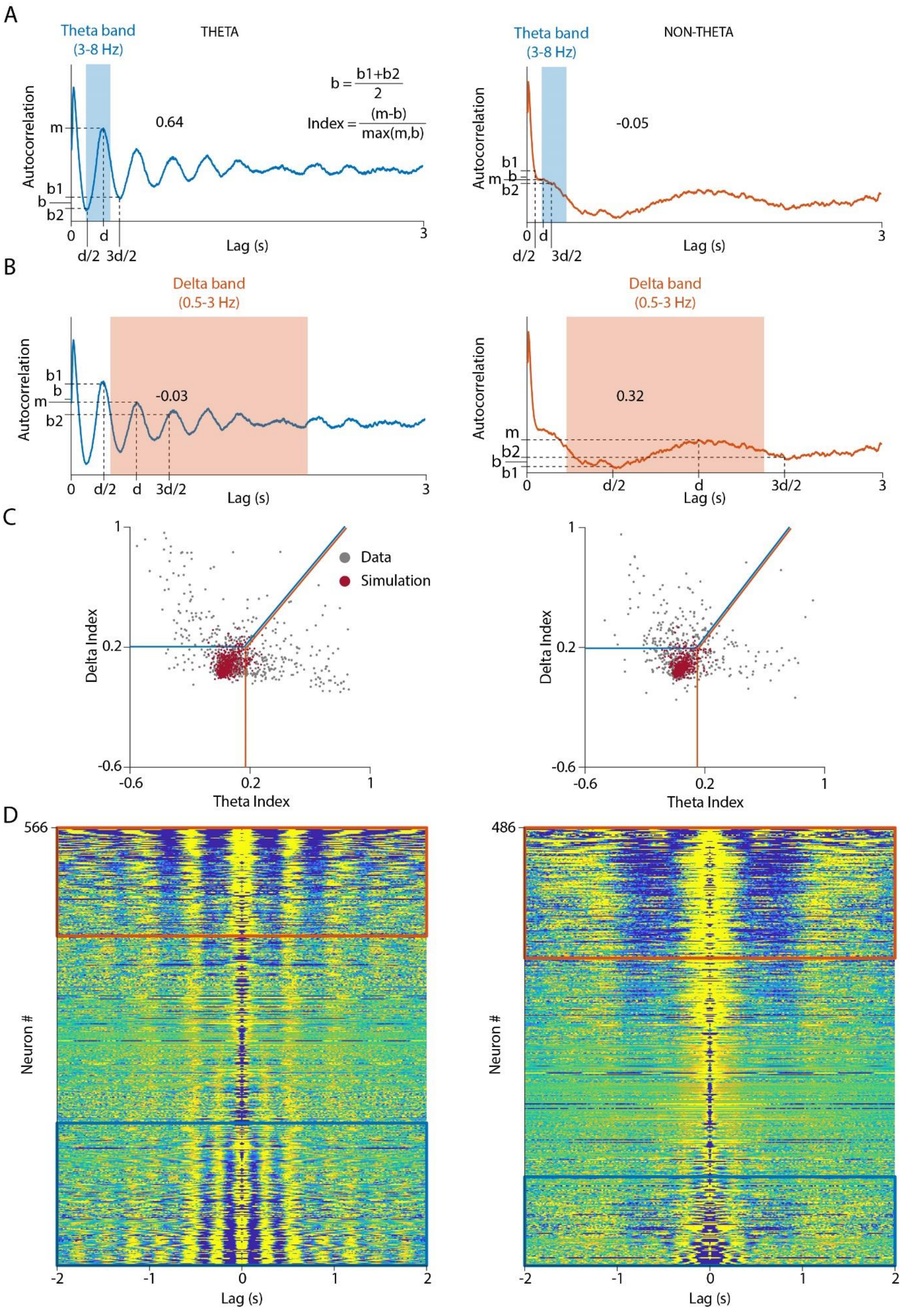

A
Theta band (3-8 Hz)
THETA
b = (b1+b2)/2
Index = (m-b)/max(m,b)
0.64
Autocorrelation
m
b1
b
b2
0
d
d/2
3d/2
3
Lag (s)
Theta band (3-8 Hz)
NON-THETA
-0.05
B
Delta band (0.5-3 Hz)
-0.03
0.32
C
Data
Simulation
Delta Index
Theta Index
D
566
486
Neuron #
Lag (s)

**Figure S3. Determining theta and delta rhythmicity.** (A) Theta Index was calculated based on autocorrelograms for theta and non-theta segments by normalizing the theta-band autocorrelation peak with the average of pre- and post-peak troughs (see Methods). The number above the trace denotes the calculated value for the rhythmicity index in the example. (B) Delta Index was calculated using a similar algorithm for the delta band. (C) Theta and Delta Index was calculated for all recorded neurons during theta (left) and non-theta segments (right; grey, recordings from urethane-anesthetized rats are shown) and for simulated Poisson-neurons with matching firing rate (red). Significant rhythmicity was defined by the 0.05 percentile of the Theta and Delta Index distributions of the Poisson-neurons (vertical and horizontal lines). If both indices exceeded the threshold, the larger of the two rhythmicity indices was considered (identity line). (D) Autocorrelograms (left, theta segments; right, non-theta segments) for all neurons with sufficient firing rate (see Methods), sorted by the larger of the two rhythmicity indices: delta-rhythmic on top and theta-rhythmic on bottom; significantly delta- and theta-modulated neurons are above and below the respective colored lines.

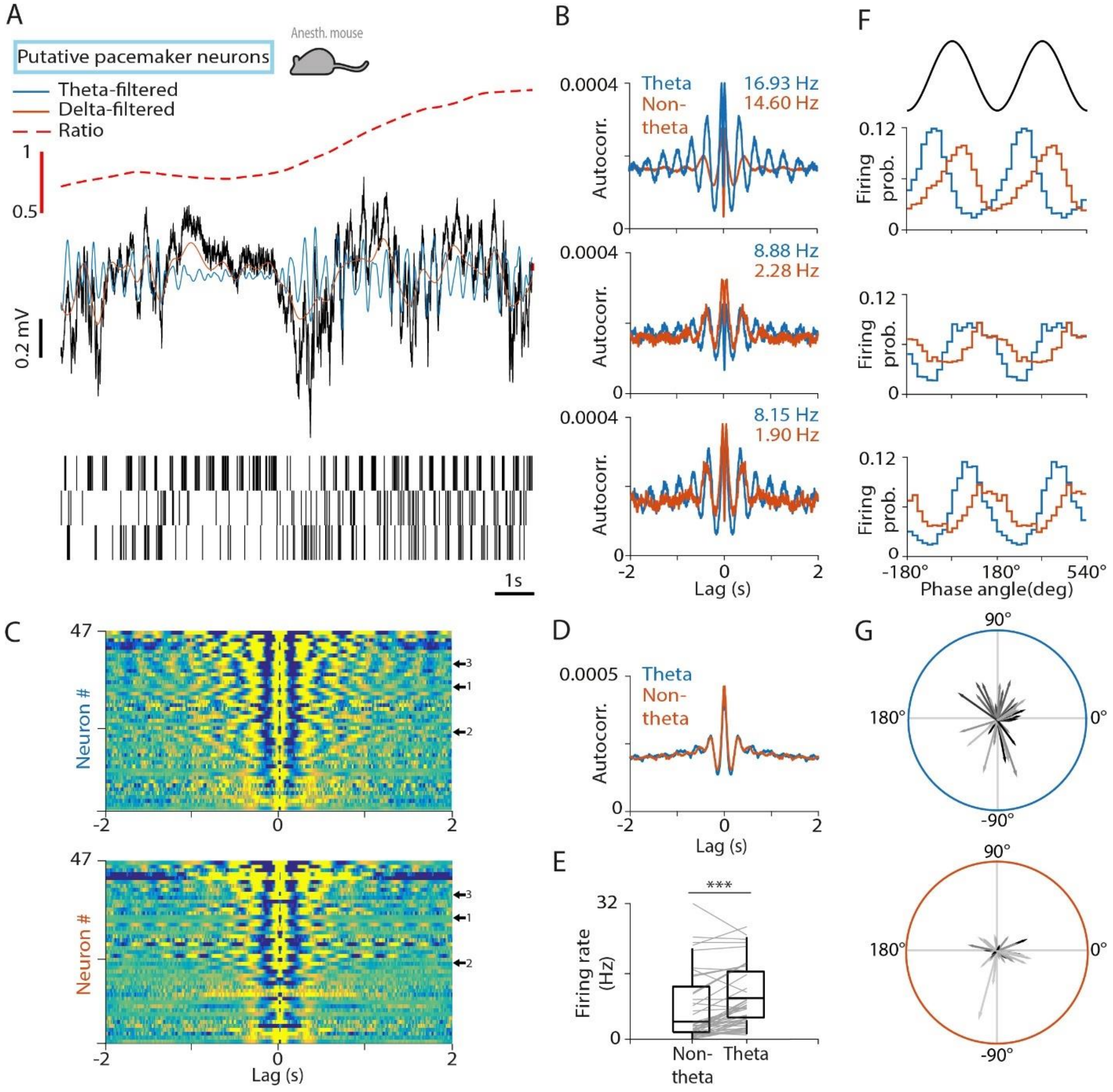


**Figure S4. Putative pacemaker neurons of the MS in urethane-anesthetized mice.** (A) Top, black, raw LFP from the CA1 shows a delta-to-theta state transition. Orange, LFP filtered in the delta band; blue, LFP filtered in the theta band; dashed, theta-delta amplitude ratio. Bottom, spike raster of three examples of putative pacemaker neurons from the same recording session. Recordings are from urethane-anesthetized mice. (B) Autocorrelograms of the three example neurons in panel (A). Numbers indicate average firing rates during theta and non-theta segments. (C) Autocorrelograms of all putative pacemaker neurons during theta (top) and non-theta (bottom) segments. Arrows indicate the example neurons. (D) Average autocorrelogram of all putative pacemaker neurons. (E) Median firing rate of putative pacemaker neurons during non-theta and theta segments. Boxes and whiskers represent interquartile ranges and non-outlier ranges, respectively. Lines correspond to individual neurons. Firing rate was significantly higher during theta. ***, $p < 0.001$, Wilcoxon signed-rank test. (F) Phase histogram of the example neurons in panel (A) relative to delta (orange) and theta (blue) oscillations. Two oscillatory cycles are shown. (G) Phase-locking of all putative pacemaker neurons to theta (top) and delta (bottom) oscillation in polar coordinates. Angle, circular phase; length, mean vector length; greyscale corresponds to average firing rate (black, higher firing rate). Phase-locking strength to delta oscillation was low for most putative pacemaker neurons.

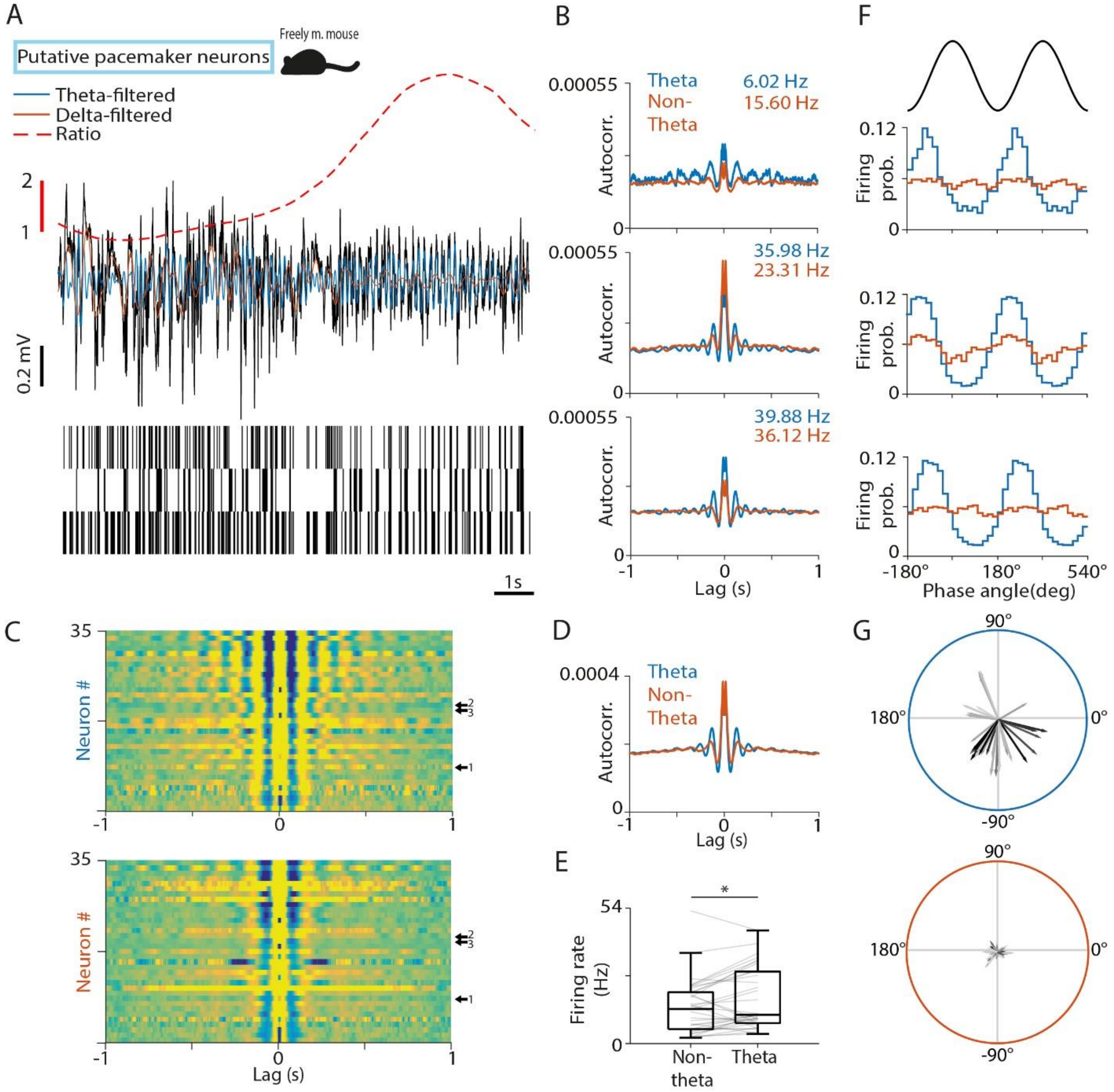


**Figure S5. Putative pacemaker neurons of the MS in freely moving mice.** (A) Top, black, raw LFP from the CA1 shows a delta-to-theta state transition. Orange, LFP filtered in the delta band; blue, LFP filtered in the theta band; dashed, theta-delta amplitude ratio. Bottom, spike raster of three examples of putative pacemaker neurons from the same recording session. Recordings are from drug-free mice. (B) Autocorrelograms of the three example neurons in panel (A). Numbers indicate average firing rates during theta and non-theta segments. (C) Autocorrelograms of all putative pacemaker neurons during theta (top) and non-theta (bottom) segments. Arrows indicate the example neurons. (D) Average autocorrelogram of all putative pacemaker neurons. (E) Median firing rate of putative pacemaker neurons during non-theta and theta segments. Boxes and whiskers represent interquartile ranges and non-outlier ranges, respectively. Lines correspond to individual neurons. Firing rate was significantly higher during theta. *, $p < 0.05$, Wilcoxon signed-rank test. (F) Phase histogram of the example neurons in panel (A) relative to delta (orange) and theta (blue) oscillations. Two oscillatory cycles are shown. (G) Phase-locking of all putative pacemaker neurons to theta (top) and delta (bottom) oscillation in polar coordinates. Angle, circular phase; length, mean vector length; greyscale corresponds to average firing rate (black, higher firing rate). Phase-locking strength to delta oscillation was low for putative pacemaker neurons.

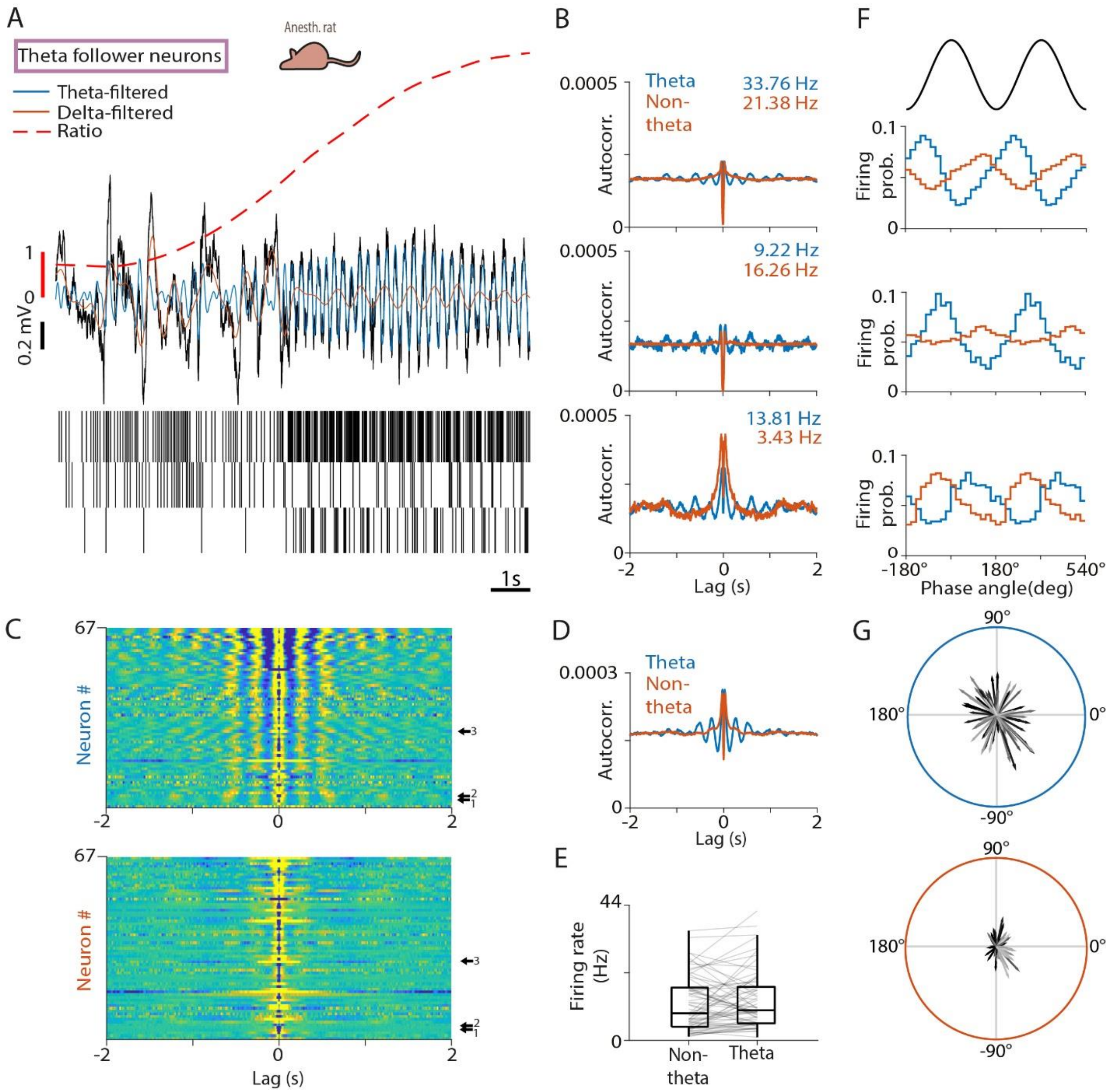


**Figure S6. Theta follower neurons in the MS of urethane-anesthetized rats.** (A) Top, black, raw LFP from the CA1 shows a delta-to-theta state transition. Orange, LFP filtered in the delta band; blue, LFP filtered in the theta band; dashed, theta-delta amplitude ratio. Bottom, spike raster of three example neurons from the same recording session. ‘Theta follower’ neurons showed non-rhythmic activity during non-theta epochs and theta-rhythmic activity during theta epochs in urethane-anesthetized rats. See Figure S8 and Figure S9 for mouse recordings. (B) Autocorrelograms of the three example neurons in panel (A). Numbers indicate average firing rates during theta and non-theta segments. (C) Autocorrelograms of all theta follower neurons during theta (top) and non-theta (bottom) segments. Arrows indicate the example neurons. (D) Average autocorrelogram of all neurons. (E) Median firing rate of neurons during non-theta and theta segments. Boxes and whiskers represent interquartile ranges and non-outlier ranges, respectively. Lines correspond to individual neurons. No systematic firing rate change was found. (F) Phase histogram of the example neurons in panel (A) relative to delta (orange) and theta (blue) oscillations. Two oscillatory cycles are shown. (G) Phase-locking of all theta follower neurons to theta (top) and delta (bottom) oscillation in polar coordinates. Angle, circular phase; length, mean vector length; greyscale corresponds to average firing rate (black, higher firing rate). Many of these neurons were phase-locked to both hippocampal oscillations.

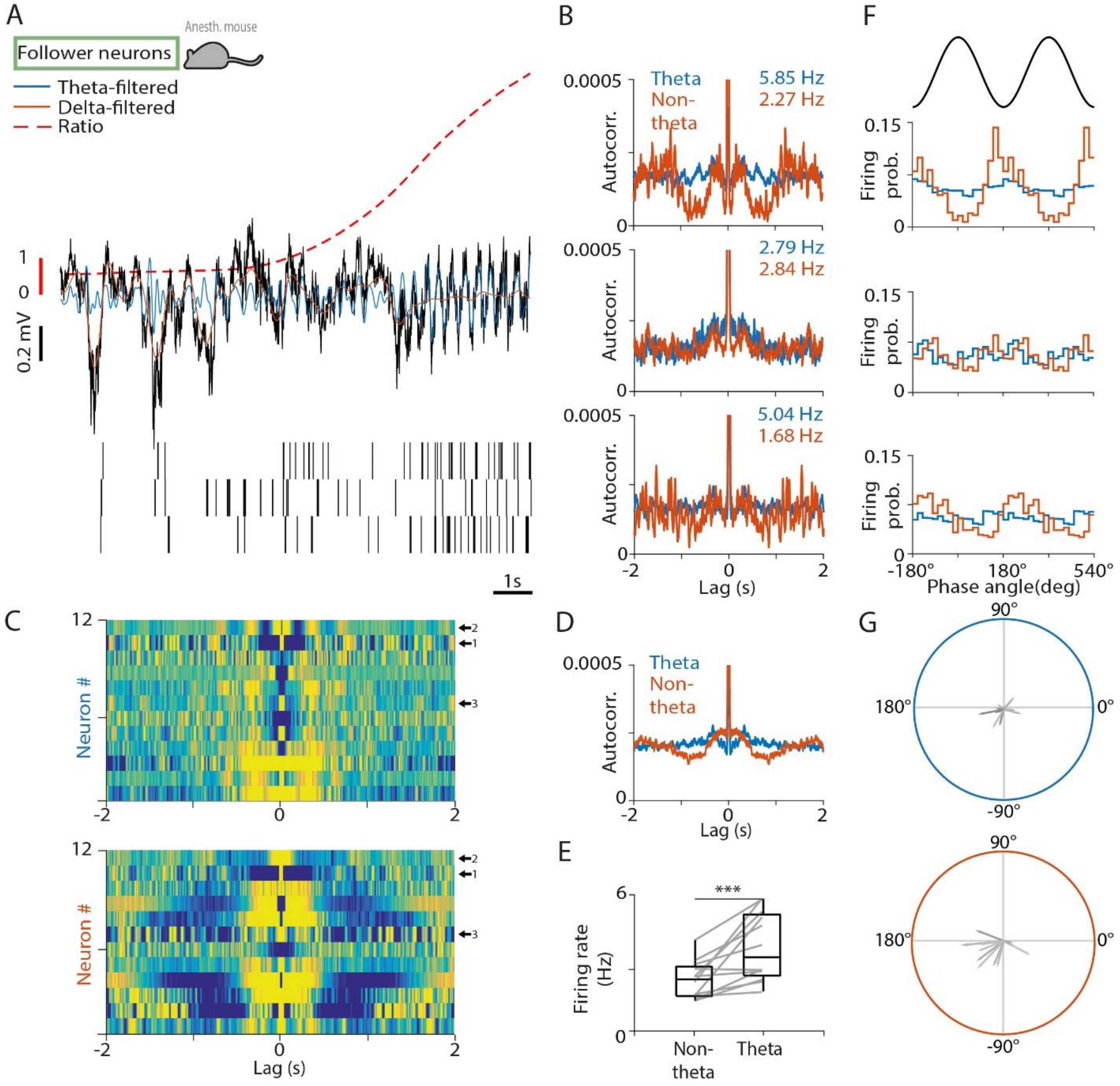


**Figure S7. Follower MS neurons in urethane-anesthetized mice.** (A) Top, black, raw LFP from the CA1 shows a delta-to-theta state transition. Orange, LFP filtered in the delta band; blue, LFP filtered in the theta band; dashed, theta-delta amplitude ratio. Bottom, spike raster of three example neurons from the same recording session. 'Follower' neurons showed delta-rhythmic activity during non-theta epochs and theta-rhythmic activity during theta epochs. (B) Autocorrelograms of the three example neurons in panel (A). Numbers indicate average firing rates during theta and non-theta segments. (C) Autocorrelograms of all follower neurons during theta (top) and non-theta (bottom) segments. Arrows indicate the example neurons. (D) Average autocorrelogram of all follower neurons. (E) Median firing rate of follower neurons during non-theta and theta segments. Boxes and whiskers represent interquartile ranges and non-outlier ranges, respectively. Lines correspond to individual neurons. Firing rate was significantly higher during theta. ***, $p < 0.001$, Wilcoxon signed-rank test. (F) Phase histogram of the example neurons in panel (A) relative to delta (orange) and theta (blue) oscillations. Two oscillatory cycles are shown. (G) Phase-locking of all follower neurons to theta (top) and delta (bottom) oscillation in polar coordinates. Angle, circular phase; length, mean vector length; greyscale corresponds to average firing rate (black, higher firing rate).

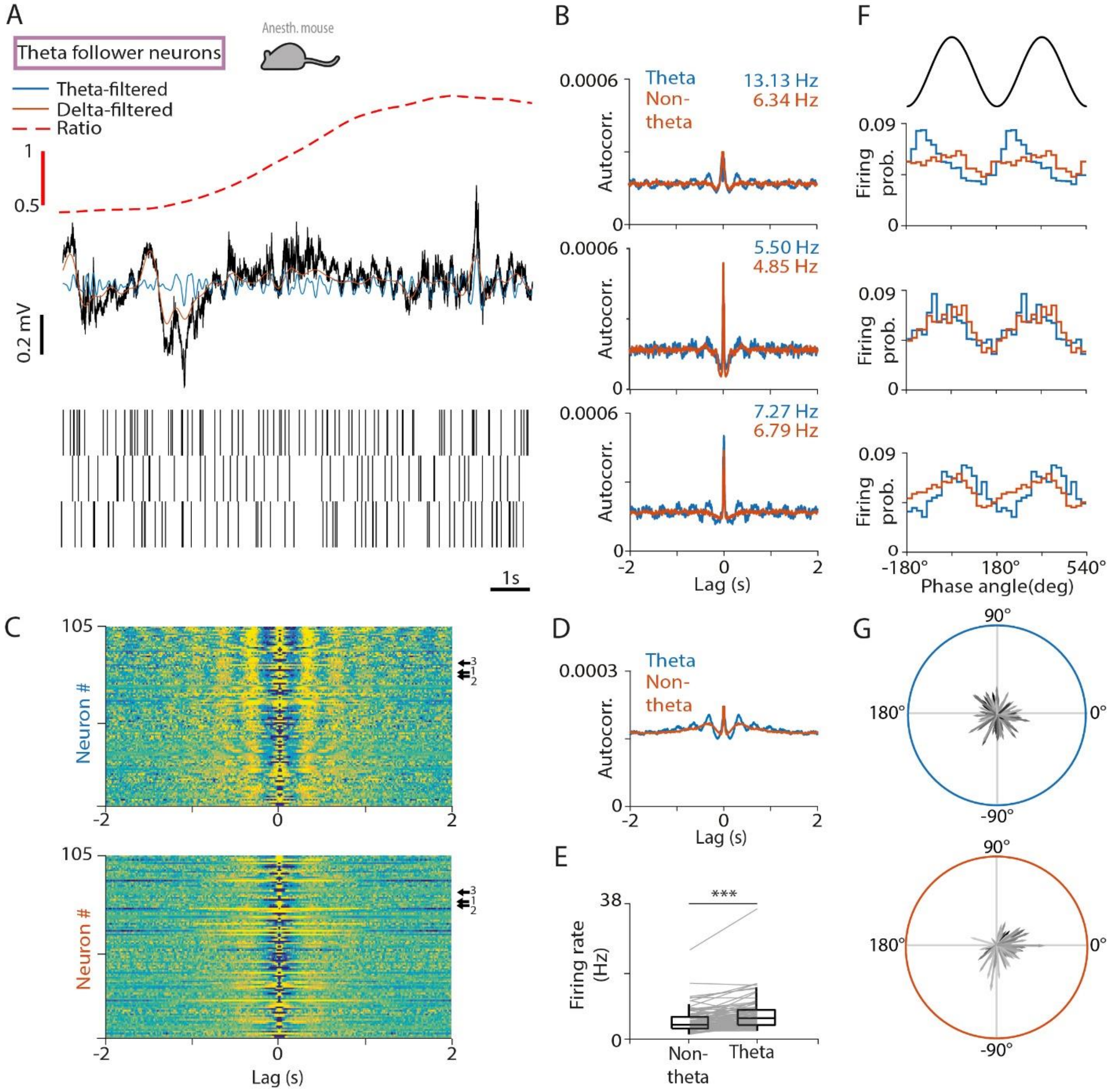


**Figure S8. Theta follower neurons in the MS of urethane-anesthetized mice.** (A) Top, black, raw LFP from the CA1 shows a delta-to-theta state transition. Orange, LFP filtered in the delta band; blue, LFP filtered in the theta band; dashed, theta-delta amplitude ratio. Bottom, spike raster of three example neurons from the same recording session. 'Theta follower' neurons showed non-rhythmic activity during non-theta epochs and theta-rhythmic activity during theta epochs. (B) Autocorrelograms of the three example neurons in panel (A). Numbers indicate average firing rates during theta and non-theta segments. (C) Autocorrelograms of all theta follower neurons during theta (top) and non-theta (bottom) segments. Arrows indicate the example neurons. (D) Average autocorrelogram of all neurons. (E) Median firing rate of neurons during delta and theta oscillation. Boxes and whiskers represent interquartile ranges and non-outlier ranges, respectively. Lines correspond to individual neurons. Firing rate was significantly higher during theta. ***, $p < 0.001$, Wilcoxon signed-rank test. (F) Phase histogram of the example neurons in panel (A) relative to delta (orange) and theta (blue) oscillations. Two oscillatory cycles are shown. (G) Phase-locking of all theta follower neurons to theta (top) and delta (bottom) oscillation in polar coordinates. Angle, circular phase; length, mean vector length; greyscale corresponds to average firing rate (black, higher firing rate).

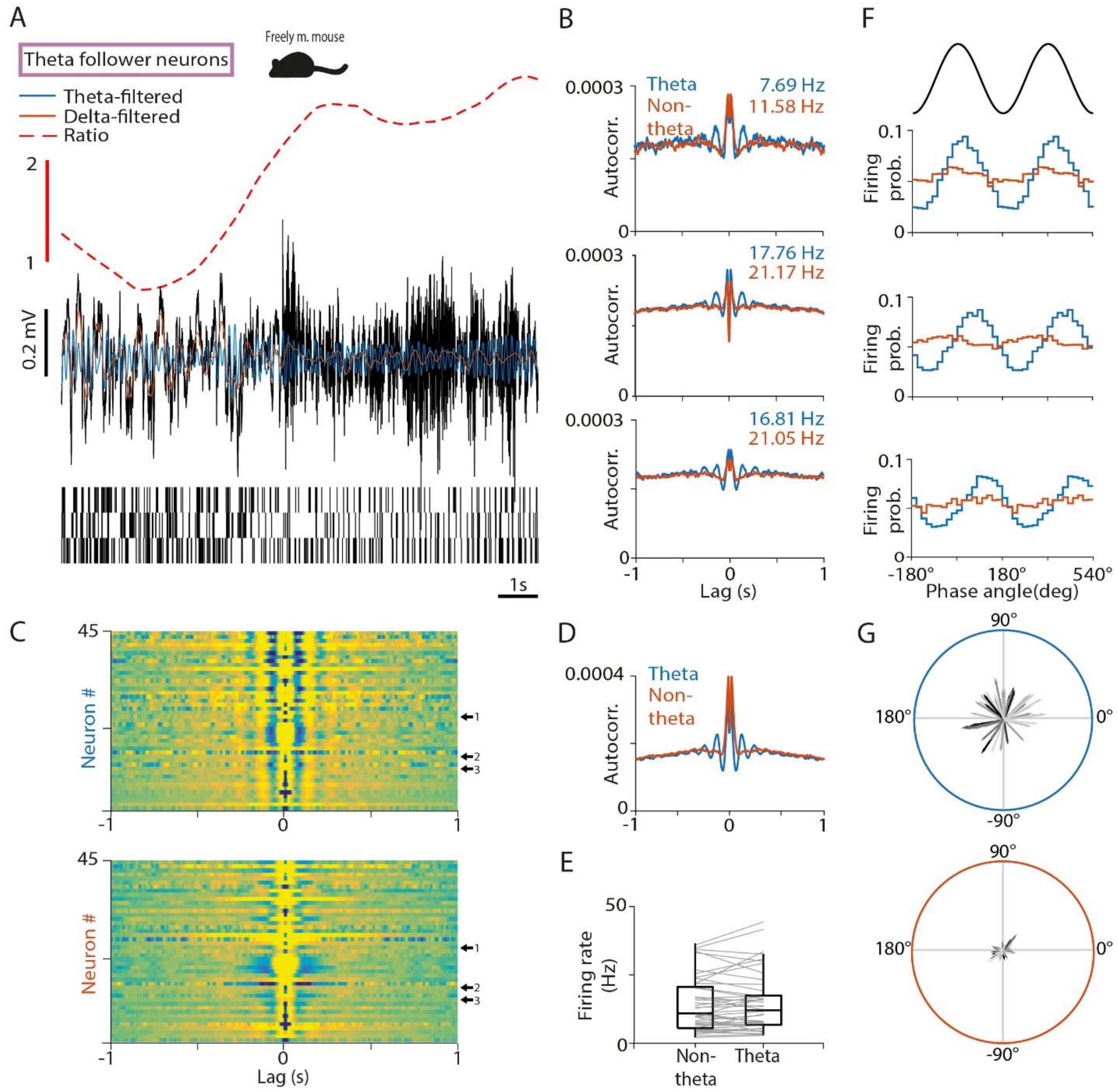


**Figure S9. Theta follower neurons in the MS of freely moving mice.** (A) Top, black, raw LFP from the CA1 shows a delta-to-theta state transition. Orange, LFP filtered in the delta band; blue, LFP filtered in the theta band; dashed, theta-delta amplitude ratio. Bottom, spike raster of three example neurons from the same recording session. Theta follower neurons showed non-rhythmic activity during non-theta epochs and theta-rhythmic activity during theta epochs. (B) Autocorrelograms of the three example neurons in panel (A). Numbers indicate average firing rates during theta and non-theta segments. (C) Autocorrelograms of all theta follower neurons during theta (top) and non-theta (bottom) segments. Arrows indicate the example neurons. (D) Average autocorrelogram of all neurons. (E) Median firing rate of neurons during non-theta and theta segments. Boxes and whiskers represent interquartile ranges and non-outlier ranges, respectively. Lines correspond to individual neurons. No systematic firing rate change was found. (F) Phase histogram of the example neurons in panel (A) relative to delta (orange) and theta (blue) oscillations. Two oscillatory cycles are shown. (G) Phase-locking of all theta follower neurons to theta (top) and delta (bottom) oscillation in polar coordinates. Angle, circular phase; length, mean vector length; greyscale corresponds to average firing rate (black, higher firing rate).

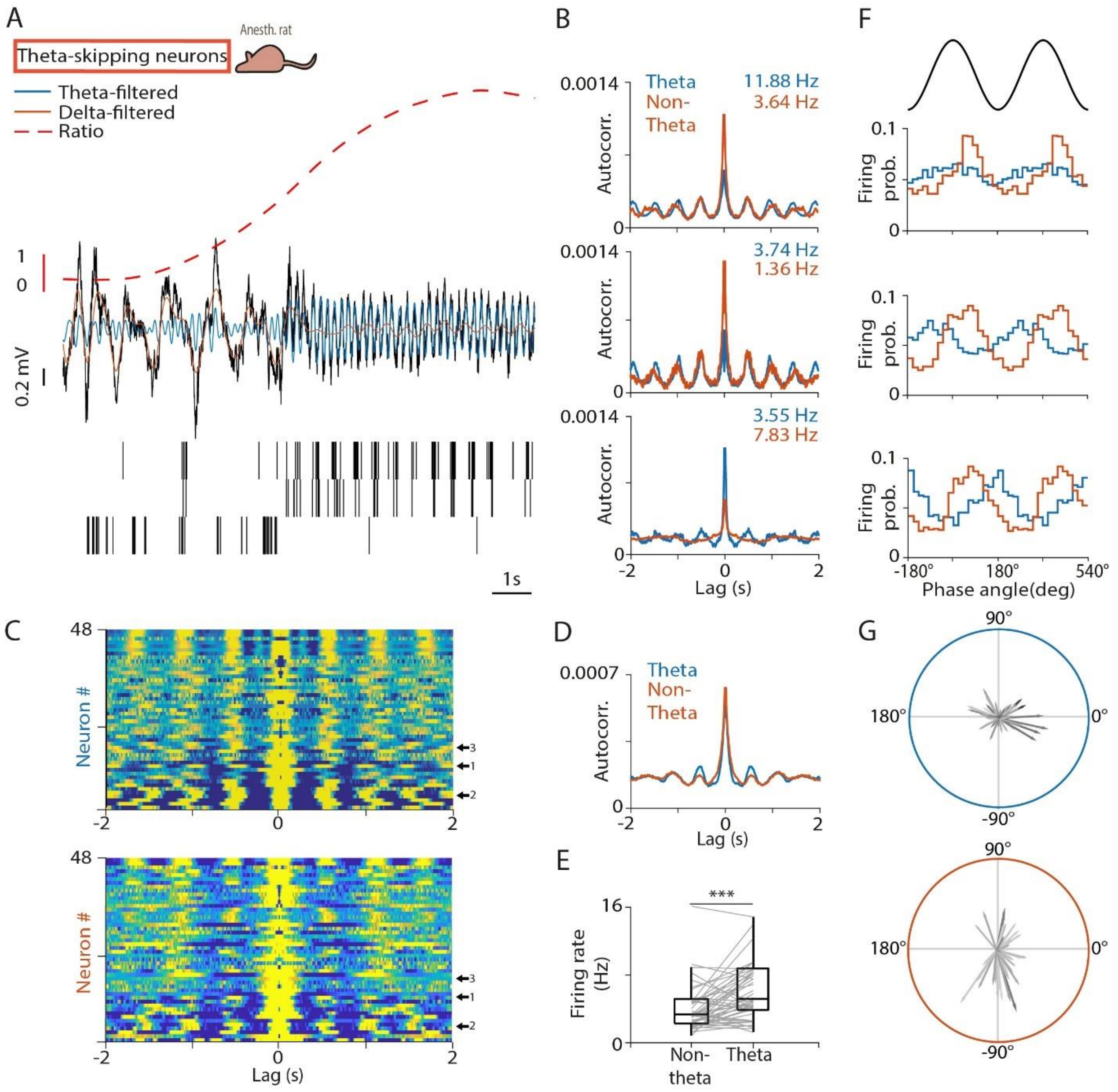


**Figure S10. Theta-skipping neurons of the MS.** (A) Top, black, raw LFP from the CA1 shows a delta-to-theta state transition. Orange, LFP filtered in the delta band; blue, LFP filtered in the theta band; dashed, theta-delta amplitude ratio. Bottom, spike raster of three examples of theta-skipping neurons from the same recording session. Recordings are from urethane-anesthetized rats; see Figure S11 for the anesthetized mouse recordings. (B) Autocorrelograms of the three example neurons in panel (A). Numbers indicate average firing rates during theta and non-theta segments. (C) Autocorrelograms of all theta-skipping neurons during theta (top) and non-theta (bottom) segments. Arrows indicate the example neurons. (D) Average autocorrelogram of all theta-skipping neurons. (E) Median firing rate of theta-skipping neurons during non-theta and theta segments. Boxes and whiskers represent interquartile ranges and non-outlier ranges, respectively. Lines correspond to individual neurons. Firing rate was significantly higher during theta. ***, $p < 0.001$, Wilcoxon signed-rank test. (F) Phase histogram of the example neurons in panel (A) relative to delta (orange) and theta (blue) oscillations. Two oscillatory cycles are shown. (G) Phase-locking of all theta-skipping neurons to theta (top) and delta (bottom) oscillation in polar coordinates. Angle, circular phase; length, mean vector length; greyscale corresponds to average firing rate (black, higher firing rate).

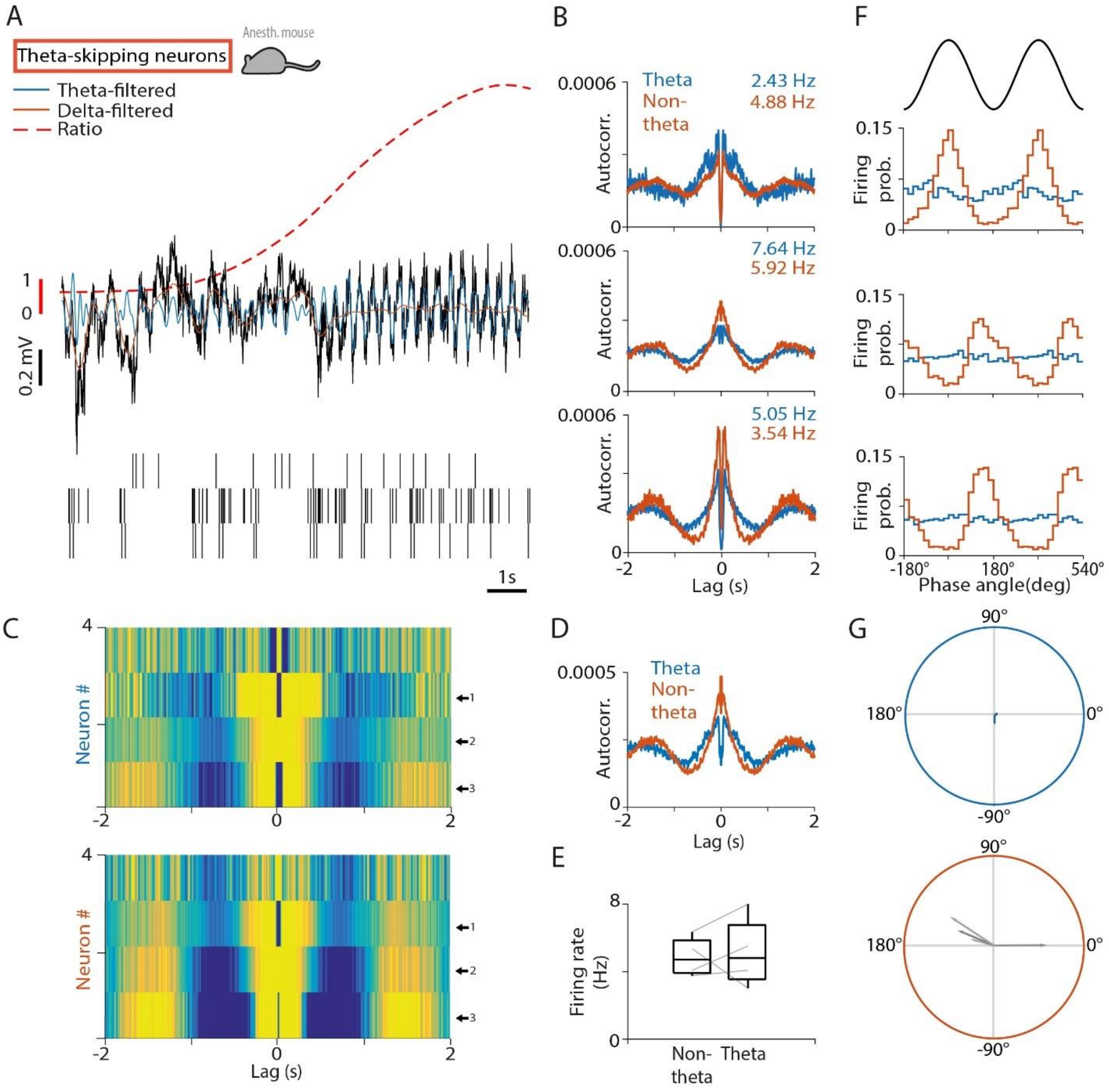


**Figure S11. Theta-skipping neurons of the MS in urethane-anesthetized mice.** (A) Top, black, raw LFP from the CA1 shows a delta-to-theta state transition. Orange, LFP filtered in the delta band; blue, LFP filtered in the theta band; dashed, theta-delta amplitude ratio. Bottom, spike raster of three examples of theta-skipping neurons from the same recording session. Recordings are from urethane-anesthetized mice. (B) Autocorrelograms of the three example neurons in panel (A). Numbers indicate average firing rates during theta and non-theta segments. (C) Autocorrelograms of all theta-skipping neurons during theta (top) and non-theta (bottom) segments. Arrows indicate the example neurons. (D) Average autocorrelogram of all theta-skipping neurons. (E) Median firing rate of theta-skipping neurons during non-theta and theta segments. Boxes and whiskers represent interquartile ranges and non-outlier ranges, respectively. Lines correspond to individual neurons. (F) Phase histogram of the example neurons in panel (A) relative to delta (orange) and theta (blue) oscillations. Two oscillatory cycles are shown. (G) Phase-locking of all theta-skipping neurons to theta (top) and delta (bottom) oscillation in polar coordinates. Angle, circular phase; length, mean vector length; greyscale corresponds to average firing rate (black, higher firing rate).

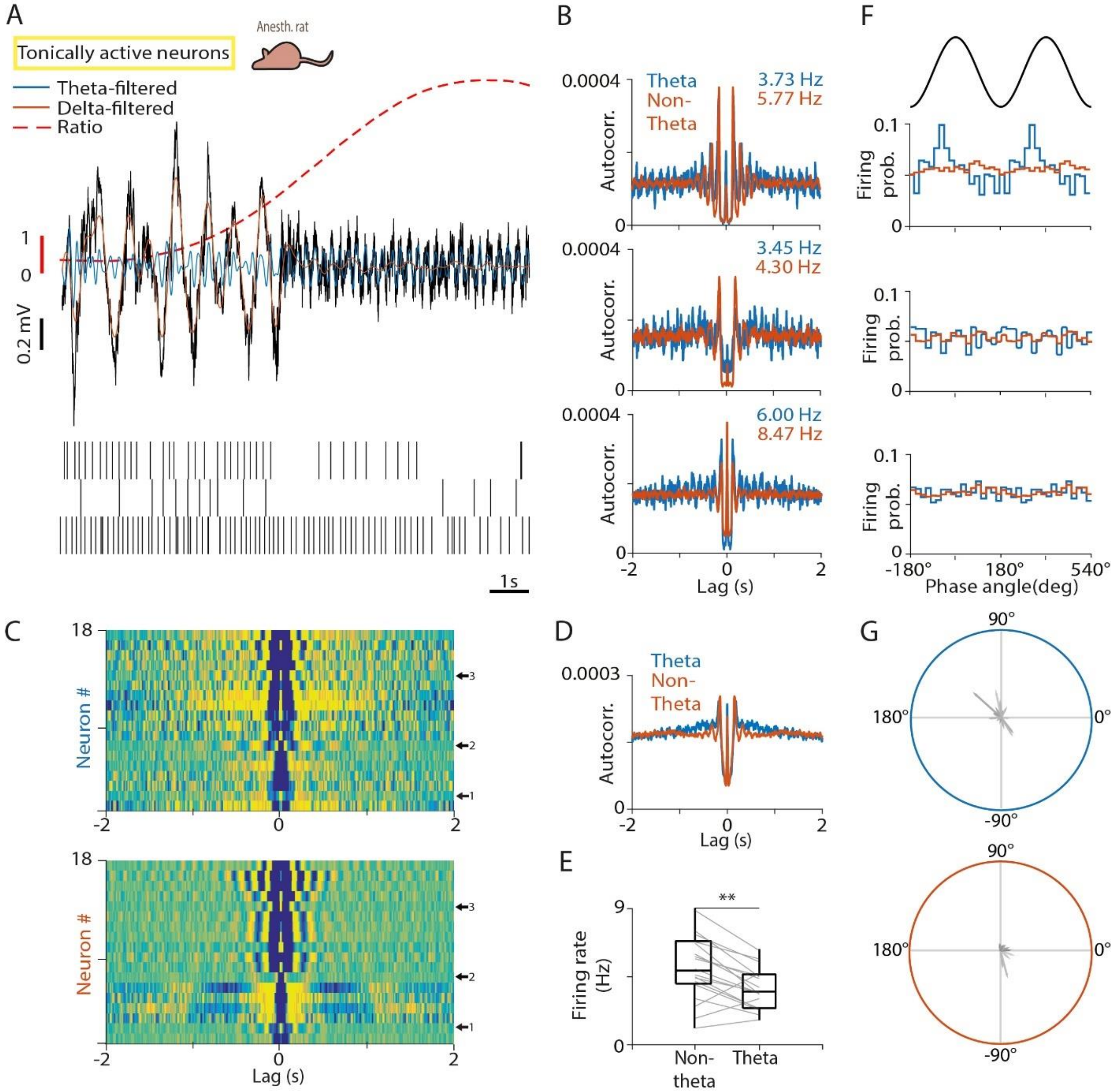


**Figure S12. Tonically active MS neurons.** (A) Top, black, raw LFP from the CA1 shows a delta-to-theta state transition. Orange, LFP filtered in the delta band; blue, LFP filtered in the theta band; dashed, theta-delta amplitude ratio. Bottom, spike raster of three examples of tonically active neurons from the same recording session. Recordings are from urethane-anesthetized rats; see Figure S14 and Figure S15 for the anesthetized and awake mouse recordings. (B) Autocorrelograms of the three example neurons in panel (A). Numbers indicate average firing rates during theta and non-theta segments. (C) Autocorrelograms of all tonically active neurons during theta (top) and non-theta (bottom) segments. Arrows indicate the example neurons. (D) Average autocorrelogram of all tonically active neurons. (E) Median firing rate of tonically active neurons during non-theta and theta epochs. Boxes and whiskers represent interquartile ranges and non-outlier ranges, respectively. Lines correspond to individual neurons. Firing rate was significantly higher during non-theta. **, $p < 0.01$, Wilcoxon signed-rank test. (F) Phase histogram of the example neurons in panel (A) relative to delta (orange) and theta (blue) oscillations. Two oscillatory cycles are shown. (G) Phase-locking of all tonically active neurons to theta (top) and delta (bottom) oscillation in polar coordinates. Angle, circular phase; length, mean vector length; greyscale corresponds to average firing rate (black, higher firing rate). These neurons did not show strong phase-locking to hippocampal oscillations.

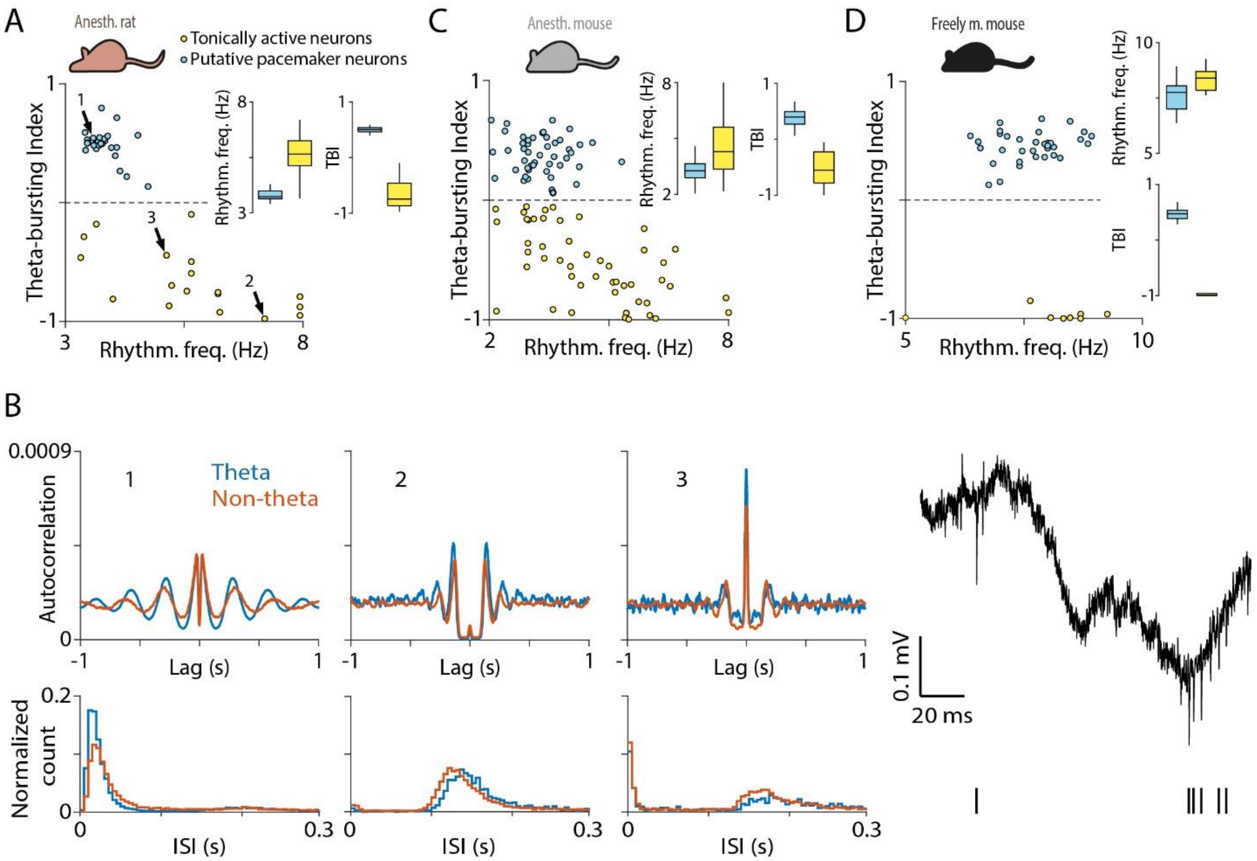


**Figure S13. Separating theta-bursting and tonically active neurons.** (A) Theta-bursting Index (TBI; see Methods) and rhythmicity frequency revealed a population that showed theta-bursting and a separate one that did not, in anesthetized rats. The former corresponded to putative pacemakers (Figure 2), while the latter formed the group of 'tonically active' neurons (Figure S12). (B) Left, autocorrelogram (top) and interspike interval (ISI) histogram of three example neurons, marked by arrows in panel (A). Example 1 is a theta-bursting pacemaker; examples 2 and 3 are tonically active neurons. Right, raw data (top) and spike raster (bottom) of example 3. Note the fast-bursting phenotype, different from the slower theta-bursting observed in pacemakers. (C-D) The same distinction was observed in anesthetized (C) and freely moving mice (D). Box-whisker plots show median, interquartile range and non-outlier range.

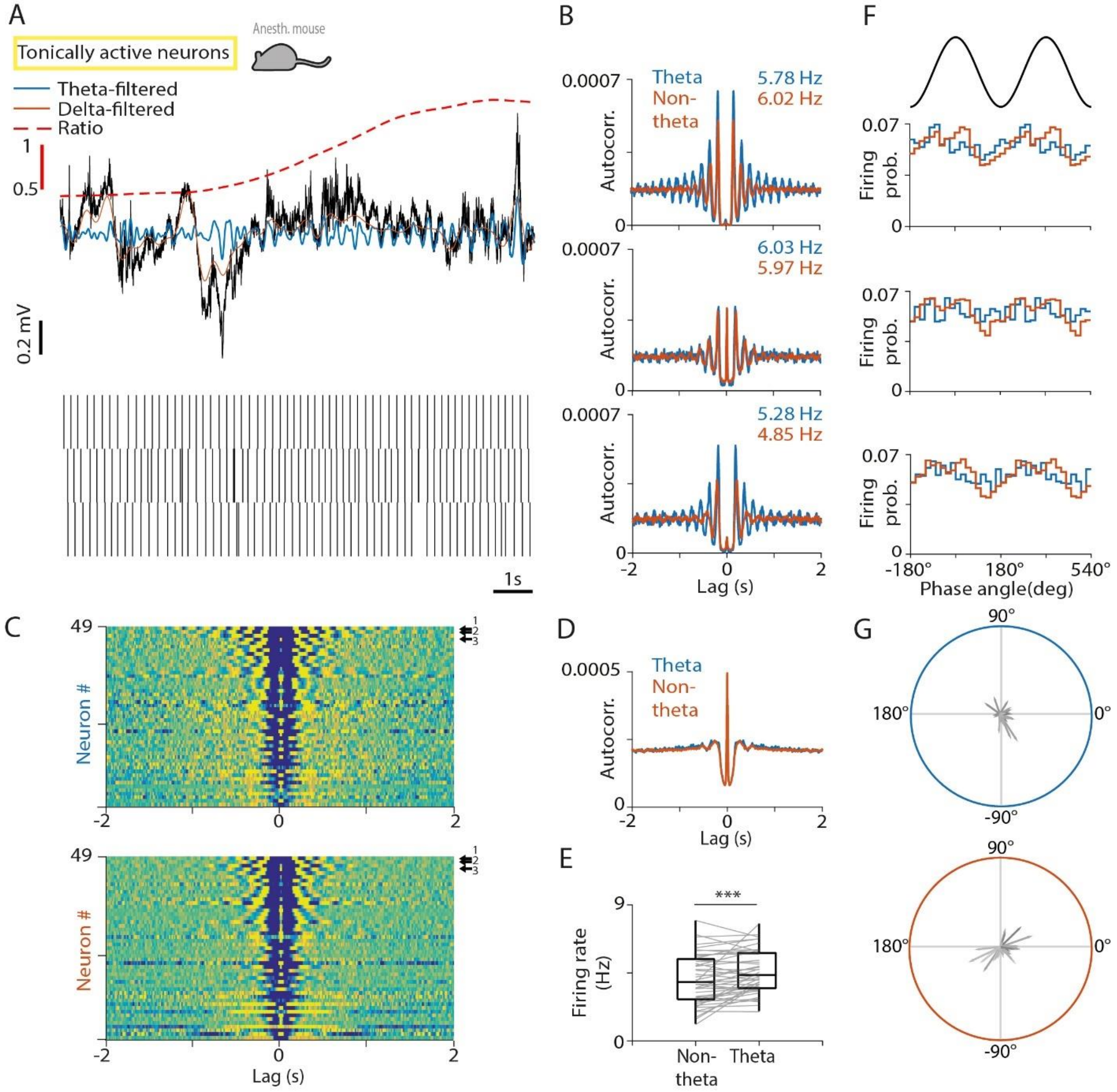


**Figure S14. Tonically active MS neurons in urethane-anesthetized mice.** (A) Top, black, raw LFP from the CA1 shows a delta-to-theta state transition. Orange, LFP filtered in the delta band; blue, LFP filtered in the theta band; dashed, theta-delta amplitude ratio. Bottom, spike raster of three examples of tonically active neurons from the same recording session. Recordings are from urethane-anesthetized mice. (B) Autocorrelograms of the three example neurons in panel (A). Numbers indicate average firing rates during theta and non-theta segments. (C) Autocorrelograms of all tonically active neurons during theta (top) and non-theta (bottom) segments. Arrows indicate the example neurons. (D) Average autocorrelogram of all tonically active neurons. (E) Median firing rate of tonically active neurons during non-theta and theta segments. Boxes and whiskers represent interquartile ranges and non-outlier ranges, respectively. Lines correspond to individual neurons. Firing rate was significantly higher during theta. ***, $p < 0.001$, Wilcoxon signed-rank test. (F) Phase histogram of the example neurons in panel (A) relative to delta (orange) and theta (blue) oscillations. Two oscillatory cycles are shown. (G) Phase-locking of all tonically active neurons to theta (top) and delta (bottom) oscillation in polar coordinates. Angle, circular phase; length, mean vector length; greyscale corresponds to average firing rate (black, higher firing rate). These neurons did not show strong phase-locking to hippocampal oscillations.

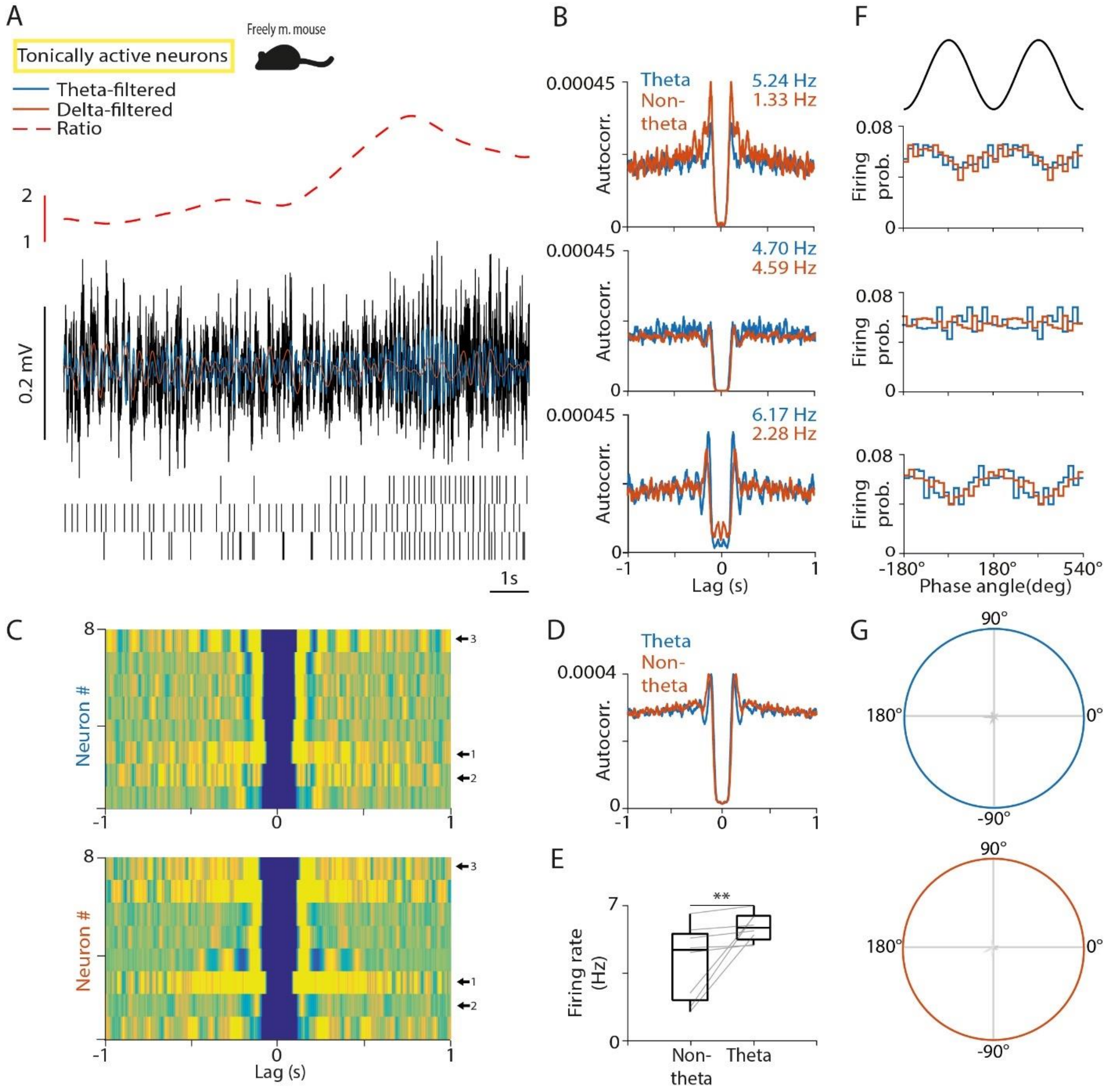


**Figure S15. Tonically active MS neurons in freely moving mice.** (A) Top, black, raw LFP from the CA1 shows a delta-to-theta state transition. Orange, LFP filtered in the delta band; blue, LFP filtered in the theta band; dashed, theta-delta amplitude ratio. Bottom, spike raster of three examples of tonically active neurons from the same recording session. Recordings are from freely moving mice. (B) Autocorrelograms of the three example neurons in panel (A). Numbers indicate average firing rates during theta and non-theta segments. (C) Autocorrelograms of all tonically active neurons during theta (top) and non-theta (bottom) segments. Arrows indicate the example neurons. (D) Average autocorrelogram of all tonically active neurons. (E) Median firing rate of tonically active neurons during none-theta and theta segments. Lines correspond to individual neurons. Firing rate was significantly higher during theta. **, $p < 0.01$,

Wilcoxon signed-rank test. (F) Phase histogram of the example neurons in panel (A) relative to delta (orange) and theta (blue) oscillations. Two oscillatory cycles are shown. (G) Phase-locking of all tonically active neurons to theta (top) and delta (bottom) oscillation in polar coordinates. Angle, circular phase; length, mean vector length; greyscale corresponds to average firing rate (black, higher firing rate). These neurons did not show strong phase-locking to hippocampal oscillations.

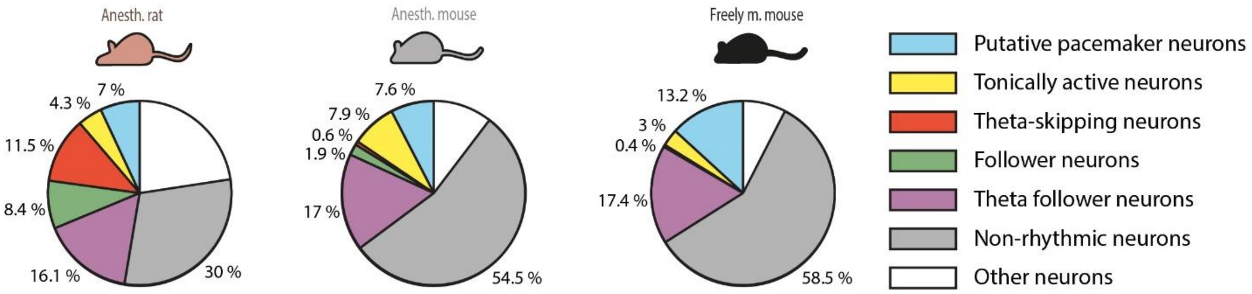


**Figure S16. Rhythmicity groups in the MS.** Proportion of MS neurons in the different rhythmicity groups in anesthetized rats (left), anesthetized mice (middle) and freely moving mice (right).

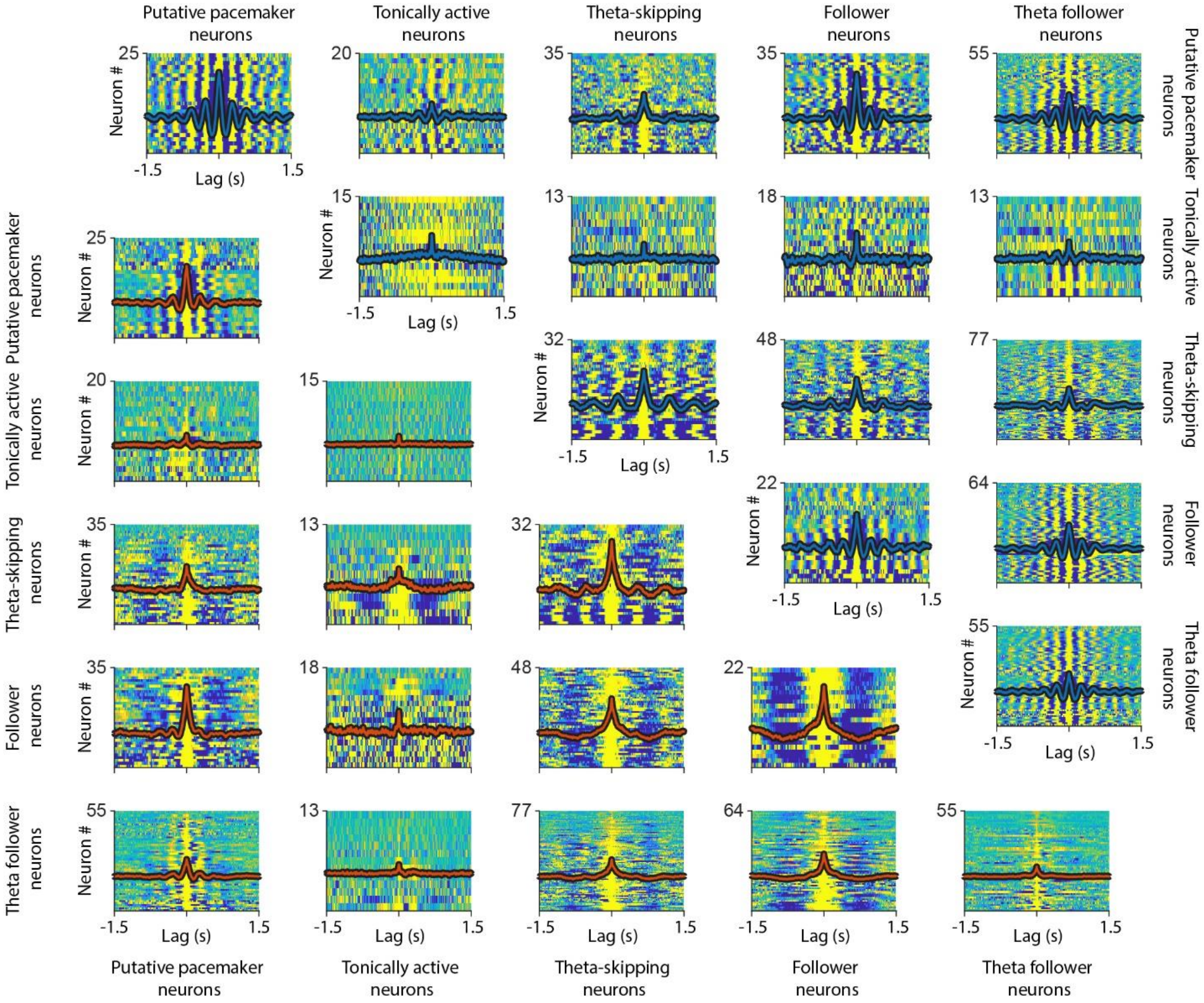


**Figure S17. Crosscorrelation within and between MS rhythmicity groups.** Crosscorrelations of all rhythmic cell pairs recorded in anesthetized rats were sorted by rhythmicity categories. Individual crosscorrelations are shown color coded (small values are blue) and average crosscorrelations are overlaid (red, during non-theta, lower left triangle; blue, during theta, upper right triangle).

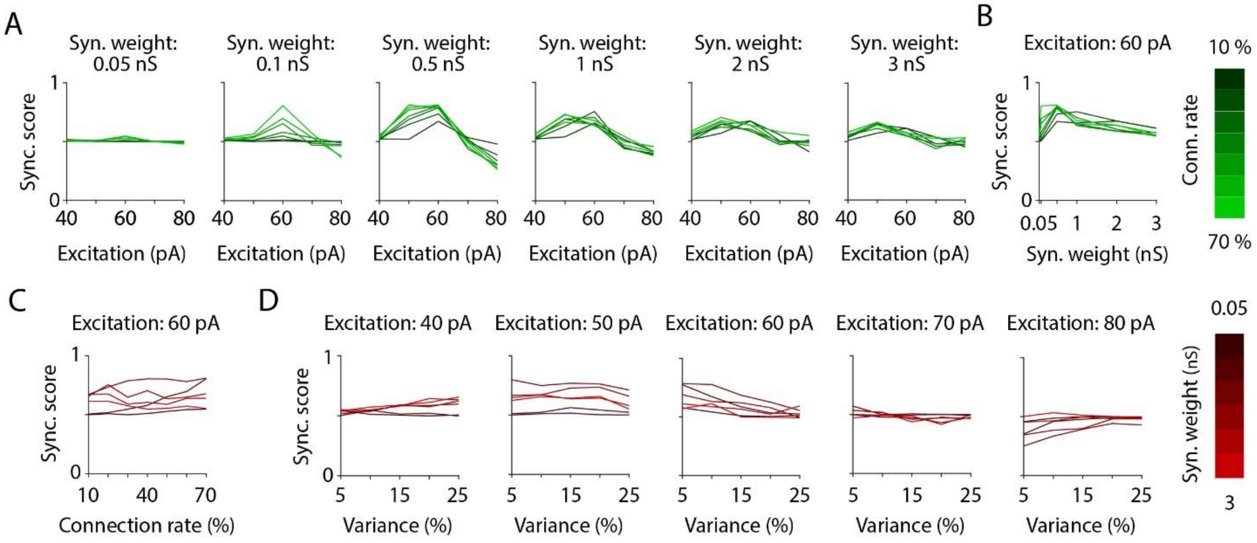


**Figure S18. Parameter-dependence of the synchronization of the network model.** (A) We systematically varied the parameters of the network model to explore the synchronization of the modelled pacemaker circuit with respect to the parameter space. The parameters of the network model were the connection rate (Conn. rate), the mean inhibitory synaptic strength (Syn. weight) and the mean baseline excitation level and its variance. The synchronization score (Sync. score) quantified the proportional time the network spent in the expected network state (non-theta state during baseline excitation and theta state during increased excitation). By systematically varying mean excitation (variance fixed at 10%) and connection rate, we found a discrete peak in synchronization score around 50-60 pA, largely independent of the connection rate. (B) Varying the synaptic weight indicated a peak in the synchronization score at 0.5 nS for connection rates of 30% and above. (C) The model tolerated a range of connection rates. Sparsely connected networks could synchronize only with stronger synapses, while densely connected networks required small inhibitory weights. (D) Next, we systematically varied the variance of tonic excitation strength at a fixed connection rate of 50%. As high variance tended to destabilize the network, reflected in low synchronization scores at 60 pA excitation, we used a moderate 10% variance in the final simulations.

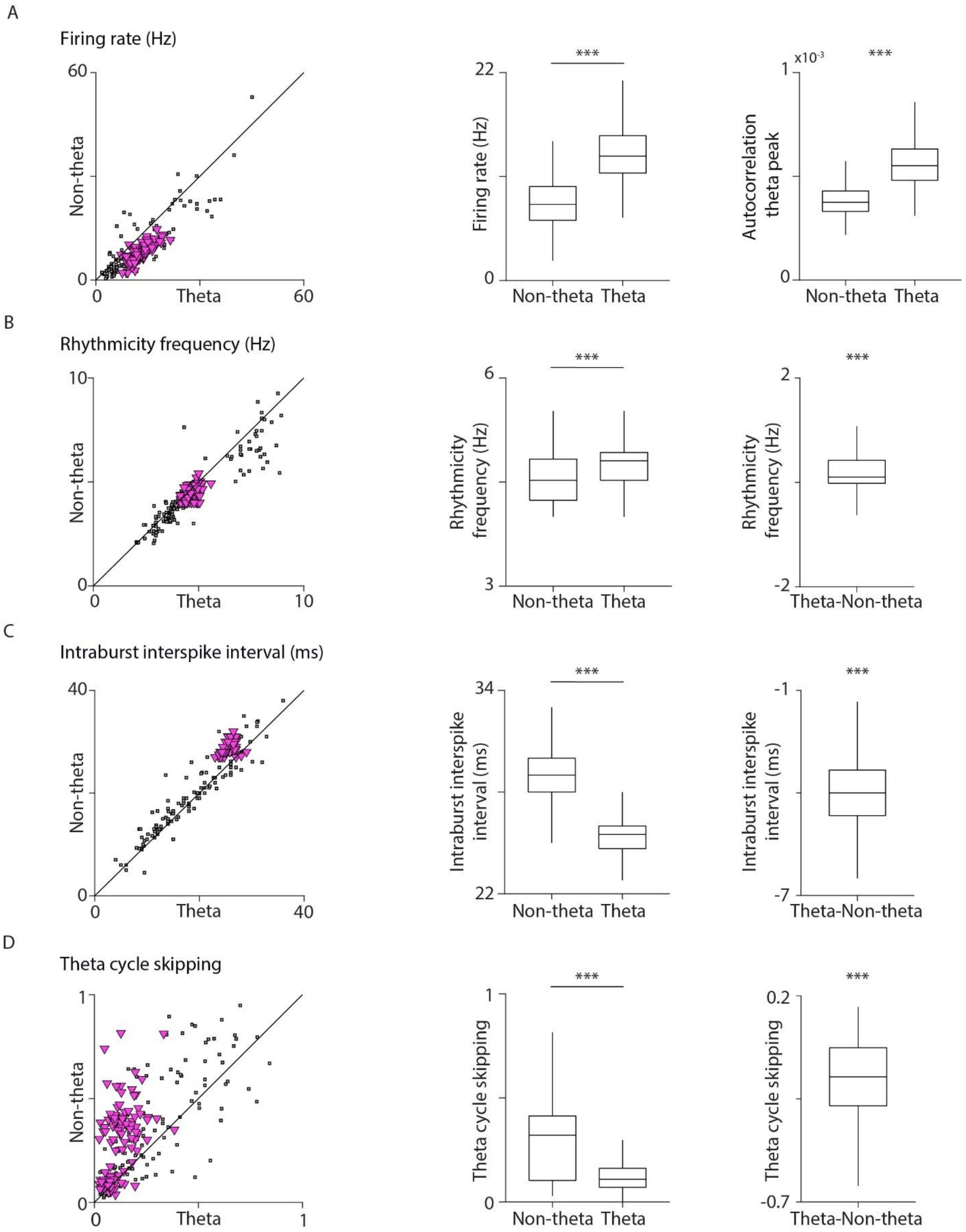
A
Firing rate (Hz)
60
Non-theta
0
0
Theta
60
***
22
Firing rate (Hz)
0
Non-theta
Theta
x10-3
***
1
Autocorrelation
theta peak
0
Non-theta
Theta
B
Rhythmicity frequency (Hz)
10
Non-theta
0
0
Theta
10
***
6
Rhythmicity
frequency (Hz)
3
Non-theta
Theta
***
2
Rhythmicity
frequency (Hz)
-2
Theta-Non-theta
C
Intraburst interspike interval (ms)
40
Non-theta
0
0
Theta
40
***
34
Intraburst interspike
interval (ms)
22
Non-theta
Theta
***
-1
Intraburst interspike
interval (ms)
-7
Theta-Non-theta
D
Theta cycle skipping
1
Non-theta
0
0
Theta
1
***
1
Theta cycle skipping
0
Non-theta
Theta
***
0.2
Theta cycle skipping
-0.7
Theta-Non-theta

**Figure S19. The pacemaker network model reproduces Huygens-synchronization properties of the MS neural circuit.** (A) Firing rates of model pacemaker neurons were higher during theta oscillation than during non-theta segments. Left, scatter plot; real data in black (three data sets pooled, see Figure 4), model data overlaid in magenta (100 randomly selected data points shown in all panels). Middle, box-whisker plot showing model statistics. Right, autocorrelation peak in the theta frequency band for model neurons. ***, $p < 0.001$, Wilcoxon signed-rank test. All box-whisker plots show median, interquartile range and non-outlier range in this figure. (B) Left, scatter plot of rhythmicity frequency measured by the time lag of the first autocorrelation peak in the theta band; real data in black (three data sets pooled, see Figure 4), model data overlaid in magenta. Middle and right, Wilcoxon signed-rank test on model neuron data indicated higher rhythmicity frequency during theta. (C) Left, scatter plot of average intraburst interspike intervals; real data in black (three data sets pooled, see Figure 4), model data overlaid in magenta. Middle and right, the theta-associated decrease of interspike intervals in model neurons indicated a moderate elevation of intraburst frequency. (D) Left, scatter plot of the ratio of skipped theta cycles; real data in black (three data sets pooled, see Figure 4), model data overlaid in magenta. Middle and right, statistics indicated that model pacemaker neurons skipped more theta cycles during non-theta segments.

| Activity during theta | Activity during non-theta | Anesth. rat (#,%) | | Anesth. mouse (#,%) | | Freely m. mouse (#,%) | | |
|---|---|---|---|---|---|---|---|---|
| theta-rhythmic | theta-rhythmic | 29 | 7.0% | 47 | 7.6% | 35 | 13.2% | **Putative pacemakers (theta-bursting)** |
| | | 18 | 4.3% | 49 | 7.9% | 8 | 3.0% | **Tonically active neurons** |
| | delta-rhythmic | 35 | 8.4% | 12 | 1.9% | 0 | 0.0% | **Follower neurons** |
| | non-rhythmic | 67 | 16.1% | 105 | 17.0% | 46 | 17.4% | **Theta follower neurons** |
| delta-rhythmic | theta-rhythmic | 6 | 1.4% | 2 | 0.3% | 1 | 0.4% | **Hypothetical 'inverse' neurons** |
| | delta-rhythmic | 48 | 11.5% | 4 | 0.6% | 1 | 0.4% | **Theta-skipping neurons** |
| | non-rhythmic | 31 | 7.5% | 12 | 1.9% | 5 | 1.9% | |
| non-rhythmic | theta-rhythmic | 21 | 5.0% | 36 | 5.8% | 2 | 0.8% | **Probably part of the pacemaker circuit** |
| | delta-rhythmic | 36 | 8.7% | 14 | 2.3% | 12 | 4.5% | **Delta follower neurons** |
| | non-rhythmic | 125 | 30.0% | 336 | 54.5% | 155 | 58.5% | **Non-rhythmic neurons** |
| **Total** | | **416** | **100.0%** | **617** | **100.0%** | **265** | **100.0%** | |

**Supplemental Table 1. Number of neurons in different rhythmicity groups in the MS.** Theta-rhythmic neurons were separated to theta-bursting MS neurons reported in previous studies and tonically active neurons that show rhythmic firing in the theta band but do not exhibit rhythmic bursting properties and show little synchrony with ongoing theta activity. Follower neurons reflected theta versus non-theta state of the network. For part of neurons, this type of activity was only confirmed statistically for either of these states (theta followers and delta followers). Neurons that were delta-rhythmic during theta oscillation were identified as theta-skipping neurons. Figure 3 focuses on those cells that were delta-rhythmic during both states. Neurons that were theta-rhythmic during non-theta segments but found non-rhythmic during theta segments may be part of the pacemaker group, where the amount of data (number of spikes and length of identified theta segments) may have been insufficient for statistical detection of rhythmicity during theta. It is theoretically possible that some MS neurons would be theta-rhythmic in non-theta state but delta-rhythmic in theta state. However, this group was negligible in all three data sets.

| | Anesth. rat | Anesth. mouse | Freely m. mouse | Pooled | Figure | Model | Figure |
|---|---|---|---|---|---|---|---|
| **Firing rate (Hz)** | $1.04 \times 10^{-4}$ | $7.98 \times 10^{-6}$ | $1.34 \times 10^{-2}$ | **$1.40 \times 10^{-9}$** | Fig. 4A | $9.42 \times 10^{-149}$ | Fig. S19A |
| **Autocorrelation theta peak** | $4.58 \times 10^{-5}$ | $1.88 \times 10^{-2}$ | $6.17 \times 10^{-6}$ | **$6.83 \times 10^{-10}$** | Fig. 4A | $3.68 \times 10^{-133}$ | Fig. S19A |
| **Rhythmicity frequency (Hz)** | $1.37 \times 10^{-3}$ | $1.53 \times 10^{-1}$ | $2.21 \times 10^{-5}$ | **$2.64 \times 10^{-7}$** | Fig. 4B | $8.97 \times 10^{-63}$ | Fig. S19B |
| **Intraburst interspike interval (ms)** | $2.95 \times 10^{-3}$ | $3.30 \times 10^{-1}$ | $1.95 \times 10^{-4}$ | **$7.34 \times 10^{-5}$** | Fig. 4C | $9.41 \times 10^{-145}$ | Fig. S19C |
| **Theta cycle skipping** | $2.06 \times 10^{-3}$ | $4.41 \times 10^{-5}$ | $4.61 \times 10^{-1}$ | **$3.02 \times 10^{-6}$** | Fig. 4D | $2.58 \times 10^{-119}$ | Fig. S19D |
| **Relative rhythmicity frequency difference** | $4.53 \times 10^{-3}$ | $5.61 \times 10^{-2}$ | $1.48 \times 10^{-5}$ | **$3.17 \times 10^{-6}$** | Fig. 4E | $2.35 \times 10^{-97}$ | Fig. 5F |

**Supplemental Table 2. Statistics for synchronization analysis in individual data sets.** Since all three data sets showed similar results, Figure 4 displays pooled results. This table provides p-values (two-sided Mann-Whitney U-test for unpaired samples and two-sided Wilcoxon signed-rank test for paired samples) for the three data sets separately and pooled. Corresponding statistical tests on simulated data are also indicated.
